# A DNA Typewriter records the cell lineage history of a mouse, from zygote to late organogenesis

**DOI:** 10.64898/2026.07.29.741625

**Authors:** Qi Yu, Haedong Kim, Sophie Seidel, James F. Acosta-Clark, Beth K. Martin, Kyle O’Connor, Riza M. Daza, Molly Gasperini, Jenny F. Nathans, Maggie Lam, Elena Gamo, Shruthi VijayKumar, Lauren Kuo, Jean-Benoît Lalanne, Kamen P. Simeonov, Marion Pepper, Cole Trapnell, Jesse M. Gray, Junhong Choi, Chengxiang Qiu, Jay Shendure

## Abstract

Mammalian biology unfolds over time, within tissues and organs opaque to our eyes and instruments. We applied DNA Typewriter, a sequential molecular recorder, to record the cell lineage of a mouse over nearly two weeks of development. From one embryo, we reconstruct a time-calibrated, parsimony-supported, zygote-rooted phylogeny of 1.28M transcriptionally-profiled cells. A burst of editing unequivocally marks the daughters of the first cleavage, which serve as inline replicates. We quantify clonal dominance arising during gastrulation. Tree siblings share cell type far above chance; heterotypic siblings mark terminal differentiations. Temporal sweeps of clade co-occurrence recover a dated hierarchy of cell-type couplings; imputed labels for internal nodes recapitulate known state paths. A lineage-anchored ontogeny of mammalian development, long out of reach, is coming into view.

## Main Text

A half-century ago, Sulston and colleagues watched *C. elegans* develop under the microscope, painstakingly drawing each cell division until they had mapped its complete lineage, zygote to adult (*1*, *2*). It remains one of the singular accomplishments of modern biology, and no comparable lineage has been mapped for any other animal since. But *C. elegans* is exceptional: transparent, small and developmentally invariant.

Nearly all other animals are opaque, and many vastly larger, *e.g.* an adult mouse contains billions, and an adult human trillions, of cells (*3*). They also develop in the opposite, regulative mode. Cell fates are shaped by extrinsic cues rather than lineage alone, and the precise sequence of divisions varies between individuals. Lineage is nonetheless far from irrelevant, and the outstanding mystery of regulative development is how a variable lineage is canalized into reproducible form and function.

To study mouse development at the whole-organism scale with genomic tools, we and others have performed molecular profiling of cells sampled from temporally staged individuals (*4–9*). However, sampling is destructive, severing each cell from its past and future. Although a cell type ontogeny can be inferred in broad strokes from such time-series, lineage relationships cannot (*10*).

That the genome is a digital medium, faithfully copied at each cell division, makes it natural to exploit for recording cell lineage. Many studies have leveraged natural somatic variation, static barcodes or recombinase-diversified barcodes to this end (*11–18*), but these strategies have not delivered dense cellular phylogenies (*19*, *20*).

Around 2016, we and others began to pursue a paradigm wherein genome editors introduce heritable marks that accrue throughout development (*19–25*). Read at an endpoint by single-cell profiling of endogenous transcripts and barcodes (*25–28*), a single destructive snapshot informs cells’ present states and their relationship to other cells. Implementations group by editor class: Cas9 nucleases, whose error-prone repair leaves heritable scars (*e.g.* GESTALT, MEMOIR) (*21–28*); base editors, which install point mutations (*e.g.* SMALT, baseMEMOIR) (*29–32*); and prime editors, which write symbolic insertions (*e.g.* DNA Typewriter, peCHYRON) (*33–35*).

Despite progress, dense cellular phylogenies of large animals have remained out of reach for a confluence of technical reasons (*19*), including: (i) <u>delivery and expression</u>: all components must reach the zygote and be expressed stably throughout development; (ii) <u>information loss</u>: double-strand breaks at densely packed sites trigger deletions that destroy adjacent records and media; (iii) <u>instability of temporal resolution</u>: as editable sites are consumed, the recording rate naturally falls; (iv) <u>event ordering</u>: if edits are unordered, their temporal order must be inferred; (v) <u>scope</u>: lineage histories alone don’t reveal the historical signals and states that shaped cells’ fates; (vi) <u>cellular coverage</u>: dense reconstructions require profiling a substantial proportion of an organism’s cells; (vii) <u>records retrieval</u>: recovering barcodes from profiled cells/nuclei is technically demanding; (viii) <u>tree building</u>: algorithms developed for 10^2^-10^4^ taxa must scale to 10^5^-10^9^ cells/nuclei; and (ix) <u>design-test cycle time</u>: optimizing recorders (or anything) *in vivo* is slow.

DNA Typewriter (*33*) addresses, partly or wholly, four of these hurdles (**Fig. 1A**). Prime editing does not require double-strand breaks (ii). Each edit consumes a monomer and activates a new one, stabilizing the number of editable monomers (iii). Insertional edits are introduced 5′→3′, so their order is preserved (iv). And specific insertions can be rendered biologically conditional, extending what can be recorded beyond lineage (v) (*36*). To date, however, we have demonstrated DNA Typewriter only *in vitro*, in cultured cells and monoclonal gastruloids, to build phylogenies of up to ∼50,000 cells (*33*, *37*).

**Figure 1.**
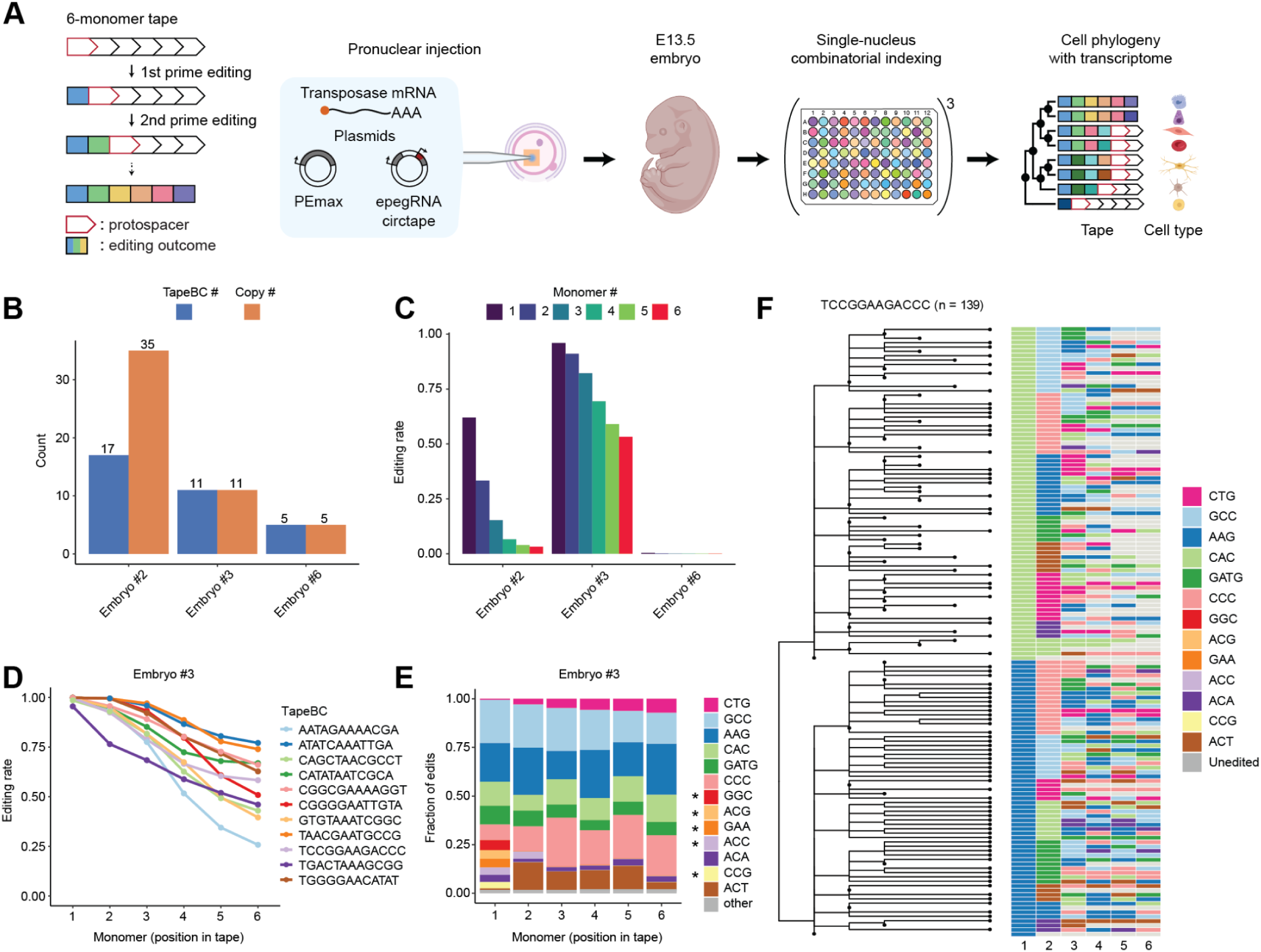
Pronuclear zygotic injection (PNI) of DNA Typewriter components identifies an E13.5 embryo with robust levels of sequential editing. **(A)** Schematic. Left: a 6-monomer tape is edited sequentially by prime editing, with each event installing an insertional symbol (colored) at the active monomer while regenerating the protospacer (red outline) at the next. Middle: constructs encoding prime editor (PEmax), guides (epegRNAs), and barcoded tape (tape-BC, circtape), were delivered to the zygote by PNI, together with piggyBac transposase mRNA. The tape is transcribed by Pol-III and circularized via the Tornado system, and additionally bears a 30-nt poly(A) region and a custom capture sequence, enabling oligo-dT-based and (optionally) targeted reverse transcription, respectively. Right: embryos were harvested at E13.5 and transcriptionally profiled by three-level single-nucleus combinatorial indexing (sci-RNA-seq3) (*39*), yielding paired circtape genotypes and transcriptomes that together facilitate the reconstruction of a cell phylogeny annotated by cell type. Some elements were created in BioRender (*86*). **(B)** Number of distinct tape-BC sequences (unique integration barcodes; blue) and the estimated total number of genomic integrations after accounting for multi-copy barcodes (orange). **(C)** Per-monomer editing rates for each embryo, expressed as the fraction of resolved reads edited at each of the six monomers (colored 1-6), collapsed with respect to independent integrations. **(D)** Per-monomer editing rate in embryo #3, shown separately for each of 11 single-copy tape-BC integrations. Editing declines progressively from monomer-1 to monomer-6, consistent with sequential editing. **(E)** Frequency of each insertional symbol at each of the six monomers in embryo #3, aggregated across integrations. The 13 symbols present at ≥0.5% frequency at one or more monomer positions are shown individually; the remainder are grouped as “other”. The 5 symbols appearing at ≥0.5% frequency only at monomer-1 and/or monomer-2 are marked with an asterisk. **(F)** Neighbor-joining lineage tree reconstructed from tape genotypes observed at a single representative integration (tape-BC “TCCGGAAGACCC”; n = 139 genotypes), with the corresponding edit patterns shown at right (colored by symbol; grey, unedited).

Here, contending with the remaining obstacles and several more that arose along the way, we apply DNA Typewriter *in vivo* to reconstruct a 1.28M-cell phylogeny of a single mouse embryo, from zygote to late organogenesis.

## Results

### Pronuclear injection of DNA Typewriter yields an extensively edited E13.5 embryo

DNA Typewriter comprises a prime editor, guide RNAs and tape (here, PEmax, epegRNAs, TAPE-1) (*38*). TAPE-1 is composed of six tandem repeats of a 14-bp monomer preceded by a 3-bp key (here, GGA). Initially, the 5’-most portion of the tape comprises an active 20-bp protospacer (“typehead”). “Keystrokes” insert an information-bearing “symbol” (here, NNNGGA) that disrupts the typehead while also activating a new, downstream typehead (**fig. S1A**). Exactly one monomer per tape is editable until the array is exhausted.

We introduced DNA Typewriter by pronuclear injection (PNI) of two constructs and piggyBac transposase-encoding mRNA into zygotes. One construct encoded constitutively expressed PEmax, and the other an NNNGGA-encoding epegRNA, 12-bp barcode (tape-BC) and 6-monomer tape (**Fig. 1A**; **fig. S1A**). Differentiated tissues preclude unbiased cellular dissociation of post-gastrulation mouse embryos, which we circumvent by profiling nuclei rather than cells (*39*). But mRNA is scarcer in nuclei, and targeted transcript recovery in single-nucleus RNA-seq workflows has been a persistent challenge (*37*, *40*). Because circular RNA reporters are recovered better than their linear equivalents (*41*), we reengineered TAPE-1 for expression as a Pol-III-driven circular transcript (“circtape”; **fig. S1A**). Following PNI, we isolated nuclei from ten embryos harvested at E13.5 (**fig. S1B**). Extraembryonic tissues were discarded. By ddPCR and bulk amplicon sequencing of tape-BCs, three bore epegRNA::tape-BC::circtape integrations, and DNA Typewriter activity at their tapes tracked PEmax copy number (**Fig. 1B-C**; **fig. S1C-D**).

We focused on embryo #3 (**Fig. 1D**). With 11 epegRNA::tape-BC::circtape integrations, we expected a “vocabulary” of at most 11 symbols; we instead observed 13 at ≥0.5% frequency, including 5 abundant at monomers 1-2 and 8 at monomers 2-6 (**Fig. 1E**; **fig. S2A**). Amplicon sequencing linking tape-BCs to their *cis*-encoded symbols recovered 10 of 11, some redundant and collectively encoding exactly the 8 symbols seen at monomers 2-6. Symbols were distributed independently of whether they were encoded *in cis* (**fig. S2B**). A parsimonious explanation of these patterns is that the 5 “extra” symbols abundant only at monomers 1-2 were written by transiently expressed epegRNAs shortly after PNI.

With 8 symbols at 6 monomers, 8^6^ (∼10^5^) “tape genotypes” (an ordered set of symbols at a given tape) are possible. To assess how much of this diversity was realized, we performed rarefaction on tape genotypes (**fig. S2C**) (*42*). Each integration was represented by an estimated 180-370 genome equivalents distributed across 76-165 distinct tape genotypes, and every accumulation curve remained steeply rising. Recorded diversity thus appears limited by input material rather than by recorder capacity.

We reconstructed neighbor-joining trees from tape genotypes with a DNA Typewriter-specific distance metric (**Fig. 1F**; **fig. S3**) (*43*, *44*). For 9 integrations, >95% of tape genotypes bore only one of two symbols at monomer-1, *e.g.* at integration “TCCGGAAGACCC”, all 139 tape genotypes bore CAC (55%) or AAG (45%). A parsimonious explanation is that a burst of prime editing occurred at the 2-cell stage.

To ask whether these patterns are reproducible, we generated four additional DNA Typewriter embryos and characterized their tapes in bulk (**Supplementary Text, note S1**). Each was comparable or superior to embryo #3 in integration number, editing rate and recorded diversity (**fig. S4**). Across 85 integrations in 5 well-edited embryos, edits inferred to arise at the 1-cell and 2-cell stages were common, while 4-cell-stage edits were uncommon, with the caveat that there are limits on how readily the latter can be discerned from bulk tape genotypes. Notably, only embryo #3 lacked 1-cell-stage edits.

We conclude that PNI-delivered DNA Typewriter records heritable, ordered, diverse marks *in vivo*. Because each tape is edited independently, their combinatorics expands the upper bound on potential configurations from 8^6^ per integration to 8^(6*11)^ (∼10^59^) per cell. However, realizing this potential requires determining which edited tapes co-occur within which cells.

### Single-nucleus RNA-seq of a DNA Typewriter mouse embryo

We subjected ∼10M nuclei isolated from embryo #3 to single-nucleus combinatorial indexing (sci-RNA-seq3) (*39*), modified to co-amplify transcriptome and circtape libraries (**Fig. 1A**). Following quality control, we obtained transcriptional profiles for 1,584,848 nuclei. To annotate cell types, we co-embedded these profiles with 1.65M E12.75-E14.25 cells from our 11.4M-cell atlas of mouse development (*5*), and transferred labels by k-nearest neighbors; cell type and major trajectory labels were transferred separately (**Fig. 2A**). Embryo #3’s cell type proportions were maximally concordant with E13.5, the intended harvest stage (**Fig. 2B**). Stress, apoptosis, interferon and DNA-damage modules were unelevated in embryo #3 across 26 abundant cell types (**fig. S5A-B**). However, of 91 genes in the vicinity of a mapped integration site (**table S1**), 8 were differentially expressed, most prominently the chromatin remodeler *Chd3*, which exhibited reduced expression across 20 cell types (**fig. S5C**).

**Figure 2.**
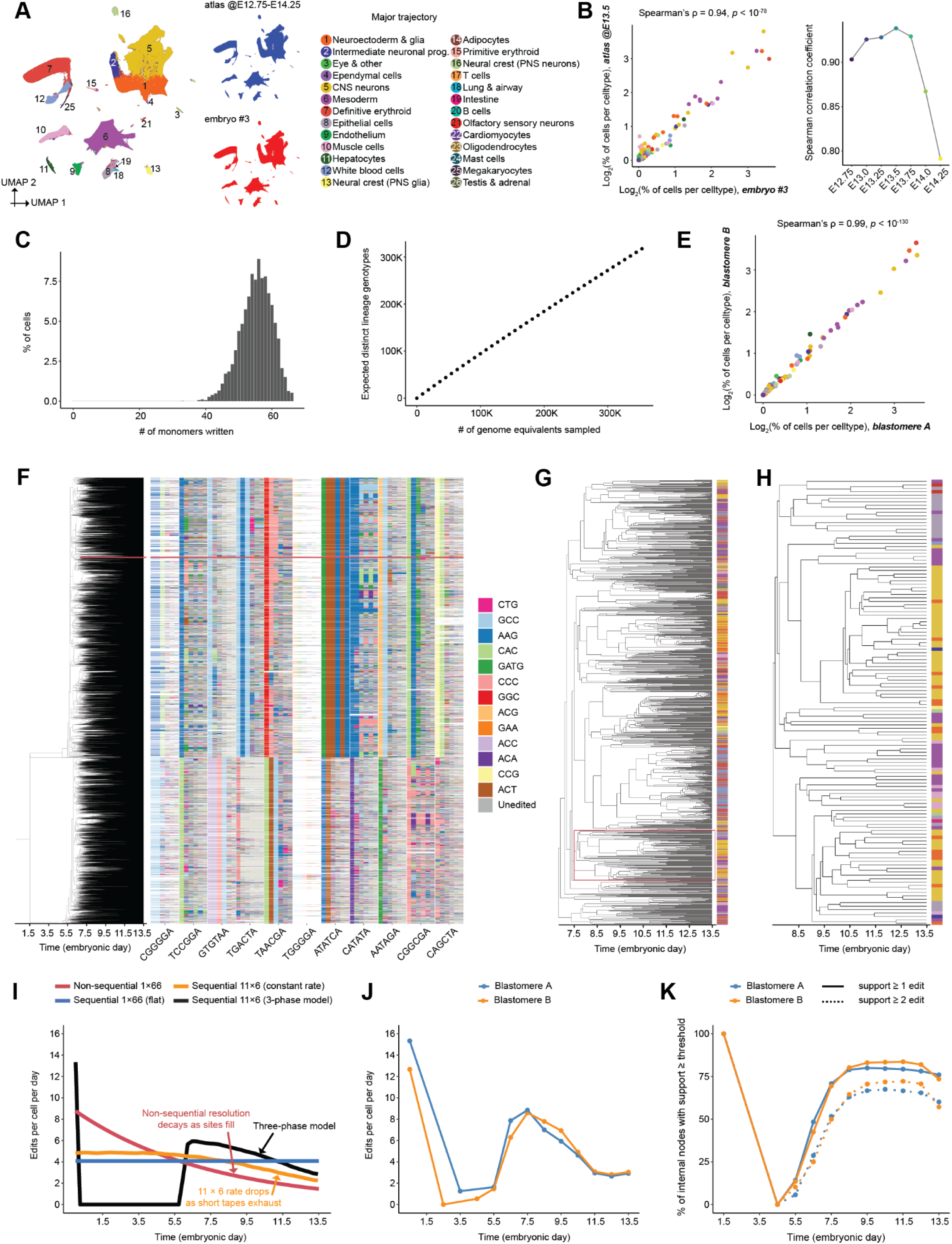
Single-nucleus RNA-seq of a DNA Typewriter embryo enables reconstruction of a time-calibrated, zygote-rooted, tip-annotated phylogeny. **(A)** 2D UMAP co-embedding of 1.58M scRNA-seq profiles from embryo #3 (this study; median UMI: 906, genes: 654) and 1.65M from 7 timepoints of our high-temporal-resolution single-cell atlas of mouse development (E12.75 - E14.25 in 6-hr increments) (*5*). Left: all cells, colored by the 26 major trajectories (numbered key); major trajectories accounting for more than 0.1% of cells are labeled. Right: the same embedding split by dataset of origin. **(B)** Left: log_2_-scaled percentage of cells per cell type, for each of the 169 cell types detected among cells shown in panel **A**. Each point is one cell type, colored by its major trajectory. Right: Spearman’s ρ between cell type proportions in embryo #3 vs. each wildtype atlas timepoint from E12.75 to E14.25. **(C)** Histogram of the number of edited monomers per cell in embryo #3, for the 4,371 cells with tape genotypes at all 11 tape integrations. Although cells genotyped at all 11 tapes are enriched for cell types with higher tape recovery, reweighting this distribution to the embryo’s major-trajectory composition shifts it only from 55.2 ± 4.8 to 54.9 ± 5.1 monomers written (mean ± s.d.). **(D)** Cell-level rarefaction analysis of lineage-genotype diversity (Hurlbert interpolation), based on the 353,681 cells with a lineage genotype call for all 5 of the most highly recovered integrations. The curve remains near-linear. **(E)** Log_2_-scaled percentages of cells per cell type in blastomere A versus blastomere B, for the 141 cell types detected in the union of A and B. Each point is one cell type, colored by its major trajectory. **(F)** Timed backbone phylogeny (root fixed E0; A and B subtrees fixed at E1.5, tips fixed at E13.5). The tree (left) is shown beside the per-cell tapes, colored by the observed edit outcome (right). **(G-H)** Zoom-in on clades outlined by red rectangles in panels **F** (1,167 cells; clade root at E6.8) and **G** (130 cells; clade root at E7.5), respectively. Major trajectory annotations of tips are shown at right (color legend in panel **A**). **(I)** Mean edits accumulated per cell versus developmental time, for four models, each tuned to reach 55 of 66 edits by E13.5. Non-sequential editing to a 1 × 66-monomer tape under a constant per-site, per-day rate (red) yields ∼6-fold variation in temporal resolution, from ∼8.8 edits/cell/day at the outset to ∼1.5 by E13.5, as sites are consumed. Sequential editing to a 1 × 66-monomer tape under a constant per-site, per-day rate (blue) holds flat at ∼4.1 edits/cell/day. Sequential editing to 11 × 6-monomer tapes under a constant per-site, per-day rate (orange) behaves similarly, except that the aggregate rate falls late as individual tapes reach their limit. Sequential editing to 11 × 6-monomer tapes under a three-phase model (black) comprises a 20-edit burst, a slowdown interval approximated as quiescence until E6, and then a constant per-site, per-day rate. **(J)** Observed tape editing rate across developmental time for blastomere A and blastomere B. On the backbone tree, ancestral tape states were inferred as the longest edit prefix shared by all descendant genotypes of each node (a conservative rule at nodes with many descendants), and branches assigned to 1-day bins from E0 to E14 by their temporal midpoint; the rate for each bin is the sum of edits across integrations divided by the total branch time falling in that bin. The peak in the E0-E1.0 bin corresponds to the founder burst, placed on each blastomere’s stem lineage. **(K)** Internal node support across developmental time for blastomere A and B, binned by node date over the same 1-day windows. Support is the number of independent edits inferred to occur along the branch entering a node, from the same ancestral-state reconstruction as panel **J**; panels show the percentage of nodes with support ≥1 or ≥2. In panels **J-K**, points are plotted at bin midpoints, and bins containing no branch midpoint (panel **J**) or dated node (panel **K**) are omitted rather than plotted as zero (**J**: E1.5 in both blastomeres, E2.5, E4.5 in A, E3.5 in B; **K**: E2.5, 3.5 in both blastomeres); panel **K** additionally begins at E1.5, the age of the blastomere subtree roots. These gaps, and the sparse sampling of neighboring early bins, reflect the small number of resolvable divisions during the slowdown interval.

Summed across all integrations, we obtained a median of 51 quality-passing circtape UMIs (unique molecular identifiers) per cell. Tape genotypes were called for 54% of cell × tape combinations among genotyped cells (49% to 70% for nine integrations; 22% and 4% for the remaining two). Circtape and transcriptome recovery were correlated, with limited evidence for cell-type-specific silencing of individual tapes (**fig. S6**). 95% of tape genotypes bore only expected prime editing events; the remaining 5% included nine classes of unexpected events (**Supplementary Text, note S2**).

Recording capacity was heavily utilized. Among the 4,371 cells genotyped at all 11 tapes, 55 ± 5 monomers had been written (83%), unimodally distributed and peaking at 56, with only 23 cells (0.5%) bearing 66 recovered edits (**Fig. 2C**). Repeating the estimate across the entire matrix by summing the per-tape means, we obtain a similar estimate of 53/66 monomers. A caveat is that although TAPE-1 was designed with six monomers, array length by E13.5 exhibits substantial variation consistent with *in vivo* slippage within the tandem array (5.2 ± 1.2 monomers; **Supplementary Text**, **note S2**), such that our estimate of utilization is a lower bound.

We define a cell’s “lineage genotype” as its constellation of tape genotype calls. Among 353,681 cells genotyped at the five best recovered circtapes, only 15% shared an identical lineage genotype with another cell, and the accumulation curve was again linear (**Fig. 2D**). Resolution is thus limited by the number of cells sampled rather than by recording activity. Reassuringly, of 11,460 lineage genotypes shared by exactly two cells, 57% were of the same cell type, 11-fold above chance, an early indication that lineage genotypes capture biologically meaningful structure.

### Recording dynamics shape dating and branch support of a 655,701-cell backbone phylogeny

The burst of prime editing activity at the two-cell stage allows us to resolve their descendants directly. From 401,367 cells with ≥8 genotyped tapes, we derived the edit state of each blastomere, hereafter “A” and “B” (A: 23 edits, B: 19 edits; **table S2**) and routed cells to their blastomere-of-origin. Of 1,584,848 transcriptionally profiled nuclei, 25,429 yielded no genotyped tape. Of 1,559,419 remaining cells, 889,849 (57.1%) routed to A and 653,335 (41.9%) to B, 1,543,184 (99.0%) in total; the remaining 16,235 (1.0%) lacked an unambiguous founder assignment and were set aside.

The cell-type compositions of A and B were highly correlated (Spearman’s ρ = 0.99; **Fig. 2E**), suggesting contributions to the embryo proper, although unequal, are fate-neutral. However, our data do not speak to the relative or compositional contributions of A and B to extraembryonic lineages, not sampled here (*45*, *46*).

We reconstructed phylogenies for A and B independently, and hereafter treat them as inline replicates. Because >10^6^ taxa exceed the practical limits of direct phylogenetic reconstruction, we proceeded in two stages: (i) tree-building and dating of backbone trees from high-tape-recovery cells; then (ii) distance-based placement of moderate-tape-recovery cells onto the backbones.

Backbone trees were built by scalable neighbor-joining (*47–49*) on distance matrices for 410,925 (A) and 244,776 (B) cells with ≥7 genotyped tapes (*43*), rooted on the founders’ states. We dated internal nodes by least-squares dating under a strict molecular clock (*50*, *51*), fixing root (E1.5) and tips (E13.5), imposing a 2-hour minimum branch length, and capping the number of ancestral lineages for each blastomere’s tree at half of the embryo’s estimated cell number, where available (**table S3**). Joining the dated trees at a common zygote root yielded a backbone phylogeny of 655,701 tips (**Fig. 2F-H**).

The floor and ceiling are principled, the former set by the fastest cell cycles of the gastrulating embryo (*52*) and the latter by the fact that a subsample can contain no more lineages at any given time than the embryo had cells. In practice, the ceiling binds only near gastrulation, where ceiling-free dating places 97,566 lineages at E6.5 against an epiblast of ∼660 cells (**fig. S7A**), yet costs only ∼1.8% in the least-squares objective, shifting early nodes ∼4 days later, then decaying monotonically toward the tips (**fig. S7B**). Although we do not know when the 1.36-fold excess of A vs. B cells arising by E13.5 accrues, node dates are robust to the ratio at which the ceiling is applied (**fig. S7C**). However, that such a correction is needed at all localizes the clock’s violation to the interval between the 2-cell stage and gastrulation, where the ceiling substitutes external knowledge of embryo size for a temporal signal we could not recover for this interval, due to limitations of the recorder, the reconstruction, or some combination thereof.

In theory, DNA Typewriter’s sequential design should hold the number of active recording monomers, and thus the rate at which independent marks accumulate, constant over time. To ask whether it did, we compared the observed accumulation against three idealized architectures, each tuned to reach the observed mean of 55/66 monomers written by E13.5 (**Fig. 2I**). Observed dynamics matched none of them, instead showing three phases (**Fig. 2J**): a ∼20 edit burst at the 2-cell stage, marking each founding blastomere; a pronounced slowdown through subsequent cleavages and blastocyst formation; and recovery around ∼E6, peaking at ∼9 edits/cell/day then declining as tapes exhaust. A minimal three-phase model reproduces these dynamics (**Fig. 2I**). Furthermore, although we lack single-cell resolution for the four additional embryos, their bulk editing patterns are consistent with similar dynamics (**Supplementary Text**, **note S1**). A caveat is that, because our parsimony approach assigns an edit to the ancestral state of a node only when all of its descendants share that edit, noise in the genotype calls can bias the reconstruction towards assigning too few edits, with earlier nodes, which have far more descendants, more likely to be affected; therefore, the magnitude of this slowdown should be read as an upper bound.

How reliable is the topology of this backbone phylogeny? Node support measures are a cornerstone of phylogenetics (*53*), but their use in the lineage recording field is inconsistent. Resampling-based bootstraps (*53*), a conventional measure of node support, were computationally infeasible at this scale. We therefore devised a simple independent-edit-count support statistic for each split in the inferred phylogeny (**Methods**). Its distribution was right-skewed (median 3, tail beyond 20; **fig. S7D**), with 78% (A) and 80% (B) of internal branches carrying ≥1, and 64% and 68% carrying ≥2, supporting edits. As further checks, we re-counted support using only edits on distinct tapes (**fig. S8A**), and confirmed that subtrees reconstructed with one tape withheld recover the held-out tape’s pairwise distances well above a label-shuffled control (**fig. S8B**).

Recording dynamics and topological confidence are necessarily related, both because an interval in which few marks are written is one in which few divisions can be resolved, and because both are estimated from the same ancestral-state reconstruction. Correspondingly, node support varied with developmental time, mirroring editing dynamics (**Fig. 2K; fig. S8A**): maximal at the first division, weakest through the more sparsely recorded window that follows, then rising as editing recovers, before falling again slightly toward the tips as tape exhaustion reduces the marks available to resolve recent splits. Notably, the low-confidence early window is precisely where unconstrained dating inflates ancestral lineages beyond the embryo’s cell number, and where the imposition of a ceiling does the most work (**fig. S7A-B**). Support recovers by E6.5-E7.5, over which the dated tree traces an expansion from 656 to 13,181 lineages, and is maintained thereafter.

A third clock violation, addressed by neither the ceiling nor the timing correction, is that prime editing rate varies across cell types (*54–57*). In this embryo, cell type-stratified editing rates fall within a tight 1.6-fold range, but the slowest types are overwhelmingly hematopoietic (**Supplementary Text, note S3**; **fig. S9**). Our dates should therefore be read as approximations rather than exact ages, least reliable at the earliest divisions, underestimated toward the tips, and least reliable again in hematopoietic cell types.

### Distance-based placement doubles the tree to 1,281,141 cells

In the second reconstruction stage, we placed 625,440 additional cells with 4 to 6 genotyped tapes onto the 655,701-cell backbone; the remaining 262,043 blastomere-routed cells were excluded, 88% for carrying <4 genotyped tapes and the rest by the doublet and founder-contradiction filters. For placements, each query cell was attached to its single closest relative under the same distance metric used for the backbone (*43*), whether a backbone tip or a previously placed query; cells were processed in order of decreasing tape recovery (**fig. S10A**). This strategy rests on two assumptions: that the backbone samples the embryo densely enough for each query cell to have a near relative among its tips, and that 4 tapes suffice to identify that relative. Neither is guaranteed, so we assessed our placements in two orthogonal ways.

First, the edit distances themselves. Query cells share post-burst, sequentially accrued edits with their anchor and rarely contradict it (**fig. S10B-C**), indicating that anchors are genuine close relatives rather than arbitrary attachment points. But most queries are genotype-identical to their anchor at every recovered monomer, so placements add little topological information and imply short divergence times (**fig. S10D-E**).

Second, because placement uses edit distances alone, with no reference to transcriptional state, cell type concordance among neighboring tips serves as independent evidence that genuine lineage relationships are recovered. Because inserting placements reshapes the terminal topology, adding polytomies where queries share identical genotypes (**fig. S10F-G**), we compared cells on the fraction of their equally related nearest neighbors sharing their cell type. Backbone- and placement-derived cells were similarly concordant, both far above a label-permuted null, with placed cells somewhat lower in every terminal context (**fig. S10H**).

Placed cells therefore sit at short edit distances from their anchors and recover cell-type concordance nearly as well as backbone cells. Furthermore, they are compositionally similar to backbone cells (**fig. S11**). On the other hand, their accuracy is sensitive to both retained tape number and edit count (**fig. S12**). Placements should thus be cautiously interpreted as being in the correct local neighborhood, but not necessarily at the correct depth, a limitation that follows from the algorithm attaching queries along terminal branches rather than internal edges in order to remain computationally tractable.

The final tree of embryo #3 is ultrametric, with 1,281,141 tips (E13.5), each carrying a single-cell transcriptome and one of 138 cell type annotations (**Fig. 3A**). Although these cells represent only ∼10% of the E13.5 embryo (*4*, *5*, *58*), coverage at depth is far higher, *e.g.* each of the 13,181 lineages dated to E7.5 is represented by a mean of 97 sampled descendants. Inspired by NextStrain (*59*), we built NextCell (*60*), an interactive explorer with which one can click any ancestral node to visualize its subclade. For example, **Fig. 3B** shows n1703121, an ancestral node dated to E7.4 whose descendants are endodermally fated, with subclades clonally partitioned among visceral organs.

**Figure 3.**
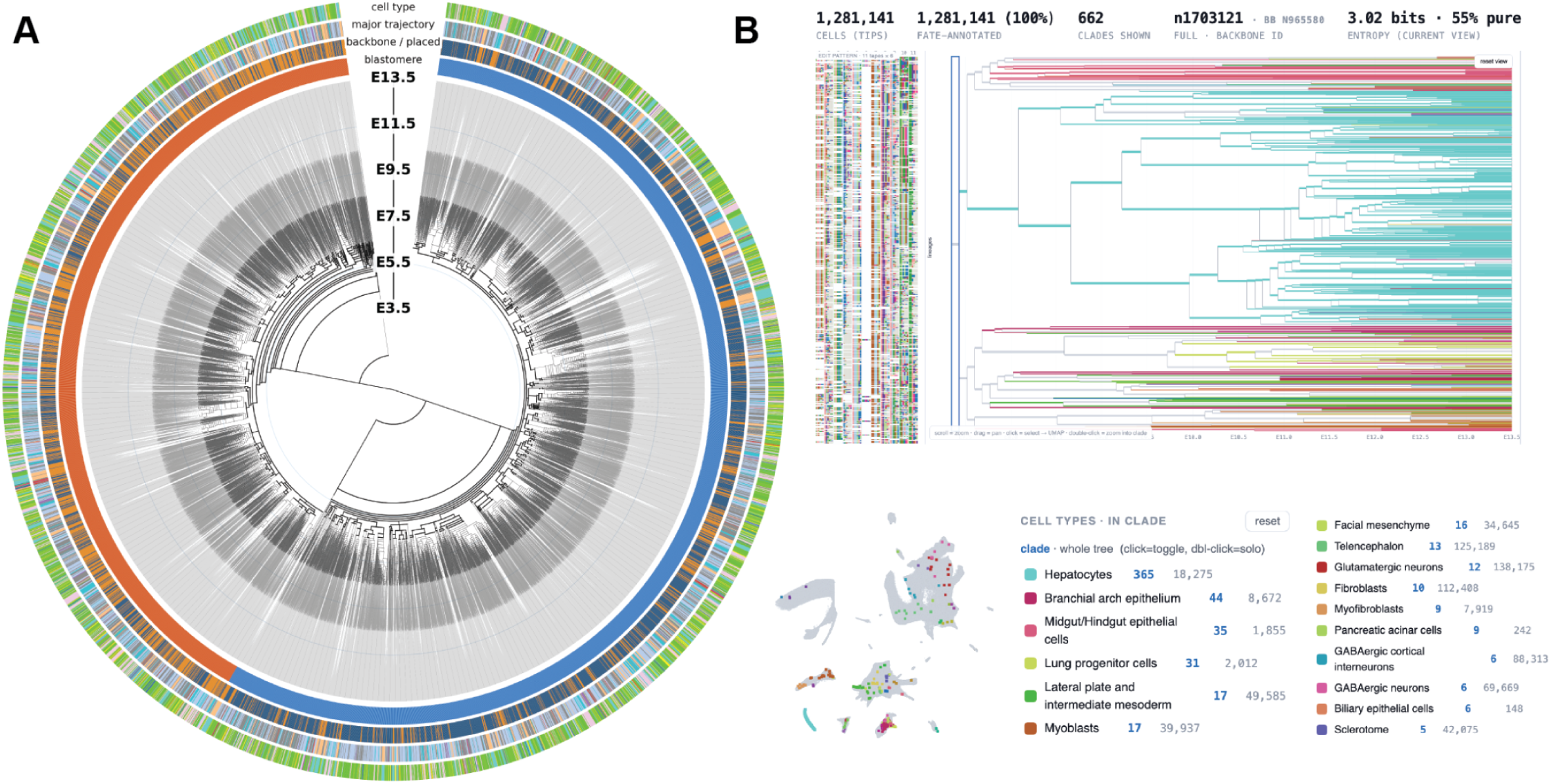
A ∼10^6^ cell phylogeny of a developing mouse from zygote to late organogenesis, and an interactive browser to explore it. **(A)** 1,281,141 cells shown as a circular dendrogram in which radial position gives inferred node age, from zygote (center) to E13.5 (perimeter). Branch widths are scaled in 2-day steps to render the sparse early backbone legible. The two blastomere sub-lineages, A and B, occupy the right and left halves. Concentric rings encode, from inner to outer: (i) blastomere identity (A, blue; B, orange); (ii) reconstruction origin (backbone, dark blue; placement, orange); (iii) major trajectory (legend in Fig. 2A); and (iv) cell type. The red asterisk and wedge mark the clade shown in **B**; the wedge’s angular width is exaggerated ∼5-fold for visibility. **(B)** Interactive visualization in NextCell (*60*), focused on ancestral node n1703121 in the full tree, an endodermally fated progenitor (E7.4) whose 662 sampled descendants are clonally partitioned among cell types associated with visceral organs, including hepatocytes (n = 365), branchial arch epithelium (n = 44), midgut/hindgut epithelium (n = 35), and lung progenitors (n = 31). Panels show the edit states of 11 genotyped tapes (top left); a dendrogram of the subclade, with branches colored where ≥60% of descendant cells share a cell type (top right); a UMAP of all cells in the final tree, highlighting subclade members (bottom left); and cell types observed ≥5 times, with counts in the subclade and among all tips (bottom right).

### A wave of clonal dominance arises during gastrulation

Clonal dominance, the progeny of some progenitors accruing disproportionately relative to others, is recurrent in vertebrate development, mostly studied within individual tissues (*61–65*). At the whole-organism scale, a few early GESTALT-marked progenitors accounted for most non-blood cells in each organ of an adult zebrafish (*21*), while LoxCode barcoding of the E5.5 mouse epiblast revealed clones differing widely in size and fate (*66*). In both cases, however, marks were established at one point and read out at another, such that dominance could be quantified but not dated.

To investigate when dominance arises, we tracked the Gini coefficient of clonal contribution across developmental time, taking lineages present at one timepoint as founders and their descendants at a later timepoint as clones. We defined founders at E7.0, by which point node support has recovered and the ceiling is no longer binding (**Fig. 2K**). This yields 3,849 clones (A: 1,970; B: 1,879), each contributing a mean of 333 profiled E13.5 descendants, but with marked inequality, *e.g.* the 170 largest account for 50% of tips, versus 720 expected under a neutral Yule (pure-birth) null (*67*), while the observed Gini coefficient of 0.80 grossly exceeds the null’s 0.50 (**Fig. 4A-C**). These patterns are similar in the backbone tree (**fig. S13A-C**), similar between A and B (**fig. S13D-F**), and robust to the ratio at which the ceiling is applied to A vs. B during dating (**fig. S14A**).

**Figure 4.**
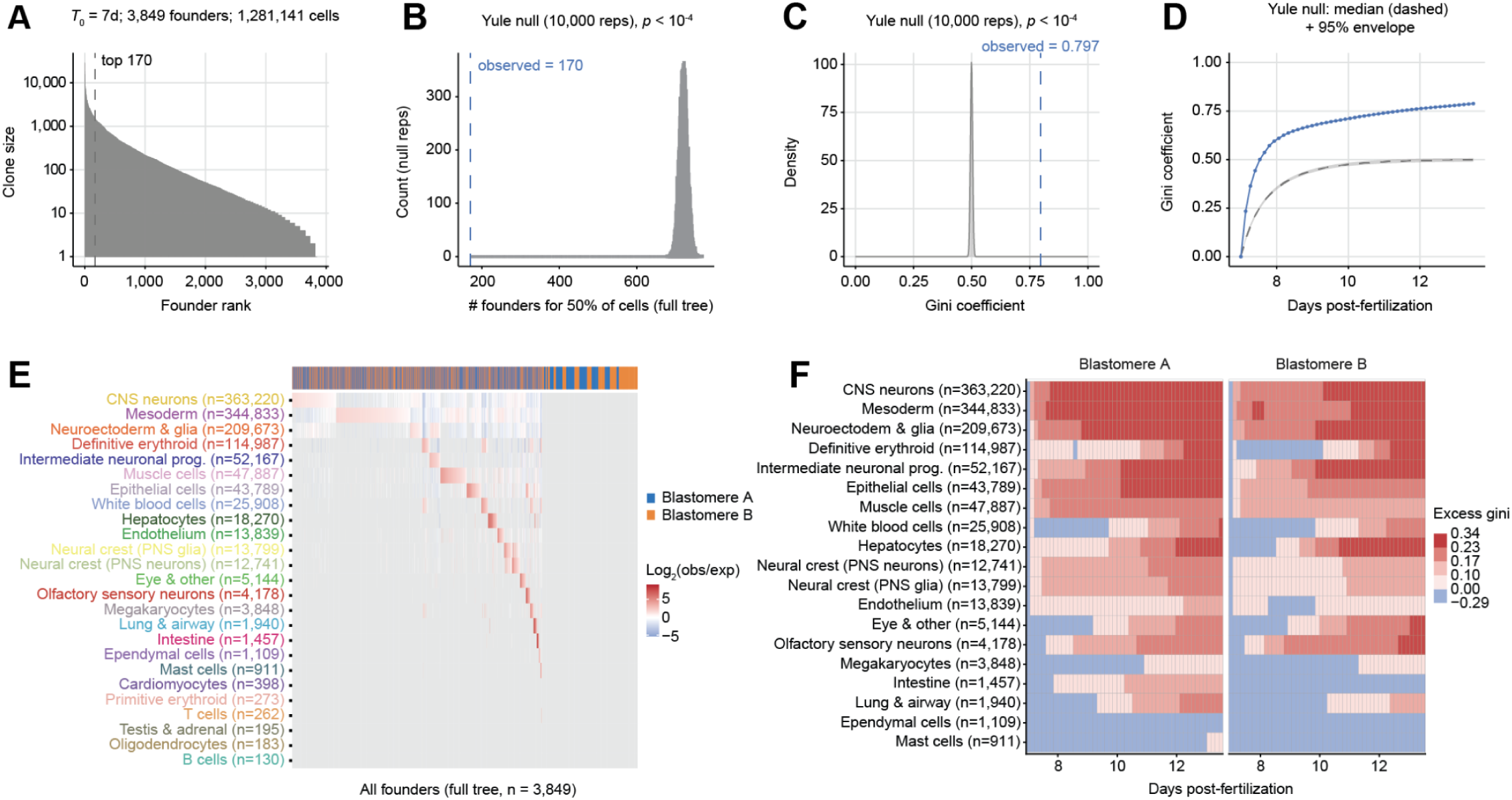
Quantitation of clonal dominance arising during gastrulation. **(A)** Rank-abundance of founder clone sizes (log_10_ scale). Founder clones (n = 3,849) are defined as the lineages present at *T*₀ = 7.0 days post-fertilization in the dated tree; each founder’s clone size is its number of E13.5 descendants. The dashed line marks the top 170 founders. **(B)** Number of founders required to generate 50% of all E13.5 cells, in observed data (blue dashed line, n = 170) vs. the Yule null (grey, 10,000 replicates). *p* = *P* (null ≤ observed). **(C)** Gini coefficient of founder clone sizes on the full tree (0 = every founder contributes equally, 1 = all cells derive from a single founder); the observed value (dashed line) is compared with its distribution under the Yule null (10,000 replicates). *p* = *P* (null Gini ≥ observed). **(D)** Emergence of clonal dominance on the backbone tree. The Gini coefficient computed across the same 3,848 founders’ descendant-lineage counts at each time *T*, tracking the fixed founder set forward from *T*₀ to E13.5 (blue), and comparing against the Yule null (grey) and its 95% envelope (grey). **(E)** Founder clone contribution to each major trajectory relative to its overall contribution to the embryo, full placement tree (n = 3,849 founders, both blastomeres combined). Color shows log_2_(observed/expected) cells of each trajectory (rows, ordered by sampled cells) from each founder (columns, ordered by most-enriched trajectory), where expected assumes a founder contributes to every trajectory in proportion to its overall clone size. Founders are ordered by the trajectory in which they are most enriched, then by enrichment strength within it. Top strip, blastomere of origin (blue, blastomere A; orange, blastomere B); gray, fewer than 10 sampled descendants of that founder in that trajectory. **(F)** Excess Gini, the observed Gini index minus the 97.5% upper bound of the blastomere-specific Gini null, on the backbone tree for major cell-type trajectories with ≥100 cells in both blastomeres (n = 19), over developmental time, split by blastomere of origin. Red, higher inequality than the blastomere’s overall neutral expectation; blue, less. Smaller trajectories may show less signal due to estimator noise, not necessarily true absence of Gini excess.

If we vary the timepoint at which clonal contributions of E7.0 founders are assessed, we find that the Gini coefficient rises steeply and plateaus by ∼E9.0 (**Fig. 4D**; **fig. S14B**). Because the Yule null rises similarly but asymptotes at 0.50, the informative quantity is the departure from it, which exceeds the null’s 95% envelope almost immediately after founders are defined, and remains above it thereafter. Similar dynamics are observed if we define founders at timepoints ranging from E5.5 to E7.5, but not E10.0, from which inequality instead accrues gradually (**fig. S14C-D**).

What might drive it? Inequality exceeding the simplistic, pure-birth Yule null requires net expansion rates that differ between lineages and persist within them, whether through division or survival. Cell cycles around gastrulation are among the shortest of the mammalian life cycle and heterogeneous, fastest in the epiblast’s proliferative zone (*52*, *68*). Over this same interval, apoptosis is elevated in the pregastrula and spatially patterned in the gastrula (*69*), where a substantial fraction of epiblast cells are eliminated by cell competition, which acts on relative fitness and, where the underlying differences are heritable, can drive non-neutral clonal dynamics (*62*, *65*). Here, the 10% most proliferative E7.0-defined founders account for 39% of E8.0-dated lineages, broadly consistent with Snow’s 1977 speculation that the proliferative zone’s contribution could rise from 10% to 50% by E7.5 (*52*). Our tree records only surviving lineages, so it cannot apportion the observed inequality between differential division and differential survival, which are not mutually exclusive. Regardless, the inequality at E13.5 is much smaller than such a skew would produce if fully inherited, suggesting that the plateau marks not the end of heterogeneity but the end of its heritability.

In contrast with A and B, whose contributions to the embryo proper are fate-neutral, many E7.0-defined founders exhibit highly skewed output (**Fig. 4E**; **fig. S14E**), *e.g.* 20 founders account for only 5% of the E13.5 embryo but 51% of its 4,178 olfactory sensory neurons. Resolved by trajectory rather than by founder, nearly all of the well-represented major trajectories exhibit elevated Gini coefficients, in a manner that is reproducible between blastomeres and between the full and backbone trees (**Fig. 4F**; **fig. S14F**). However, founders already occupy distinct positions in the gastrulating embryo by E7.0, so both trajectory-specific dynamics and founder skews plausibly reflect spatial position rather than fate bias, and we are wary of interpreting either further. This also complicates comparison with prospective clonal marking at E5.5 (*66*), both because we define founders 1.5 days later (avoiding the low-confidence pre-gastrulation interval) and because gastrulation-associated clonal dominance and the later-arising fate-specific enrichments are distinct quantities that are likely decoupled.

### Tree siblings share fate far in excess of chance, and mark terminal differentiations

We next turned to the tips of the tree, beginning with tree siblings (cherries and terminal polytomies; **fig. S10F**), to ask how strongly lineage predicts fate at the finest scale available. For annotations observed in both subtrees, we asked how often tree siblings share it, relative to a tip-label-permuted null. Because our tree captures at best ∼10% of the estimated 13-61M cells of an E13.5 embryo (*4*, *5*, *58*), most tree siblings are not true sisters but nearest sampled relatives, so any concordance we observe is a lower bound on true sibling concordance. Globally, 51% of tree-sibling pairs share a cell type label, a 9.4-fold enrichment over a permuted null (5.9-fold for strict two-tip cherries). Enrichment varied widely across fates but was reproducible between blastomeres (Spearman’s ρ = 0.94; **fig. S15A-B**; **tables S4-5**). Extreme concordances corresponded to cell types understood to commit from a small, spatially restricted progenitor pool and then expand clonally within an anatomical niche, *e.g.* lung progenitor cells at 145-fold (**fig. S15B**).

Half of tree siblings were cell-type discordant. To ask whether heterotypic pairs might inform terminal differentiations, we extracted the most closely related subset (shared ancestor dated ≥E12.5; at most one discordant tape monomer) wherein exactly one member was among 27 post-mitotic cell types (**fig. S15C**; **table S6**). This yielded 82,300 post-mitotic cells distributed across 640 progenitor-derivative pairings. Against a within-blastomere permutation null, 37 pairings were ≥3-fold enriched at 1% false discovery rate (FDR) (max 29-fold), with enrichments reproducible between subtrees (Spearman’s ρ = 0.82; **fig. S15D**). Together these draw on 24 distinct progenitor states and assign at least one progenitor to 18/27 post-mitotic types, and recover canonical differentiation events at the resolution of individual dated divisions (**Fig. 5A**) spanning neurogenesis, terminal maturation, and hematopoiesis; the latter warrant caution, as the blood-specific slowdown may leave our edit-distance threshold less able to isolate recent divisions.

**Figure 5.**
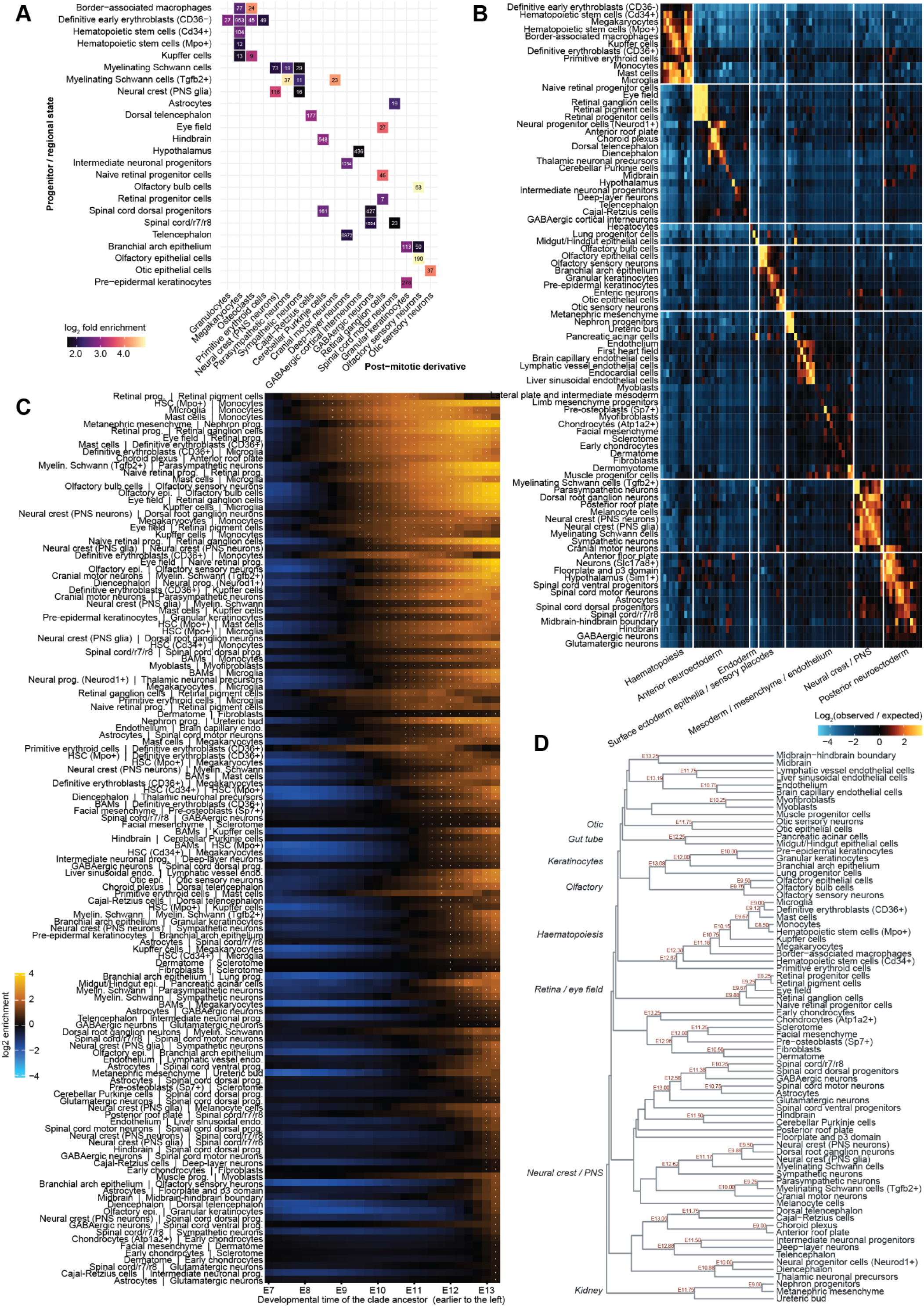
Lineage predicts fate at every scale, from sibling pairs to a dated hierarchy of cell-type couplings. **(A)** Heatmap of the 37 progenitor → post-mitotic heterotypic pairings that significantly and reliably recur among late-splitting tree siblings above chance. Rows, 24 progenitor or regional states; columns, 18 post-mitotic cell types; both ordered by germ layer. Color, log_2_-fold enrichment of the pairing over a tip-label-permuted null. Numbers within cells, post-mitotic cells supporting that pairing (range 7-6,972; 13,549 in total). White, pairing not captured. All pairings shown are supported by at least 3 post-mitotic cells, are at least 3-fold enriched, and clear a Benjamini-Hochberg FDR of 1% across the 640 pairings tested. **(B)** Briefly, we partitioned the tree into “maximal clades” (**fig. S16**), and for each pair of abundant cell types, counted the number of clades containing both (including self-pairs) and compared this to the expected count under random cell-type label shuffling (100 permutations). Log_2_-enrichment (observed / expected), clipped to [−5, 3.5] is shown at K=15, capturing local family groups. Cell types are ordered by hierarchical clustering (Ward’s linkage on Euclidean distance) of the K=15 matrix, with 4 of 88 cell types manually repositioned to make the annotated groupings contiguous (see **Methods** for details of manual repositionings); the same order is used across rows and columns. White lines were manually added and demarcate clusters of cell types corresponding to major groupings shown at bottom. **(C)** Timed fate couplings between cell types. Each row is one of the 129 cell-type pairs with a defined coupling depth. Each column is one of the 26 developmental time slices from E7.0 to E13.25 in 0.25-day steps, with earlier ancestors to the left. Color, log_2_-enrichment (observed / expected under tip-label permutation null) at that slice, averaged over timings calculated on each blastomere subtree independently. White dots mark slices at which the pair is co-enriched in both blastomere sub-trees independently. The leftmost white dot in each row therefore corresponds to a given pair’s coupling depth. Rows are ordered by coupling depth, deepest first, then by peak enrichment. Within each pair label, the cell type appearing earlier in the Qiu *et al.* developmental graph (*5*) is listed first (a presentational guidepost, not a claim of directionality, which this analysis does not compute). Source values are provided in **table S7**. **(D)** A draft hierarchy of timed fate couplings. Average-linkage hierarchy over the matrix of pairwise coupling depths, built on 75 cell types that couple with at least one partner. The tree is drawn unscaled and cannot be read as a phylogeny; horizontal distance is merge rank, not time. Relatedly, the hierarchy appears inverted with respect to a lineage splitting through time, because branches here are joined, rather than split, at their deepest coupling. Of its 74 branch points, the 51 that close at or before E13.25, the last slice, are labelled with that coupling depth; the remaining 23 merge only beyond the last slice and so carry no coupling depth. As coupling depth reports how far back two fates remain confined to a shared pool of lineages, not when they diverged, groups closing earliest stay co-restricted deepest. Labels at the right of cell types name manually curated coherent groups. 13 of 88 cell types (≥100 cells in each blastomere subtree) couple with no partner at any slice by this method and are omitted.

### A hierarchy of fate couplings dated from gastrulation to organogenesis

Tree siblings probe only the immediate neighborhood of each tip. To reach older relationships, we partitioned each blastomere subtree into “maximal clades” (*70*), *i.e.* subtrees of ≥3 but no more than K cells whose parent exceeds K, and asked which of 88 abundant cell types co-occur within them more often than expected under a tip-label-permuted null (**fig. S16**) (*27*, *71*). We ran this at K=15 (median ancestor age E11.2) and K=200 (E8.8).

Hierarchical clustering of the K=15 enrichment matrix revealed clear blocks corresponding to hematopoiesis, anterior and posterior neuroectoderm, neural crest, mesoderm/mesenchyme/endothelium, endoderm, and surface ectoderm epithelia (**Fig. 5B**). Because the tree was built without reference to cell identity, recovering this organization from clade co-occurrence alone further indicates that its topology reflects genuine lineage relationships, as does its reproducibility between the A and B sub-trees (off-diagonal Spearman’s ρ = 0.79 at K=15; **fig. S17A**).

Block structure was much sharper at K=15 (recent clades; **fig. S17A**) than K=200 (ancient clades; **fig. S17B**). Of 3,828 cell-type pairs, 143 were significantly enriched in both blastomeres at K=15, but only 22 at K=200, and 18 at both (**fig. S17C**). Whereas pairs enriched only at K=15 overwhelmingly reflect post-gastrulation relationships, those enriched at both remain coupled within clades whose common ancestor predates E9, implying couplings dating to gastrulation or earlier. That some couplings persist across a 13-fold change in the clade-size cap while most do not, suggested to us that the depth at which an enrichment disappears is itself informative, and that sweeping developmental time directly might allow the coupling of cell-type pairs to be dated systematically.

To this end, for each time *t* from E7.0 to E13.25 in 0.25-day steps, we defined the clade of every lineage present at *t* as all of its sampled descendants, and computed the same pairwise co-occurrence enrichment as above against a tip-label permutation null. For each qualifying pair we recorded its coupling depth, *i.e.* the earliest *t* at which the two remained significantly co-enriched in both blastomere subtrees. This yielded coupling depths for 129 cell-type pairs, touching 75/88 cell types with dates spanning E8.25-E13.25 (**Fig. 5C**; **table S7**). Significance formed a single contiguous interval in time for 122 of these 129 pairs (95%). Average-linkage clustering of these depths yielded a hierarchy in which most branch points carry a developmental date; clusters generally corresponded to major germ layers or organs, and the most deeply coupled groups “closed” first (**Fig. 5D**). The order and dating of the cell type couplings were biologically sensible and reproducible between blastomeres (**fig. S17D**).

Because our placement heuristic could in principle manufacture the terminal structure these analyses rest on, and because global tip-label permutation is a permissive null, we repeated the sibling, heterotypic-pairing and clade co-occurrence analyses on the backbone tree alone, and against a within-clone null (**Supplementary Text, note S4**; **tables S5-7**). Enrichments fell in magnitude but persisted, and the rank order of coupling depths was preserved throughout, while their absolute dates shifted by a median of 0.75 days, deeper on the backbone and shallower under the within-clone null. We conclude that sibling enrichments and the ordering of cell type couplings are robust to both tree and null, but also that absolute coupling dates should not be overinterpreted.

### Ancestral state imputation recovers known developmental transitions

Each of the tree’s 1,281,141 tips carries a transcriptional state, but its 1,109,894 internal nodes are unobserved. We sought to develop an exploratory, heuristic imputation strategy that leverages our scRNA-seq atlas of mouse development (*5*). Within individual cell types, tree siblings are on average closer in transcriptional state than non-siblings of the same type, and transcriptional similarity tracks lineal relatedness (**Fig. 6A-B**). However, these trends are subtle, such that we favored approaches that aggregate information across many descendants of each internal node.

**Figure 6.**
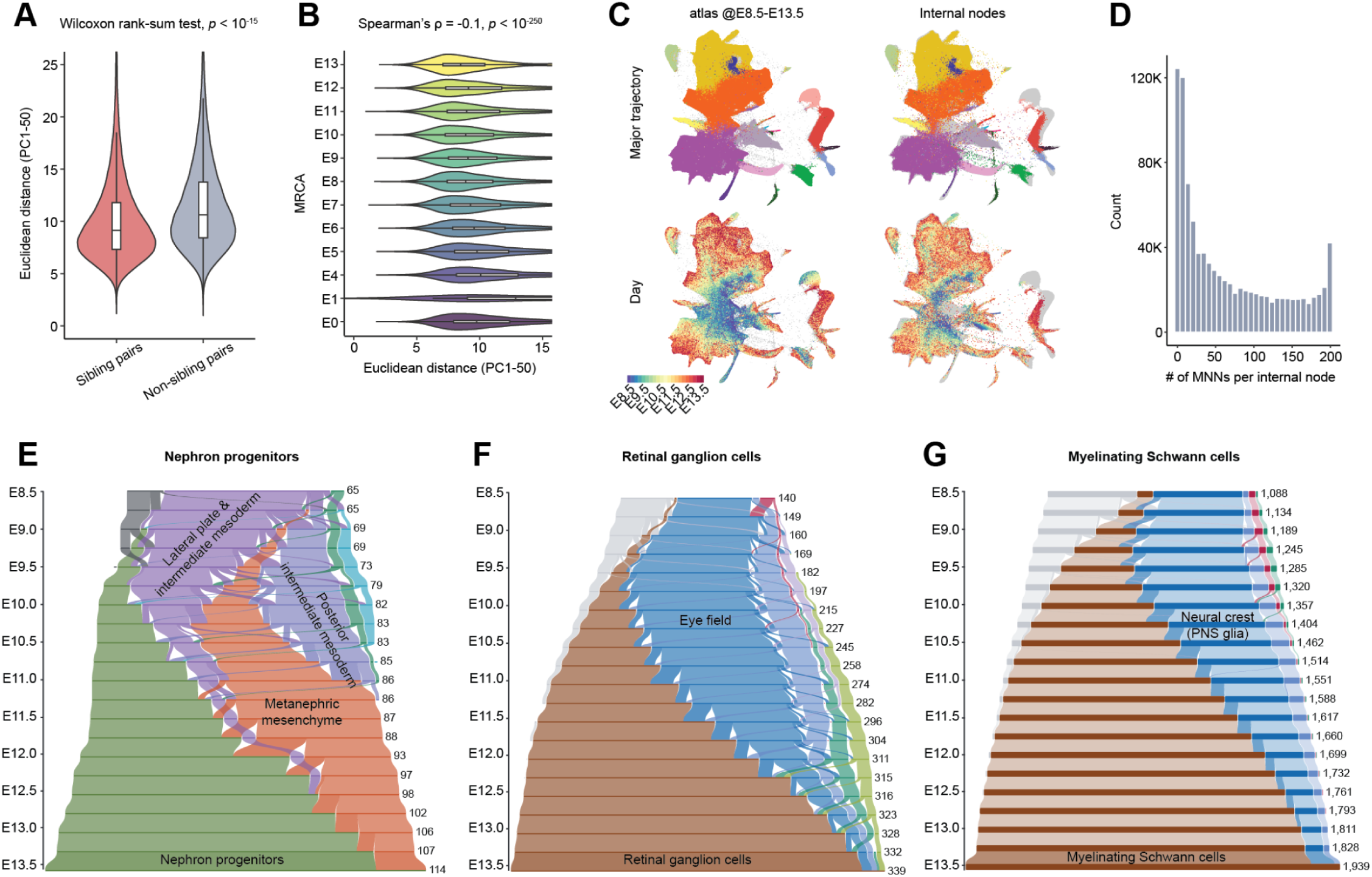
Heuristic inference of the transcriptional states and cell type annotations of ancestral nodes. **(A)** For each cell type, up to 10,000 sibling pairs were randomly sampled, along with a matched number of non-sibling control pairs of the same cell type (excluding known sibling and self-pairs). Transcriptional distance was defined as the Euclidean distance in the first 50 principal components (PCs). Inner box plots indicate the median and interquartile range. **(B)** Same-cell-type pairs, binned by most recent common ancestor (MRCA) depth on the lineage tree, were sampled at up to 10,000 pairs per unit-depth bin. Transcriptional distance was defined as the Euclidean distance in the first 50 PCs. Inner box plots indicate the median and interquartile range. Spearman’s ρ between transcriptional distance and MRCA unit-depth bin are reported. **(C)** 2D UMAP co-embedding of 1,973,105 single-cell profiles, comprising 926,852 internal nodes with imputed transcriptomes from embryo #3 and ∼1 million cells from 21 timepoints of our high-temporal-resolution single-cell atlas of mouse development (E8.5–E13.5, 6-hour increments; each timepoint downsampled to 50,000 cells). UMAP was first computed on the atlas cells, and internal nodes were then projected onto this embedding. Both internal nodes and atlas cells were represented in a joint 50-dimensional PCA space, computed by integrating embryo #3 (all E13.5) with ∼5.2 million single-cell profiles from the atlas (E8-E13.5); each internal node’s transcriptome was defined as the mean of the PCA coordinates of its assigned MNNs. **(D)** Histogram of the number of MNNs assigned to each of 926,852 internal nodes. **(E-G)** Examples illustrating ancestral state views in NextCell (*60*). **(E)** Developmental origins of a cell type at E13.5, illustrated for nephron progenitors. Cells of the indicated type were traced back through the lineage tree, and at each 0.25-day slice from E13.25 to E8.5, every lineage was assigned the inferred state of its nearest annotated ancestor; ancestral state calls extend back only to E8.5. Cells whose lineage carries no called ancestor, and those on rare paths (carrying <1% of the cell type’s cells), are set aside rather than plotted, leaving 114 of 607 nephron progenitors. Vertical position indicates developmental time. Bar width is proportional to the number of ancestral lineages, given at right, and segments are coloured by inferred state (up to the 8 most abundant shown individually, the remainder grouped as “Other”; grey, the portion of a lineage earlier than its own earliest called ancestor). Within each gap, one ribbon per state links lineages between adjacent slices (faint for same-state, solid for a state change) so ribbon crossings mark inferred state transitions. Single-slice A→B→A excursions (0.25 d) were removed. Going backward in time, lineages coalesce into fewer ancestors; the dashed bottom row is the E13.5 cells themselves, one lineage per cell and therefore a single state by construction. **(F)** As in panel **E**, for retinal ganglion cells (339 of 1,645 plotted). (G) As in panel **E**, for myelinating Schwann cells (1,939 of 7,663 plotted).

Traversing from tips to root, we assigned each internal node the mean transcriptional profile of its children, then linked ancestral nodes to atlas cells by mutual nearest neighbors in successive 6-hour windows from E13.5 back to E8.5, assigning each matched node an updated profile and a cell type by majority vote (**Fig. 6C-D**). Critically, the lineage relationships here are given by the tree, with the atlas serving only to identify the contemporaneous state most consistent with each ancestral node’s descendants.

This heuristic yielded cell type labels for 926,852 of the 1,054,879 internal nodes within the atlas interval (88%), with coverage lowest early (46% at E8.75) and best late (96-98% between E11.5 and E13.0). Imputed labels are robust to random masking of 20% of tips, albeit less so for rare annotations (**fig. S18**), and are almost entirely lost when cell identities are permuted across tips (0.06% label agreement). But they are also noisy, implying 132,536 distinct “state paths” that are overwhelmingly rare and biologically implausible.

We therefore defined a state path as “qualifying” if it was heterotypic, non-recursive, carried by at least 5 instances and ≥1% of its terminal cell type’s traced cells, and composed of constituent steps that were individually well-supported (≥50 overall transitions) and asymmetric (>60/40 preference for observed direction) in the overall tree. This yielded 275 qualifying state paths, 98% of which are recovered independently in both blastomeres, covering 82 of 136 cell types and 117,704 tips (**table S8**). These criteria use only the phylogeny and imputed labels, and no external knowledge of developmental biology.

Although qualifying state paths are short (193 with two states, 77 with three, 5 with four), imputed rather than recorded, and lack validation, they are clearly enriched for well-established differentiations (*e.g.* lateral plate and intermediate mesoderm → posterior intermediate mesoderm → metanephric mesenchyme → nephron progenitors). To evaluate this systematically, and with the caveat that this is not fully independent validation, we scored imputed paths against our previously reported, manually curated cell-type transition graph (*5*). Of 360 testable steps, 160 (44%) connect states that are directly adjacent in the 474-edge transition graph and 207 (58%) connect states within two edges of one, 15.7-fold and 7.4-fold enrichment over chance. Imputed annotations and qualifying paths can also be explored in NextCell (*60*), including as temporal flow-allocation diagrams (*9*) (**Fig. 6E-G**).

## Discussion

Molecular technologies, like algorithms, have a theoretical scaling that is clean and a practical scaling that is not. At the outset of this field around 2016, we and others imagined that the path from proof-of-concept to a “Sulston mouse” would be relatively straightforward. But the devil, of course, is in the details.

Of the five obstacles described in the introduction that are not addressed by DNA Typewriter itself, we make progress on all but fully resolve none. Delivery by PNI and piggyBac established a stable, constitutively expressed system through E13.5, but was mosaic and uncontrolled in integration number and location (i); PNI offers a faster design-test loop than cell culture or organoids, directly *in vivo*, but does not result in a distributable “recorder mouse” line (ix). sci-RNA-seq3 recovered >10^6^ cells, ∼10% of this embryo but orders of magnitude short of what is required for dense sampling of an adult mouse (vi). Circtape exhibited subjectively improved recovery relative to our prior attempts at targeted transcript recovery from single nuclei (*37*, *40*), yet only 54% of the cell × tape matrix was filled, possibly because circular RNAs are exported to the cytoplasm (*72*) (vii). Finally, although we succeeded in building a time-calibrated, parsimony-supported phylogeny of 1.28M cells, its dating and topology are limited by clock violations and the placement heuristic (viii).

Four further obstacles, not on our list, loomed large in practice. First, cell type annotation of this phylogeny’s tips was tractable only because of a parallel, multi-year effort to generate a highly resolved scRNA-seq atlas of mouse development (*4–6*). Second, a barren phylogeny does not directly inform the identities of ancestral nodes; our co-occurrence and state-imputation heuristics are first approximations, and the toolkit for “developmental statistics” remains sparse (*73*). Third, with a single embryo, reproducibility rests on comparisons of blastomere A vs. B subtrees, a useful but limited form of replication, *e.g.* we cannot assess how integration-mediated disruption of genes such as *Chd3* (*74*) impacted the phylogeny. As such, our results here and any follow-on analyses of the phylogeny should be regarded as preliminary, pending confirmation in independent animals.

The fourth new obstacle was the frustrating-yet-fascinating dynamics of PNI-delivered DNA Typewriter. A massive editing burst, helpful for rooting, spanned the 1-cell (minor ZGA) and 2-cell (major ZGA) stages (**Supplementary Text, note S1**), which may be permissive for genome editing (*75*, *76*). Less helpfully, the editing rate then sharply fell, recovering by gastrulation, albeit not to the level of the 1-cell/2-cell burst. We speculate that post-ZGA epigenetic silencing of integrated cassettes and/or a lower intrinsic editing rate in epiblast precursors, which dominate the embryo proper at this stage, reduce prime editing during this window. Single-cell profiling of extraembryonic tissues from DNA Typewriter embryos may shed further light.

From its perigastrulation recovery through E13.5, DNA Typewriter recorded robustly, although rates varied modestly between cell types. The marked blood-specific slowdown could be experiment-specific (*e.g.* silencing of specific PEmax integration sites), setup-specific (*e.g.* silencing of the CAG promoter driving PEmax) or biological (*e.g.* dNTP limitation in hematopoietic cells); further experiments are required to distinguish these. Regardless, cell type-specific rates violate the strict molecular clock. Looking forward, we favor a transition to maximum likelihood methods that more explicitly accommodate molecular biology, missing data, and complexity in both phylodynamics and recorder dynamics, as well as divide-and-conquer algorithms that may scale to 10^8^ cells and beyond (*43*, *77–80*).

Other avenues for improvement are arithmetic and scale. Recording capacity is the product of tape number and length. Many of the 11 × 6-monomer tapes were saturated by E13.5, eroding the rate stability that DNA Typewriter’s sequential recording affords. By contrast, 50 × 20-monomer tapes would afford 1,000 positions, enough to stably record to adulthood. Sampling DNA Typewriter embryos at more stages, in conjunction with chemical, genetic or environmental perturbations, and/or with integration of signal-, state- and spatially-resolved recording, would each add dimensions that our single tree lacks (*36*, *81*, *82*).

Large, dense cellular phylogenies of regulatively developing animals are a genuinely new kind of data object, enabling investigation of the statistical regularities that recur within an individual, or hold across individuals, even when no single lineage does. As we have argued elsewhere, such phylogenies are poised to anchor the “consensus ontogeny” that we envision for organisms such as the mouse, a lineage-anchored reference onto which multimodal measurements can be layered (*83*). The 1.28M tree reported here, one embryo at one stage comprising ∼10% of its cells, is at best a single viewpoint on such an ontogeny. Yet it recovers, from clonal co-occurrence alone and without reference to transcriptional state, a dated hierarchy of cell-type couplings that agrees with canonical relationships of mouse development accrued over decades of investigation. That so much can be read from a single, incomplete lineage is itself encouraging. We have only scratched the surface of what might be learned from this one dataset, and we welcome its further exploration, and its integration with other cellular phylogenies of the developing mouse (*84*).

Following years of work across many groups (*19*), we hope that this and contemporaneous studies (*84*) mark an inflection point for the molecular recording field. As more sophisticated *in vivo* recording devices are brought to bear, the phenotypic equivalent of a mammalian reference genome, from zygote to birth to adulthood to death, comes into view (*83*).

**The materials and methods are available in the supplementary materials.**

## Acknowledgements

We thank the members of the Shendure lab, the Seattle Hub for Synthetic Biology, Allen Discovery Center for Cell Lineage, and UW Genome Sciences communities for helpful discussions over the course of many years (particularly the Shendure/SeaHub Recording subgroup, A. Schier, M. Elowitz, R. Waterston, and W. Colgan). We also thank the Transgenic Colony Management, Neurosurgery & Behavior, Lab Animal Services, and the Veterinary Services teams at the Allen Institute for their contributions to animal work. QY, HK and SS contributed equally to this work as co-first authors. When referencing this paper in their CVs or other professional materials, each co-first author may list their own name first. Several authors used AI-based tools during this work. The NextCell interactive browser was built with Claude Code (Anthropic). Data exploration, data analysis, data visualisation, and code development were additionally supported by Claude Opus 5, Claude Opus 4.7, and Claude Sonnet 5 (Anthropic), which were also used to improve the readability and language of the manuscript. AI-assisted tools were not used to generate or alter primary data, to draw scientific conclusions, or to perform other key authoring tasks. All AI-assisted code, analyses, and text were reviewed, tested, and edited by the authors, who take full responsibility for the data, code, analyses, and conclusions presented here. The content is solely the responsibility of the authors and does not necessarily represent the official views of the NIH. This research was supported in part by the Brotman Baty Institute for Precision Medicine, as well as the Allen Institute, founded by Jody Allen – chair and co-founder of Allen Family Philanthropies, and the late Paul G. Allen – investor, philanthropist, and co-founder of Microsoft. We gratefully acknowledge their vision and generosity, which make this work possible. J.S. is an Investigator of the Howard Hughes Medical Institute.

## Funding

Allen Institute (JS)

Brotman Baty Institute for Precision Medicine (JS)

Cancer Research Society of Canada grant 1155581 (J-BL)

Chan Zuckerberg Initiative Foundation, Seattle Hub for Synthetic Biology award CZIF2023-008738 (JS, CT, MP)

Charles H. Hood Foundation, Inc., MA (CQ)

Damon Runyon Cancer Research Foundation grant DFS-64-24 (JC)

Damon Runyon Cancer Research Foundation grant DRG-2435-21 (J-BL)

Howard Hughes Medical Institute (JS)

National Institutes of Health grant R00HG012973 (JC)

National Institutes of Health grant P30CA008748 (JC)

National Institutes of Health grant P20GM130454, via Dartmouth’s Center for Quantitative Biology (CQ)

Paul G. Allen Frontiers Group, Allen Discovery Center for Cell Lineage Tracing (JS, CT)

Searle Scholars Award SSP-2025-101 (JC)

Swiss National Science Foundation Postdoc. Mobility fellowship grant 239394 (SS)

Washington Research Foundation Postdoctoral Fellowship (HK)

## Author contributions

Project initiation: QY, JC, JS

Circtape development: HK, with assistance from JFN, QY, J-BL, KPS

Design of *in vivo* experiment: QY, JFA-C, MG

Design and execution of ddPCR: EG, LK, SVK

Pronuclear zygotic injection platform development: JFA-C, KO

Pronuclear zygotic injections and embryo manipulations: KO, QY

Bulk amplicon sequencing and data analysis: QY, HK, RMD, JS

Sci-RNA-seq3 experiments: BKM, QY

Sci-RNA-seq3 data analysis: CQ, with assistance from JS, SS, CT

Phylogenetics: SS, with assistance from JS, CQ, ML

Analysis of timed, tip-annotated trees: SS, CQ, JS

Software: SS, CQ, JS

Funding acquisition: JS, CT, MP

Project administration: JS, JMG, JFA-C

Visualization: CQ, SS, HK, QY, JS

Manuscript writing: SS, CQ, HK, QY, JS, with assistance from JFA-C, BKM, RMD

Project oversight: JS

## Data, code, and materials availability

Bulk and scRNA-seq sequencing data are available at Gene Expression Omnibus with identifier GSE341627. Intermediate data files (*e.g.* full and backbone trees in Newick format) are available at NextCell (*60*) or at our GitHub site for this project (*85*). All other data are available in the main text or the supplementary materials. Code is available at our GitHub site for this project (*85*). PEmax and epegRNA::Tape-BC::circtape constructs are available from author JS and from Addgene.

## Supplementary Materials for

### Materials and Methods

#### EXPERIMENTAL METHODS

##### Molecular cloning

###### PEmax plasmid

The chicken HS4 insulators, CAGGS promoter, and PEmax-P2A-dTomato-WPRE-bGHpA cassette were synthesized as gene fragments with compatible overhangs (GenScript). These fragments were excised using PaqCI restriction enzyme (NEB, R0745S) and assembled into a piggyBac transposon vector via Golden Gate assembly.

###### epegRNA-circtape plasmid library

A partial segment of epegRNA, including the region encoding the partially degenerate insertion (NNNGGA), was synthesized as a library (IDT, Ultramer). The 6-monomer tape and the remaining regions—including the two ribozymes required for the Tornado circularization system—were synthesized as a DNA oligonucleotide (Ansa Biotechnologies) and clonal DNAs (GenScript), respectively (*87*). These components were amplified by PCR and cloned into a piggyBac transposon vector by Gibson assembly using NEBuilder HiFi DNA Assembly (NEB, E2126S). Next, an oligo library for the tape barcode (NNNNNAANNNNN; AA was incorporated to exclude the possibility of a BsaI recognition site arising) was synthesized by IDT, amplified by PCR, and cloned into the epegRNA-circtape construct via Golden Gate assembly using NEBridge Ligase Master Mix (NEB, 1100S) and BsaI-HFv2 (NEB, R3733L). The sequences of synthetic oligonucleotides are listed in **data S1**.

##### Animal husbandry

All animal experimentation was conducted according to the protocols approved by the Institutional Animal Care and Use Committee of the Allen Institute (Protocol #2530 and #2401). Mice were housed in a dedicated animal vivarium operating on a 14hr-10hr light-dark cycle and had access to food and water ad libitum.

##### Zygote collections, pronuclear injection, and embryo transfer surgeries

Superovulation, zygote collection, pronuclear zygotic injection (PNI), and embryo transfer were performed as detailed in Manipulating the Mouse Embryo: A Laboratory Manual (4^th^ ed.; Behringer, Gertsenstein, Nagy, and Nagy, 2014). In brief, 4-week-old C57BL6/J females (The Jackson Laboratory, 000664) were superovulated via intraperitoneal injection of 5 I.U.s of human chorionic gonadotropin (hCG; Sigma-Aldrich, CG5) and 5 I.U.s of pregnant mare serum gonadotropin (PMSG; Ilex Life Sciences, A22725K). Superovulated females were then paired with fertile C57BL6/J males overnight to generate fertilized zygotes. E0.5 zygotes were collected from superovulated females the morning after mating. E0.5 zygotes were injected with a mixture of piggyBac transposon mRNA—synthesized with a Cap-1 structure, a poly(A) tail, and N1-methyl-pseudouridine (m1Ψ) modification to enhance mRNA stability and translational efficiency (GenScript, RP-A00064-0.2)—and plasmid DNA in sterile, filtered IDTE (IDT, 11-01-02-02). For the initial PNI session that yielded embryo #3, the final injection mix contained 100 ng/µl piggyBac mRNA, 20 ng/µl PEmax plasmid, and 10 ng/µl epegRNA::tape-BC::circtape plasmid library. For the PNI sessions that yielded the embryos described in **note S1**, the final injection mix contained 100 ng/µl piggyBac mRNA, 25 ng/µl PEmax plasmid, and 5 ng/µl epegRNA::tape-BC::circtape plasmid library. The male pronuclei of zygotes were injected and immediately transferred at the 1-cell stage into the oviducts of pseudopregnant CD-1 recipient females (Charles River, 022).

##### Harvesting and processing of E13.5 embryos

Once embryos reached the desired gestation stage, surrogate CD-1 females were euthanized and embryos were harvested, rinsed in 1x PBS, dabbed dried and immediately flash frozen in liquid nitrogen. Nuclei of E13.5 embryos were then isolated and fixed following the sci-RNA-seq3 protocol.

##### Bulk amplicon-seq of genomic tape

10% of the fixed nuclei from each embryo were subjected to genomic DNA (gDNA) extraction using Qiagen AllPrep DNA/RNA kit (Qiagen, 80204). Bulk amplicon-sequencing libraries spanning the tape region or the epegRNA-tape region were generated from the purified gDNA using two-step PCR and sequenced on a NextSeq 2000. An estimated 5 ng (or 50 ng, for additional embryos described in **note S1**) of purified gDNA was used for the first PCR with 0.4 µM of forward and reverse primers that included sequencing adaptors at their 5′ ends in a final reaction volume of 25 µl. In the first PCR, samples were incubated for 3 min at 95 °C; 15 s at 95 °C, 10 s at 65 °C and 45 s at 72 °C for 25–28 cycles; and 1 min at 72 °C. Products of the first PCR were purified using 1.0x AMPure beads (Beckman Coulter, A63882) and added to the second PCR that appended dual sample index sequences and flow cell adaptors. The second PCR used the same thermocycling program as the first, with the cycle number reduced to 6–8. Products were again purified using 1.0x AMPure beads and assessed on a TapeStation D1000 chip (Agilent, 5067-5582 Screentape and 5067-5583 reagent) before the sequencing run. Sequencing was performed on a NextSeq 2000 with a read structure of 318/10/10 or 200/10/10 (Read 1/Index 1/Index 2) for tape libraries, and 70/10/200 (Read 1/Index 1/Read 2) for epegRNA-tape libraries, using standard Illumina primers (TruSeq Read 1, Nextera Index 1, TruSeq Index 2). The sequences of synthetic oligonucleotides are listed in **data S1**.

##### ddPCR on transgene copy number variation

To count PEmax integrations, 5 ng of purified gDNA was used as input for ddPCR. BioRad 2X ddPCR Supermix for Probes (BioRad 186-3023) was used according to the manufacturer’s protocol. Primers specific to a region of the Cas9 sequence were designed and validated. GAPDH ddPCR Copy Number Determination Assay (BioRad Assay ID dMmuCNS559838272) was used as the reference. The Cas9 probe was detected using FAM and the GAPDH probe was detected using Cy5.5. Assays were run on a QX600 AutoDG Digital Droplet PCR System (BioRad 17008371) and analyzed using QX Manager V2.2 Standard Edition software (BioRad). The sequences of the oligonucleotides, probes, and amplicons are listed in **data S1**.

##### scRNA-seq (sci-RNA-seq3) of E13.5 embryos

Sci-RNA-seq3 was performed on fixed nuclei as previously described (*39*) with the following differences: A new set of indexed reverse transcription primers and indexed ligation primers was used to widen the combinatorial space. (**data S2**). The nuclei were fixed in two steps, 1 ml of resuspended nuclei was fixed first with 4 ml methanol 15 min on ice, and then rehydrated with Sucrose/Phosphate Buffered Saline/Triton X-100/MgCl2 (SPBSTM) buffer containing 2.5 mg/ml BS3 (bis(sulfosuccinimidyl)suberate) (ThermoFisher #21580) for a second fixation for another 10 min. In the first experiment, fixed nuclei (∼24M of embryo #2, ∼10M of embryo #3) were then split over 6 x 96-well plates for reverse transcription, incorporating the first of three indexes. The nuclei were then pooled and split over 6 fresh plates for ligation of the second index. They were pooled again, counted, and distributed to 23 new plates, aiming for 400,000 nuclei per plate. In a second, smaller, experiment, an additional 4M nuclei from embryo #3 were processed similarly, adding 2 more final plates for 25 plates total. These plates were then subjected to second strand synthesis, and the nuclei disrupted with protease (Qiagen #19157). At this point, the freed cDNA is split, with 1/3 continued with the usual sci-RNAseq3 protocol and fragmented with Tn5 transposase (Diagenode #C01070010-100) loaded with

Nextera N7 oligos, and then amplified with sci-RNAseq3 PCR primers. The remaining 2/3 underwent a separate amplification specific to the tape. First, an indexed primer (CAAGCAGAAGACGGCATACGAGAT##########GAAATAGGCCTGCATTAGTCTTGAGG), was used to perform 5 cycles of linear PCR at 60C annealing, with either KAPA Robust polymerase (Fisher #50-196-5219) or NEB OneTaq (#M0484L). After these linear cycles, the other primer was added for an additional 17 cycles of exponential PCR. Each plate of sci-RNA-seq3 PCR and tape PCR was pooled by plate, and size-selected by a dual-AMPure XP (Beckman Coulter #A63882) purification. The sci-RNA-seq3 and tape library pools were sequenced at a ratio of (∼80% sci-RNA-seq3/∼20% tape). Sequencing was performed on either Illumina NextSeq 2000 or NovaSeq X with custom primers. Read1 - 35 cycles, primer: Truseq_R1. Index1 - 10 cycles, primers: Nextera_idx1, tape_seqindex1. Index2 - 10 cycles, primer: Truseq_idx2. Read2 - 183 cycles, primers: Nextera_read2, tape_seqread2. The sequences of synthetic oligonucleotides are listed in **data S2**.

#### COMPUTATIONAL METHODS

##### Primary analysis of bulk tape amplicon-seq data

###### Input sequencing data

Bulk tape amplicons were generated by PCR from gDNA purified from embryo nuclei, and sequenced for each embryo in which tape was detected, giving one amplicon FASTQ per embryo. Embryos #2 and #3 were single-end sequenced, and #6 was paired-end sequenced. For #6, the amplicon is carried on R2, so we reverse complemented it and then processed it identically to embryos #2 and #3. Each amplicon read spans a 12-bp integration barcode, the six-monomer array, and the 3′ terminal landmark.

###### Integration barcode and insertion vocabulary derivation (tape/reference.py)

We defined two reference sets per embryo that are used for all downstream analysis (and, for embryo #3, reused for the single-cell data; see below). (i) A whitelist of tape integration barcodes (a 12-bp tag, NNNNNAANNNNN, that falls between GGCATG and GCTCTGG flanks), derived from ∼2M sampled reads as the 12N barcodes above ≥0.2% frequency and separated by ≥3 mismatches, to exclude single-mismatch “shadows” of abundant barcodes. (ii) An insertion vocabulary of monomer edit symbols. Because real edits are clonal and therefore concentrate at a specific integration and monomer, we counted each candidate insertion per (integration barcode × monomer position) across all reads. We admitted insertions that reached ≥10,000 reads at some single (barcode × position) combination to the insertion vocabulary of a given embryo.

###### *Read parsing* (tape/parser.py)

Each read’s integration barcode is extracted and corrected to the nearest whitelist entry within 2 mismatches (only when unambiguous). The 6-monomer array is resolved both forward from the 5′ anchor (using the fact that editing is strictly ordered, so once a monomer is unedited all downstream monomers are expected to be unedited) and backward from the 3′ terminal landmark (which recovers the distal monomers of heavily edited molecules whose forward read-through might otherwise truncate). Reads with an edit following an unedited monomer (ordering violations) are discarded. A read is retained as valid only if its barcode is whitelisted, all six monomers are resolved, and every edit matches the insertion vocabulary. Valid reads are tallied per integration barcode as counts over their 6-monomer genotype.

###### *De-noising* (tape/denoise.py)

Barcodes with <2,000 total valid reads were not analyzed. Within each retained integration barcode, we removed amplification and sequencing artifacts in four steps: (i) Error-collapse, folding a rare genotype into its most abundant single-mismatch neighbor, where that neighbor was at least 2-fold more abundant. Here a neighbor is a genotype differing by a single base substitution within one monomer’s insertion symbol; an unedited monomer, carrying no sequence to misread, is never mutated into an edited one. Genotypes were processed rarest-first, so chains of errors collapse onto a common parent rather than onto each other. (ii) Chimera removal, discarding fully-edited genotypes explained as the prefix of one abundant genotype joined to the suffix of another (partially-edited genotypes are exempt, since their unedited tails are shared legitimately). (iii) A minimum of 2 supporting reads. (iv) A cross-barcode filter removing deeply-edited genotypes (≥3 edits) whose read support was highest in a different integration barcode, which flags integration-barcode swap recombinants.

###### Integration copy number (tape/copynumber.py)

piggyBac integrations are sometimes duplicated (*88*). Because every barcode is sampled from the same cells at the same sequencing depth, a barcode’s total read abundance reports its genomic copy number, *i.e.* single-copy integrations share one abundance level, and barcodes carried as multiple genomic copies sit at integer multiples of it. We estimated the single-copy level iteratively -- beginning from the median abundance across all barcodes, assigning provisional copy numbers, and re-estimating the level as the median abundance of those barcodes assigned a single copy -- and then assigned each barcode an integer copy number -- round(abundance / level) -- with a floor of one. The total number of integrations is the sum of copy numbers, which exceeds the number of distinct valid barcodes when any are multi-copy, and genomic-equivalent counts were divided by copy number to give cell numbers.

###### Cell-number estimation (tape/depth.py)

Because bulk data lack UMIs, we estimated genomic equivalents from read depth. For each integration in a given embryo, we took λ, the single-genomic-equivalent read depth, as the mode of the per-genotype read-count distribution in log_10_ space, estimated by Gaussian kernel density (bandwidth 0.3) over the genotypes supported by ≥200 reads. The number of genome equivalents (cells) carrying a genotype was round(reads/λ), and the genomic equivalents per integration was their sum.

###### *Per-integration lineage trees* (tape/editchain.py + tape/tree.py)

For each integration in embryo #3, we built a prefix tree over the ordered edit symbols, in which every root-to-tip path traces a lineage’s editing history and tip depth equals the number of monomers written. Reads representing 3′-truncated versions of a deeper genotype were attributed to that deeper lineage (folded onto the tree only when the deeper lineage was ≥4× more abundant, so that genuinely silenced shallow lineages are retained), yielding one lineage tree per integration barcode.

##### Primary analysis of sci-RNA-seq3 sequencing data

For the sci-RNA-seq3 experiments, data were processed independently in batches of four PCR plates for read alignment and gene count matrix generation using our previously described sci-RNA-seq3 pipeline (*4*, *89*) with minor modifications to accommodate new RT and ligation oligos. The first experiment comprised 23 PCR plates divided into six processing batches, and the second experiment comprised two PCR plates processed as a single batch. Nuclei of embryos #2 and #3 were co-processed in the first experiment, occupying different reverse-transcription wells of the same six RT plates, so each cell is assigned to its embryo by its RT barcode. Base calls were converted to FASTQ format using Illumina *bcl2fastq*/v2.20 and demultiplexed by PCR i5 and i7 barcodes using the maximum-likelihood demultiplexer *deML* (*90*) with default settings. Downstream processing followed our sci-RNA-seq protocol (*91*). Briefly, PCR-indexed demultiplexed reads were further filtered on RT and ligation indices (ED < 2) and adapter-trimmed with *trim_galore*/v0.6.5 (*92*) using default settings. Trimmed reads were aligned to the mouse reference genome (mm39) using *STAR*/v2.6.1d (*93*) with default parameters and GENCODE vM37 gene annotations. Uniquely mapping reads were retained, and duplicates were collapsed using the combination of UMI sequence (ED < 2), RT index, ligation adaptor index, and read 2 end coordinate (*i.e.* reads sharing an RT index, ligation index, pcr index, tagmentation site, and UMI within ED < 2 were treated as duplicates). Mapped reads were then split into constituent cellular indices by the combination of all three indexes. Digital expression matrices were generated with the *HTseq* package (*94*) (*python*/v2.7.13) by counting strand-specific UMIs mapping to exonic and intronic regions of each gene. Multi-mapping reads (*i.e.*, those assigned to multiple genes) were resolved by choosing the gene whose 3’ end was closest to the mapped position; reads mapping within 100 bp of the 3’ end of more than one gene were discarded. For downstream analyses, both expected-strand intronic and exonic UMIs were included in the per-gene single-cell expression matrix. Low-quality cells (UMI count < 100, genes detected < 50, or unmatched rate [proportion of reads not mapping to any exon or intron] ≥ 0.4) were filtered out.

##### Doublet removal

To exhaustively identify and remove potential doublets, we applied a two-step procedure. First, we detected doublets directly with Scrublet. To reduce memory and runtime, the dataset was randomly partitioned into four subsets, and the *scrublet*/v0.1 pipeline (*95*) was run on each subset with the following parameters: min_count = 3, min_cells = 3, vscore_percentile = 85, n_pc = 30, expected_doublet_rate = 0.06, sim_doublet_ratio = 2, n_neighbors = 30, and scaling_method = ‘log’. Cells with a doublet score above 0.2 were flagged as putative doublets. Second, we performed two rounds of clustering to identify subclusters enriched for these flagged cells, using *Scanpy*/v.1.6.0 (*96*). Gene counts on sex chromosomes (chrX and chrY) and genes with zero counts across all cells were removed. Counts were normalized by each cell’s total UMI count and scaled to 10^5^, then log-transformed with a pseudocount of 1, and the top 3,000 most highly variable genes were identified. The raw count matrix was then subset to these genes, and normalization (to 10^5^ per cell) and log-transformation were re-applied, followed by scaling of each gene to zero mean and unit variance. Dimensionality was reduced by PCA (30 components), and a neighborhood graph was constructed with scanpy.pp.neighbors (n_neighbors = 50), followed by Louvain clustering (scanpy.tl.louvain, resolution = 1). Each resulting cluster was then subclustered using the same procedure with a smaller neighborhood and higher resolution (n_neighbors = 30, resolution = 3). Subclusters in which more than 15% of cells had been flagged by Scrublet were designated as doublet-derived. Cells were removed if they were either flagged individually by Scrublet or belonged to a doublet-derived subcluster.

The previous two steps were performed automatically, leaving some small clusters driven by doublets difficult to detect. We therefore applied a third step to further identify and remove doublets. It consists of a series of six substeps: (i) Each cell’s expression vector was reduced to retain only protein-coding genes and lncRNAs, with sex chromosomes excluded. (ii) Doublets identified in the previous two steps were removed. (iii) The key difference from the previous step is that here we detected cell partitions using the *partitionCells* function implemented in Monocle3 (*4*), which automatically partitions cells to learn disjoint or parallel trajectories based on concepts from approximate graph abstraction. (iv) Each cell partition was downsampled to 2,500 cells, differentially expressed genes across partitions were computed, and a gene set was assembled by combining the top 200 marker genes for each partition. (v) Cells from each main cell partition were subjected to dimensionality reduction using this set of top partition-specific gene markers. (vi) Subclusters that expressed low levels of the genes identified as differentially expressed in step 4, showed high expression of markers specific to a different partition, and had relatively high doublet scores were labeled as doublet-derived subclusters and removed from the analysis. Overall, 7-14% of cells were removed from each experiment through this procedure.

##### Correction of RT primer cross-contamination

We identified a contamination issue affecting a subset of the RT primer wells used during the sci-RNA-seq3 experiments. Briefly, an unusually high number of cell pairs shared both ligation and PCR barcodes but not RT barcodes. These pairs, which collectively impacted ∼10% of cells, overwhelmingly: (i) derived from a few dozen RT well pairs, (ii) shared cell type, and (iii) had lower UMI counts. We recognized this as the signature of “cell splitting” (i.e., a single cell or nucleus manifesting in the data as two cells because its molecules are split across two distinct RT barcodes), effectively the opposite of “cell doublets”. To resolve this, we merged reads associated with the splitting artifact into single cells, across 54 RT wells forming 23 confusable groups (2-6 wells per group; cross-contamination occurred pairwise within each group), and reprocessed the data from raw reads. Relative to the pre-correction cell × gene matrix, the total number of profiled cells dropped by ∼10% while UMI counts per cell increased by ∼15% (before: n = 1,753,895 cells, median UMI = 790, median genes = 583; after: n = 1,584,848 cells, median UMI = 906, median genes = 654).

##### Integration site mapping

We sought to leverage the sci-RNA-seq3 reads to map the integration sites of DNA Typewriter components. The piggyBac insertions into the genome are flanked by Inverted Terminal Repeats (ITRs). We did a simple grep search of the raw FASTQ files for the ITR sequences and their reverse complements. We then extracted flanking sequences and searched for alignments in the mouse reference genome (mm39) with BLAT. The PEmax and epegRNA-circtape plasmids have differing sequences next to the ITRs, so we could also assess which vector was integrated at which junction. Junction coordinates and the numbers of supporting reads were tallied and are reported in **table S1**.

##### Cell clustering and cell-type annotation

Following doublet removal, we applied additional quality filters to exclude: cells with >85% of reads mapping to exons; cells with log_2_ UMI counts greater than mean + 2 SD or less than mean − 1 SD; cells with a *Scrublet* doublet score > 0.1; cells with more than 5% of reads mapping to ribosomal genes (Ribo%); and cells with more than 5% of reads mapping to mitochondrial genes (Mito%). After filtering, 1,584,848 cells from the E13.5 embryo (embryo #3) were retained, with a median of 906 UMIs and 654 genes detected per cell. Rather than processing the new dataset in isolation, we integrated it with our recently published time-resolved scRNA-seq atlas of mouse development (*5*), 11.4 million nuclei from 74 well-staged embryos sampled every 6 hours from E8 to P0, annotated with 26 major developmental trajectories and 190 distinct cell types. For annotation transfer, we selected 7 timepoints spanning E12.75 to E14.25 from the atlas (1,645,424 cells) and combined them with the embryo #3 data. Because the atlas was originally aligned to mm10, we re-aligned the raw reads to mm39 and re-annotated genes using GENCODE vM37 to match the new dataset. The combined dataset was processed with *Scanpy*/v.1.6.0 (*96*) as follows: (i) genes were restricted to protein-coding and lncRNA biotypes, and counts on sex chromosomes and the mitochondrial chromosome were removed; (ii) UMI counts were normalized by the total count per cell, scaled to 10^4^ counts per cell, and log-transformed with a pseudocount of 1.; (iii) the 2,500 most highly variable genes were selected, and their expression was scaled to zero mean and unit variance; (iv) PCA was performed, and the top 50 components were used to construct a neighborhood graph (n_neighbors = 50); (v) cells were visualized in 2D and 3D via UMAP (min_dist = 0.1). Within this shared PCA space (dims = 50), each new cell was assigned a major trajectory and cell-type label by majority vote among its 20 nearest neighbors in the atlas. Cell type and major trajectory labels were transferred separately.

##### Primary analysis of single-cell circtape data

###### Input sequencing data

Circtape amplicons were generated from the same indexed cDNA as the transcriptome libraries, by the tape-specific two-step PCR described above, in the course of both sci-RNA-seq3 experiments; nuclei of embryo #2 were co-processed with those of embryo #3 in the first of these. All single-cell circtape data analyzed here derive from embryo #3. Because the two embryos’ integration barcodes are disjoint and no closer than 4 mismatches to one another (twice the correction radius applied below), embryo #2 molecules are discarded at barcode correction; such reads account for ∼2.5% of barcode-extractable reads. Libraries were subjected to single-end (NextSeq 2000 P3 200 platform, Read1: 35bp, Index1 10bp, Index2 10bp, Read2: 183bp; NovaSeq X Plus 10B 300 platform, Read1: 35bp, Index1 10bp, Index2 10bp, Read2: 183bp) or paired-end (NextSeq 2000 P2 600 platform, Read1: 250bp, Index1 10bp, Index2 10bp, Read2: 250bp) sequencing. Reads were processed in the first part of the sci-RNA-seq3 pipeline to pull out the three indexes that would identify the cell to match it with its transcriptome; Read1 of the paired-end run retains 215–216 bp of insert once those bases are removed. Each processed read header carries the cell identifier and an 8-bp unique molecular identifier (UMI).

###### *Read merging* (tape/merge.py)

In the paired-end data, R2 is the amplicon read (spanning the integration barcode through the array to the 3’ terminal landmark) and R1 is its mate from the 3′/poly-A end. R1 and R2 mates were merged into a single full-length consensus read by overlap using fastp (*97*) (--merge --correction, with paired-end adapter detection and quality- and length-filtering disabled; minimum overlap 15 bp, allowing ≤8 mismatches and ≤30% mismatched bases within the overlap), which corrects low-quality overlap bases using the higher-quality mate. Because the amplicon read (R2) alone spans the full barcode-array-terminal region at high quality while R1 mates were of substantially lower quality (mean Phred 24, versus 33 for R2), we used a merge-with-fallback scheme: where a pair merged (22.6% of pairs, those in which R1 is clean), the merged read was used; otherwise the amplicon read (R2) was used. Merged reads were reverse-complemented to the amplicon orientation. The single-end run required no merging. Two sequencing runs contribute to the final callset: a single-end run covering cells from both sci-RNA-seq3 experiments, and a paired-end run covering cells from the first experiment only, sequenced at greater read length. Both were called in a single pass, pooling reads by (cell, integration, UMI) so that a molecule sequenced in both runs is counted once. The callset comprises 1,559,419 cells with at least one called tape genotype, 98.4% of the 1,584,848 nuclei that passed transcriptome QC; the remaining 25,429 yielded no genotyped tape and are absent from the matrix.

###### *Reference sets, parsing, and de-noising* (scripts/sc_consensus_par.py, using tape/sc.py with tape/parser.py and tape/denoise.py as above)

The integration-barcode whitelist and insertion vocabulary for embryo #3 were taken directly from the bulk analysis above. Single-cell reads begin at the integration barcode (rather than after a GGCATG flank), so the barcode is taken as the 12-bp tag immediately 5′ of GCTCTGG and corrected to the whitelist as in the bulk pipeline, and the six-monomer array is resolved and validated identically. Reads were tallied per cell, per integration, and per UMI. PCR duplicates were then collapsed to molecules by UMI. A UMI within one mismatch of a more-abundant UMI at the same cell and integration is merged into it, a merged UMI supported by fewer than 2 reads is then discarded (single-read UMIs are enriched for PCR and ambient error), and each molecule was assigned the dominant genotype among its reads. Within each (cell, integration), the per-barcode de-noising steps from the bulk pipeline (error-collapse of single-mismatch neighbors and prefix/suffix chimera removal) were applied to the molecule counts, and any genotype supported by fewer than 2 molecules was discarded. The cross-barcode swap filter is specific to the bulk population and is not applied per cell.

###### *Per-cell consensus* (tape/sc.py + tape/editchain.py)

A single tape genotype was called per cell and integration using an edit-chain model. Because tape transcripts carrying earlier edit states may persist across cell divisions, a molecule with fewer edits that lies on the same ordered edit chain is a prefix (carry-over) of the cell’s current genotype rather than a distinct allele; the consensus therefore descends the most-supported edit chain, folding such prefix molecules onto it, and reports the deepest dominant genotype. Two different edit symbols competing at the same next monomer (which neither a single lineage nor carry-over can produce) define a branch; a locus is flagged as a putative doublet when any such off-path branch carries ≥2 molecules. Branching does not truncate the call (the consensus is always the deepest dominant tip) but it lowers the locus’s dominance, defined as the molecules on the dominant root-to-tip path (the called genotype’s own molecules plus every prefix molecule on that path) divided by all molecules at the locus, equivalently 1 − (branching molecules / all molecules). A locus is emitted only when the called lineage is supported by ≥3 molecules and its dominance is ≥0.9; otherwise it is left missing, on the principle that a wrong call harms the tree while a missing one does not. This yields a cell × integration character matrix of tape genotypes, together with per-cell quality metrics: the number of integrations recovered (n_loci), the number carrying a competing allele (n_doublet_loci), and the mean fraction of molecules supporting the called allele across loci (mean_dominance).

###### *Blastomere routing* (blastomere_route.py)

Before tree building, cells were partitioned into the two daughters of the first cleavage (A and B) from their founder edits directly, rather than letting the neighbor-joining distance resolve that split, which misassigns a few percent of low-coverage cells at the first division, either through shared missingness or an ambient bleed edit at a founder locus. Blastomere-defining alleles were taken at monomer-1 of each integration, the monomer at which the two founder chains diverge, by bipartitioning well-covered cells (≥8 recovered integrations, so that shared missingness could not drive the split) on the leading principal component of their rarity-weighted monomer-1 alleles, split at the threshold that maximizes between-group variance, and retaining alleles present in ≥85% of recovered calls on one side and enriched by ≥0.30 over the other. This gave 20 defining alleles across all 11 integrations. Every cell was then routed by majority vote over these alleles, provided it carried at least one and no more than one allele of the opposite side; the integration carrying that opposite-side allele was treated as ambient bleed and its genotype set to missing in that cell, which affected 58,711 of the routed cells. Cells with no founder allele, a tie, or opposite-side alleles at ≥2 integrations were set aside rather than assigned, and handed off rather than discarded, since their composition is not fate-neutral. Of 1,559,419 cells, 889,849 (57.1%) routed to A and 653,335 (41.9%) to B, 1,543,184 (99.0%) in total; the remaining 16,235 (1.0%) lacked an unambiguous founder assignment and were set aside. The per-cell metrics from consensus calling were then used for a final filter: cells were flagged as passing QC when they carried ≥4 recovered integrations, ≤1 doublet locus, and mean_dominance ≥0.95, retaining 1,296,799 of the 1,543,184 routed cells (84.0%). These divide by tape count into the two inputs to tree building: 663,785 cells with ≥7 of 11 tape genotypes, used for backbone reconstruction, and 633,014 with 4 to 6, reserved for placement onto the dated backbone (see below). At both stages we further required that a cell carry no tape genotype contradicting its blastomere’s founder state, *i.e.* the near-fixed prefix of edits written before the first cleavage. This removed 8,084 (1.2%) of the backbone-eligible and 7,574 (1.2%) of the placement-eligible cells, giving the 655,701-cell backbone and the 625,440 cells placed onto it. All routed cells remain in the released matrices; the tree stages select on the QC flag and the founder-state requirement above.

##### Phylogenetic reconstruction of the backbone tree

We reconstructed the time-calibrated, zygote-rooted, 1,281,141-tip phylogeny in three stages. First, we built a blastomere-resolved backbone tree from 655,701 cells with high-confidence tapes, using neighbor-joining on independently constructed distance matrices for each of the two founding blastomeres (A: 410,925 cells; B: 244,776 cells). Second, we applied molecular clock dating, independently to each subtree, to recover time-calibrated branch lengths. Third, we grafted an additional 625,440 cells onto this timed backbone using a fast sample-placement approach, avoiding full re-inference at this scale.

###### Cell selection and blastomere partitioning

From the blastomere-routed cells that passed in the upstream filtering procedure, we retained those with at least 7 genotyped tapes out of all 11 (663,785 cells, of which 655,701 survived the further filters noted above).

###### Distance matrix

For each blastomere, we constructed a pairwise distance matrix from processed tape sequences, using a distance function adapted from Choi *et al.* (*33*). Let e_i_(z) denote the number of edits that cell *i* acquired in tape *z*, and p_ij_(z) denote the length of the shared edit prefix between cells *i* and *j* in tape *z*. Writing Z_ij_ for the set of tapes recovered in both cells *i* and *j*, we computed the distance as the average, over Z_ij_, of the number of diverging edits:

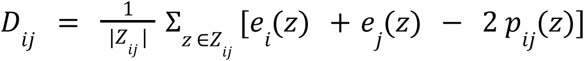

This distance metric essentially counts the number of diverging edits between two cells. We use the average over tape distances to handle missing tapes.

Per blastomere, we added one synthetic cell to the distance matrix that carries only the founder blastomere edits at each tape (**table S2**) and otherwise unedited. This synthetic cell is later used as an outgroup to root each subtree.

###### Neighbor-joining tree inference

We built phylogenies independently for the daughters of the first cell division (blastomeres A, B), from 410,925 (A) and 244,776 (B) cells plus 1 synthetic outgroup cell per blastomere. For each blastomere, we inferred a neighbor joining tree with DecentTree’s RapidNJ (*49*), rooted the tree on its synthetic outgroup, and then dropped that outgroup tip. Negative branch lengths, 6.5% (A) and 4.9% (B) of branches, were corrected with the BAT R package (v2.11.1) tree.zero() (*47*), a Felsenstein–Kuhner correction that zeroes each negative edge while shortening its two child edges by the same total amount, approximately preserving tip-to-tip distances (*48*).

###### Tree dating

Rooted trees were time-scaled by least-squares dating in LSD2 (v2.4.4) (*51*), fixing the root at day 1.5 (the two-cell blastomere founder) and all tips at E13.5, with variance mode -v 1, a 2-hour minimum branch length (0.083333 d), and near-zero input branches collapsed to polytomies at -l 0.01. Because a subsample genealogy cannot carry more coexisting lineages at any developmental time than the whole embryo had cells, we constrained dating with a literature-derived embryo cell-count ceiling (**table S3**), halved to reflect each blastomere’s expected share of the whole embryo. This per-side ceiling was inverted into an order-statistic ladder of per-node minimum-age lower bounds such that the split raising the tree’s coexisting-lineage count to K cannot predate the first day the embryo held K cells. We supplied this ranking to LSD2 as a -d date file, with dating iterated over three passes that re-ranked nodes each time. The two dated per-side trees were finally joined onto a shared day-0 zygote root by a 1.5-day stem, producing a timed backbone tree of 655,701 tips.

##### Phylogenetic placement onto the backbone tree

Reconstructing a tree of >1 million cells is challenging. One option is to first define early splits in the tree and then reconstruct the resulting subtrees independently. But the earliest divisions after the 2-cell stage had the sparsest editing and a cell sorted into the wrong subtree stays there, along with everything descended from it. So we opted for another strategy, where we first infer a backbone tree from a subset of high-quality cells and then place the remaining cells onto the backbone one at a time, scaling linearly.

Cells were added to the backbone tree in two stages. First, a <u>placement</u> stage assigned every unplaced cell to a best ancestor on the basis of shared edits, defining the topology of attachment. Second, a <u>grafting</u> stage converted each placement into a time-scaled branch on the dated backbone tree, preventing us from redating the entire tree.

To place a query cell, we identify an anchor among the tips of the backbone tree and any previously placed query cells. The anchor is the cell with the smallest edit distance to the query, in the same edit distance that built the backbone: the minimum number of edits separating two cells. Because this metric is coarse and frequently produces ties, we first favor candidates sharing the greatest number of tapes with the query and then those with the highest tape recovery.

Exhaustively comparing every query cell with every candidate would be computationally prohibitive. Instead, we use an inverted index of rare edits to retrieve a set of likely candidates (top 500 cells in terms of shared rare edits) and compute the exact edit distance only for these, selecting the closest as the anchor. Because previously placed query cells are also eligible as anchors, closely related query cells naturally form local sublineages rather than attaching independently to the backbone. Query cells with identical genotypes are represented as a single polytomy rather than being arranged into an arbitrary ladder.

A placed cell needs an upstream branch as well as a position. We insert a new node between the anchor and the query, at the point implied by their shared edited prefix, the closest approximation to their real common ancestor. Because tape edits accumulate sequentially, each cell’s own distance to that ancestor is simply the edits it privately acquired past the shared prefix, and their sum defines the total distance between the query and the anchor. We convert the query’s edit distance to time by dividing it by the molecular-clock rate estimated for that blastomere. The new internal node is then placed at that interval before the sampling time, splitting the anchor branch as needed. If this would place the node older than the anchor’s parent, we instead cap it at the parent’s age. Both the anchor and query remain sampled at the observed time. Thus, placing a query rescales a single branch without altering the dates of the existing tree.

##### Internal node support

Resampling-based bootstrap support is computationally infeasible at this scale, as rebuilding a 655,701-cell tree hundreds of times is prohibitively expensive. Instead, we report a per-branch edit-count support statistic that conditions on the inferred topology and measures how many independent editing events across tapes support each branch, rather than how often that branch recurs across bootstrap replicates. This statistic is conceptually related to the branch parsimony score used to assess support in large viral phylogenies, including UShER (*98*), but addresses a different question. Whereas the UShER score quantifies confidence in the placement of a newly added sample, our statistic quantifies the support for a branch already present in a fixed tree.

Because edits are acquired irreversibly and in a fixed write order within each recording tape, the topology and tip genotypes uniquely determine the most parsimonious ancestral tape state at every internal node. If two descendant lineages differ at a monomer, that monomer and all downstream monomers are inferred to be unedited in their common ancestor. Each edit is then counted once, on the branch where it first appears.

Summing these counts across all 66 recording monomers (6 monomers on each of 11 independent tapes) gives a branch’s edit-count support: the number of independent loci whose reconstructed edit history supports that branch. This rule is deliberately conservative and is not symmetric in clade size, *e.g.* early nodes with many descendants lose edits to a single discordant call, while recent nodes with a handful of descendants are less affected simply because they have fewer opportunities for a cell with a discordant call. Both the support statistic and the per-branch rates derived from it may therefore be biased downward at early nodes. Descendants lacking a genotype call at a given tape are skipped rather than treated as disagreements; a descendant that is genotyped but whose edit chain is truncated by dropout does, however, shorten the inferred prefix.

##### Clonal dominance

###### Founder definition

We cut the time-scaled tree at day T_0_=E7.0; each ancestral branch in the tree at that time defines one founder and its tip count at E13.5 is that founder’s clone size.

###### Dominance statistics

We quantified skew in the founder clone size distribution in two ways: (i) the number of founders required to reach 50% of all cells in the embryo, and (ii) the Gini index of clone sizes across all founders computed with ineq::Gini from the R package ineq (v0.2-13) (*99*, *100*). The Gini index summarises inequality among founder contributions into one number (0 = all founders contribute equally, 1 = one founder produces all cells). To determine when clonal dominance emerged, we held the set of T_0_-defined founders fixed and recomputed the Gini index at 50 successive developmental time points by recounting the descendants of each founder present in the reconstructed cell phylogeny at each time point.

###### Null model

Observed skew was compared to a neutral-drift null in which N_0_ founders (matching the T_0_ founder count) each undergo an independent Yule (pure-birth) process (*67*) at a shared rate λ for the same duration ΔT (T_0_→E13.5). λ was fit so the null’s expected total population matched the observed cell count. The required founders for 50% statistic were tested via 10,000 replicate draws from this null; Gini was tested the same way. The null was refit independently for each blastomere.

###### Tree basis

Clonal-dominance panels in **Fig. 4A-C,E** were computed on the full placement-augmented tree (3,849 founders at T₀ = E7.0). The backbone tree, cut at the same T_0_ (3,848 founders), was used for **Fig. 4D,F** and **Fig. S14A** for analysing the Gini index over time. Further, it served as replication: **fig. S13A-C** repeats **Fig. 4A-C**; **fig. S14E** repeats **Fig. 4E**,;**fig. S14F** compares per-trajectory Gini indices between the two trees; and **fig. S14A** varies the apportionment of the lineage-count ceiling. Blastomere-split panels for **fig. S13D-F** use the A and B subtrees of the full tree and for **fig. S14B** the subtrees of the backbone tree.

##### Tree sibling analyses

Tree siblings were defined as pairs of tips sharing an immediate parent. This included: (i) cherries, in which the parent had exactly two tip children, and (ii) terminal polytomies, in which the parent had ≥3 tip children. Within each terminal sibling group all unordered pairs were enumerated, so a group of n tips contributes n(n−1)/2 pairs. Tips whose only siblings are internal nodes (367,636 of 1,281,141; 28.7%) contribute no pairs, to the observed data and to the null alike. Tree topology was extracted from the final placement tree (1,281,141 tips) using ape (v5.7.1), yielding 870,785 sibling pairs across 406,906 terminal sibling groups -- 349,595 cherries, and 57,311 polytomies (up to 157 tips) contributing 521,190 pairs -- with the A and B blastomere subtrees analyzed independently.

Enrichment was computed on the combined tree, with the independently reconstructed blastomere A and B subtrees analysed separately as replication. For each of the 24 major trajectories and the 99 annotated cell types present at ≥0.01% in both blastomere subtrees, we counted sibling pairs in which both cells shared the same annotation. To generate a null distribution, major-trajectory or cell-type labels were permuted across annotated tips with the tree held fixed (100 permutations), which preserves both the sibling structure and each label’s abundance. Sibling enrichment was calculated as (observed+1)/(mean permuted+1), with a pseudocount of 1 added to stabilize estimates for rare annotations. Reproducibility between blastomeres was assessed by Spearman’s ρ of enrichment values across fates.

##### Heterotypic sibling analyses

Within the same sibling-pair set, we asked which progenitor→post-mitotic pairings recur among tree siblings above chance, taking each as a captured terminal differentiation division. Of the 138 cell types, 27 were classified as post-mitotic (**table S6**), and a sibling pair was heterotypic when exactly one of its members was post-mitotic. Analysis was restricted to pairs whose shared ancestor dated ≥E12.5 and whose members differed at no more than one discordant tape monomer among the monomers called in both cells. Both criteria are independent of cell identity, so the eligible pair set is identical for observed and permuted labels. Each post-mitotic cell was counted at most once, credited to its nearest qualifying sibling (fewest discordant monomers; ties broken by most shared edits), so that a terminal polytomy cannot contribute more candidate divisions than it has cells. Pairings supported by fewer than three such cells were not tested, leaving 640. The null permuted cell-type labels across annotated tips within each blastomere with the tree held fixed, recomputing the whole procedure for each of 500 permutations. Fold enrichment was (observed+1)/(expected+1), p was the upper tail of a Poisson with mean (expected+1), or of a negative binomial matched to the permuted mean and variance where the null was overdispersed (24% of tested pairings), and a pairing was called a captured division at Benjamini–Hochberg FDR < 1% with fold ≥3, testing over the whole tree and treating the two blastomeres as validation rather than as a gate.

##### Clade co-occurrence

Maximal clades were identified independently on the A and B blastomere subtrees of the full placed tree. For a given size cap K, we traversed each subtree and, at each internal node, evaluated the number of tips in its subtree counted over that blastomere’s leaves alone: if this count fell between 3 and K inclusive while the count for its parent’s subtree exceeded K, the node’s subtree was called a maximal clade. This yields a set of non-overlapping clades covering the largest possible groupings not exceeding K tips (terminal groups of fewer than 3 tips are not called, so the clades cover rather than exactly partition each subtree). Clades were identified separately at K=15 (small, tightly related family groups; 82,749 (A) and 57,969 (B) clades; median ancestor age E11.3 (A) and E11.2 (B)) and K=200 (large, loosely related family groups; 10,047 (A) and 6,769 (B) clades; median ancestor age E8.9 (A) and E8.8 (B)).

For each pair among the 88 cell types represented by ≥100 cells in both blastomere subtrees, we counted the number of maximal clades in which both cell types occurred (co-occurrence count), separately for each blastomere and each value of K. The null is tip-label permutation within each blastomere with the tree and clade partition held fixed. Enrichment for each pair was calculated as log_2_((observed+1)/(expected+1)), the pseudocount stabilising pairs with few co-occurrences. A pair was called significant when observed co-occurrence exceeded the null expectation by ≥3 Poisson standard deviations on at least 5 co-occurring clades; for the recent-versus-deep classification this was required in both blastomeres independently.

Hierarchical clustering of the K=15 log_2_-enrichment matrix was performed using Ward linkage on Euclidean distance between matrix rows. When making the heatmap, the clustering order was manually adjusted so that cell types sharing a developmental origin appeared contiguously. Because the rotation of any internal node of a dendrogram is arbitrary, the raw order places a small number of cell types away from others of the same lineage. In the old order, the retinal trio (rows 14–16) was split from the rest of the anterior neuroectoderm by the three endoderm rows (Hepatocytes, Lung progenitor, Midgut/Hindgut, rows 17–19); moving endoderm down to rows 31–33 rejoins the retinal trio with the rest of the anterior neuroectoderm as one contiguous block. Separately, Cranial motor neurons moved from row 79 (inside posterior neuroectoderm) to row 75, landing at the boundary between neural crest/PNS and posterior neuroectoderm. These adjustments affect display order only; the same order is applied to rows and columns, and the log_2_-enrichment values, the permutation-based null, and the clustering itself are unchanged.

##### Timed fate couplings

For each time point from E7.0 to E13.25 in 0.25-day increments, we defined time-slice clades on the full placed and dated tree. Each branch intersecting time *t* defined one clade consisting of all sampled descendant tips. Time-slice clades were identified independently for the A and B blastomere subtrees as branches whose parent node predated *t* and whose child node was dated at or after *t*. Clades containing fewer than three cells of the blastomere under analysis were excluded. The number of lineages present rises from 3,849 at E7.0 to 309,584 at E11.0.

For each of the 88 cell types (≥100 cells in each blastomere subtree), and for every cell-type pair, we counted the number of time-slice clades containing at least one cell of each type. The tip-label-permutation expectation was evaluated exactly in closed form, holding tree topology and time-slice clades fixed; the procedure involves no sampling and is deterministic. A cell-type pair was considered significantly co-enriched at a given time slice by the same criterion used for clade co-occurrence, applied independently in both blastomere subtrees. Coupling depth was defined as the earliest time point at which a cell-type pair first satisfied these criteria. Sensitivity is lowest at the deepest slices, where a tail of very large clades is expected under a tip-label permutation to contain nearly every cell type, leaving little contrast to detect; consistent with this, no pair was co-enriched before E8.25. Reproducibility of coupling depths between blastomeres was assessed using Spearman’s ρ and the median absolute difference between blastomere-specific estimates.

##### Hierarchy construction

A cell-type hierarchy was constructed from the matrix of pairwise coupling depths using average-linkage agglomerative clustering (scipy v1.13, scipy.cluster.hierarchy.linkage, method=average), restricting the analysis to the 75 cell types with a defined coupling depth to at least one partner. Pairs that never couple were assigned a censoring value of E13.5, and dendrogram branch points were labelled with their average-linkage merge height, which is a coupling depth in embryonic days rather than an abstract merge distance. Branch points whose height exceeded the last time slice (E13.25) can only be carried by the censoring value and were left undated (23 of 74).

##### Heuristic inference of the transcriptional states and cell type annotations of ancestral nodes

For ancestral state imputation, we repeated the joint preprocessing described above using the embryo #3 dataset together with 5.2 million atlas profiles spanning E8.5-E13.5 (*5*). The combined dataset was normalized, scaled, and subjected to principal component analysis in Scanpy (v1.6.0), retaining the first 50 principal components.

##### Ancestral state imputation

Starting from the observed E13.5 tips, we traversed the full placement tree from tips to the root using a post-order traversal. Each internal node was assigned a principal-component profile equal to the unweighted mean of its children’s profiles, with tip nodes contributing their observed PC coordinates and internal nodes their previously computed mean profiles. Ancestral nodes were linked to atlas cells by mutual nearest neighbors (MNN), proceeding in successive 6-hour windows from E13.5 to E8.5. Within each window, an ancestral node and atlas cell were linked if each ranked among the other’s 200 nearest neighbors in the 50-dimensional PC space (Euclidean distance). Nodes without a MNN match were left unassigned (12%). Each matched node received a cell-type label by majority vote across its linked atlas neighbors. Each ancestor was then assigned an updated transcriptional profile -- the mean PC profile of its MNNs.

###### Traceback path

For each E13.5 tip we walked root-ward and recorded the imputed ancestral labels, skipping uncalled nodes and collapsing contiguous runs of the same label, giving one sequence of successive distinct states per lineage, read oldest to youngest and ending on the tip’s own observed cell type. Paths were required to be heterotypic (≥2 distinct states) and non-recursive (no state appearing twice). The latter removes lineages in which a state recurs after having been left, which reflects oscillation of the imputation between two co-resident labels rather than differentiation. Of 1,281,141 tips, 129,204 (10.1%) had no heterotypic ancestry and 595,304 (46.5%) were recurrent, leaving 556,633 traced on 132,536 distinct paths across 136 of the 138 cell types.

A path was additionally defined as qualifying if: (i) it was followed by at least 1% of that cell type’s traced cells and at least 5 cells, and (ii) every step was directionally asymmetric. Transitions were counted over nearest-called-ancestor pairs: for each node carrying a call whose nearest called ancestor carried a different call, one transition was recorded from the ancestor’s to the node’s label, with tips included. A step A→B was retained only if that pair had at least 50 transitions and asymmetry (n_AB − n_BA)/(n_AB + n_BA) ≥ 0.2, *i.e.* the observed direction favored at least 60:40. Both criteria use only the phylogeny and the imputed labels. This gave 275 qualifying paths in 82 cell types (117,704 cells; **table S8**). For each we recorded the number of independent origins, defined as distinct tree nodes at which its earliest state occurs, and whether it was recovered in both blastomere-derived halves of the embryo, defined by the two largest clades descending from the root.

###### Traceback evaluation

Qualifying paths were scored against the curated cell-type relationship graph of Qiu *et al.* (*5*), reconstructed from their Supplementary Tables 20 and 22; merging states that share a name and removing self-edges gives 262 states and 474 undirected edges. Our labels were mapped onto these states by normalised string matching with four manual assignments. The graph was used only to evaluate paths, never to select them.

For each step we computed the shortest-path distance d between its two states. d = 1 indicates direct adjacency; we also report d ≤ 2, which permits one intervening state, since paths retain only called nodes and collapse contiguous repeats and a single step may therefore legitimately span more than one curated edge. The 275 paths comprise 362 steps; both endpoints mapped into the graph for 360 of them; the two exceptions both involving Spinal cord ventral progenitors, which has no counterpart among the curated states. Significance was assessed by permuting the 102 mapped cell-type labels appearing in the qualifying paths among themselves and recomputing every step distance (10^6^ permutations; empirical P = (1 + k)/(1 + N) for k permutations reaching the observed value); the observed values exceeded every draw. Directional agreement was assessed only for curated-adjacent steps: because the graph is undirected, orientation was taken from each state’s shortest-path depth from the “Oocyte” node, with a step counted as concordant if the ancestral state is strictly shallower; steps whose states lie at equal depth cannot be oriented and were excluded. Paths were additionally classified by whether all of their states share a germ-layer assignment.

## Supplementary Text

### Note S1. Preliminary characterization of four additional DNA Typewriter embryos

To assess whether certain patterns observed in embryo #3 (*e.g.* the 2-cell burst and subsequent slowdown/recovery, recording efficiency, as well as the fact that PNI-mediated generation of DNA Typewriter embryos worked at all) are reproducible or idiosyncratic, we performed additional PNI sessions with three modifications: (i) the molar ratio of the epegRNA::tape-BC::circtape to PEmax construct was changed from 2:1 to 1:1; (ii) extraembryonic tissues were retained rather than discarded; and (iii) the gDNA input for bulk amplicon sequencing was increased from 5 to 50 ng. Of 8 embryos recovered from 4 foster mothers, 4 exhibited editing; these were harvested at E14.5, E18.5 (n = 2) and E19.5. Because these embryos are staged later than embryo #3, substantially more nuclei will need to be profiled to achieve an equivalently dense cellular phylogeny. However, bulk amplicon sequencing of their tapes is sufficient to ask some questions.

First, are these embryos closer to embryo #2 (poorly edited) or #3 (well edited)? By integration number, editing rate and tape genotype complexity, each of the four new embryos is equivalent or superior to embryo #3 (**fig. S4**). At a minimum, this establishes that embryo #3 was not a one-off, and that PNI followed by piggyBac-mediated integration is a robust means of generating DNA Typewriter mouse embryos at stages ranging from E13.5 to E19.5.

Second, to ask when recording begins, we inferred the stage of the earliest edits at each of 85 integrations across the five well-edited embryos (the four new embryos, together with embryo #3). Walking each array 5′→3′ over cell-count-weighted tape genotypes, we required at every monomer that the classes considered jointly account for ≥95% of genome equivalents, that each account for ≥10% individually, and that ≥50 genome equivalents remain. A monomer at which a single symbol met this bar was scored as a 1-cell (1C) edit (one write event ancestral to every recovered copy) and absorbed before advancing; a monomer at which the top two symbols only jointly met it was scored as a 2-cell (2C) split, with each half then examined independently. Thirty-two integrations (38%), distributed across four of five embryos, carry edits inferred to have occurred at the 1-cell (1C) stage. Twenty-five (29%), distributed across all five embryos, carry edits inferred to have occurred at the 2-cell (2C) stage. In fact, of the five, embryo #3 is the one in which we detect no 1C edits (even embryo #2, though excluded from this analysis owing to its weak overall editing, carries 6 inferred 1C edits and no 2C split). Extending the same criteria to larger class sets, only 2 of 85 integrations (2%) support three classes at a single monomer and none support four; applying the two-way rule within each half of a 2C split identifies a further split in one half of 4 integrations and in both halves of none. Recording at the 4-cell stage is therefore detectable but uncommon by this criterion (which may miss markedly unequal 4-cell stage divisions due to the ≥10% criterion). Consistent with this, manual review of single-cell genotypes from embryo #3 indicates a markedly unequal division of blastomere A (**Fig. 2F**), into daughters marked by three and by one concordant independent edits and contributing ∼40% and ∼7% of the embryo, while no corresponding division of blastomere B is marked. This structure is not recovered by the criterion above, which operates on one integration at a time and therefore cannot aggregate concordant edits at different integrations, nor by neighbor-joining, which separates this bipartition poorly, as it does the first cleavage, which is the reason blastomere assignment was performed explicitly from founder edits rather than inferred. The falloff across the first two cleavages in embryo #3 is nonetheless steep, as the founder states of blastomeres A and B carry 23 and 19 edits respectively, against 4 edits distributed across the four second-cleavage branches.

Two conclusions follow. First, the burst-then-slowdown pattern we report for embryo #3 is reproducible, in that editing is concentrated in the 1C and 2C cleavages but uncommon by 4C in every embryo examined. Second, and newly, the initial burst typically spans both the 1C and 2C stages; embryo #3 is unusual in that its earliest edits appear confined to 2C, which is what permits the two daughters of the first cleavage to be distinguished so cleanly. We have not yet profiled these embryos at single-cell resolution and so cannot date the recovery of recording in them, but that it recovers is well supported by the extent of editing and the rarefaction behavior of their tape genotypes (**fig. S4**).

### Note S2. Evaluation of the frequency and nature of unexpected prime editing events

We sought to assess the frequency and characteristics of *in vivo* prime editing errors at sequenced tapes. A challenge to implementing this assessment is that our production pipeline for tape analysis from single-cell data parses each read into per-monomer editing events, then takes consensus across the reads sharing a UMI, and again across the UMIs recovered for a given integration in a given cell. Because a read carrying an unexpected event fails at the parsing step, it is discarded before reaching either consensus. We therefore built an alternative pipeline that keeps both consensus levels but inverts their order with respect to parsing, *i.e.* taking consensus across the reads of each UMI to give one sequence per molecule, then across the molecules of each integration in each cell to give one tape sequence for that locus, and parsing only at the end. Because each integration in a cell is a single locus carrying a single array, all of its molecules should report the same sequence. Requiring that removes sequencing and amplification errors while preserving whatever the template carries, expected or not.

We ran this alternative pipeline on embryo #3 tape sequences, requiring each UMI to be supported by ≥3 reads, each locus to be supported by ≥3 UMIs, and ≥80% of reads and of UMIs to agree on the architecture of the tape array. A locus whose molecules disagreed yielded no call rather than a majority call, as our aim here is to characterize the distribution of bona fide prime editing events, intended and unintended.

Of 8,057,501 consensus tape sequences from 1,546,765 cells meeting these criteria, 7,676,636 (95.3%) fell into the category of “expected”, meaning that every insertion belonged to the expanded vocabulary set or the site was unwritten, and that all edits occurred sequentially. The main variation within this class is that the arrays were not all six units long. Among the 7,573,214 expected calls in which the 3’ terminal monomer was located, array length was distributed as follows:

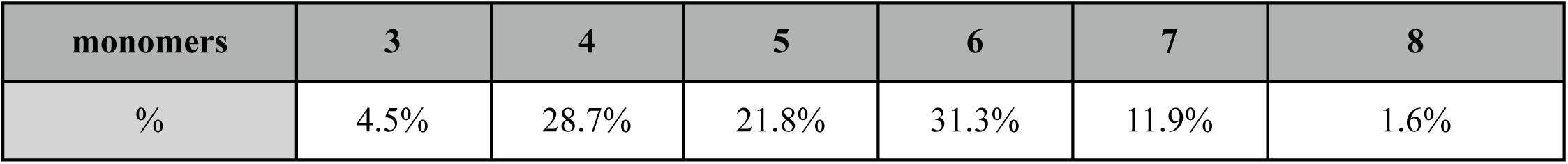

Array length therefore varies substantially between loci. This variation is not simply amplification noise, as even restricting to loci at which every read of every UMI reports the same monomer count leaves the distribution essentially unchanged (six units in ∼32% of loci in both cases), which is not what slippage during library preparation would produce. We count these “non-six-monomer” consensus sequences among those our pipeline would admit, because the production pipeline does not discard them. However, more explicitly representing the variably sized tapes is a clear axis of potential improvement for future iterations of the tape genotyping pipeline.

Turning to the 4.8% of consensus sequences carrying at least one unexpected feature, these break down into the following nine categories:

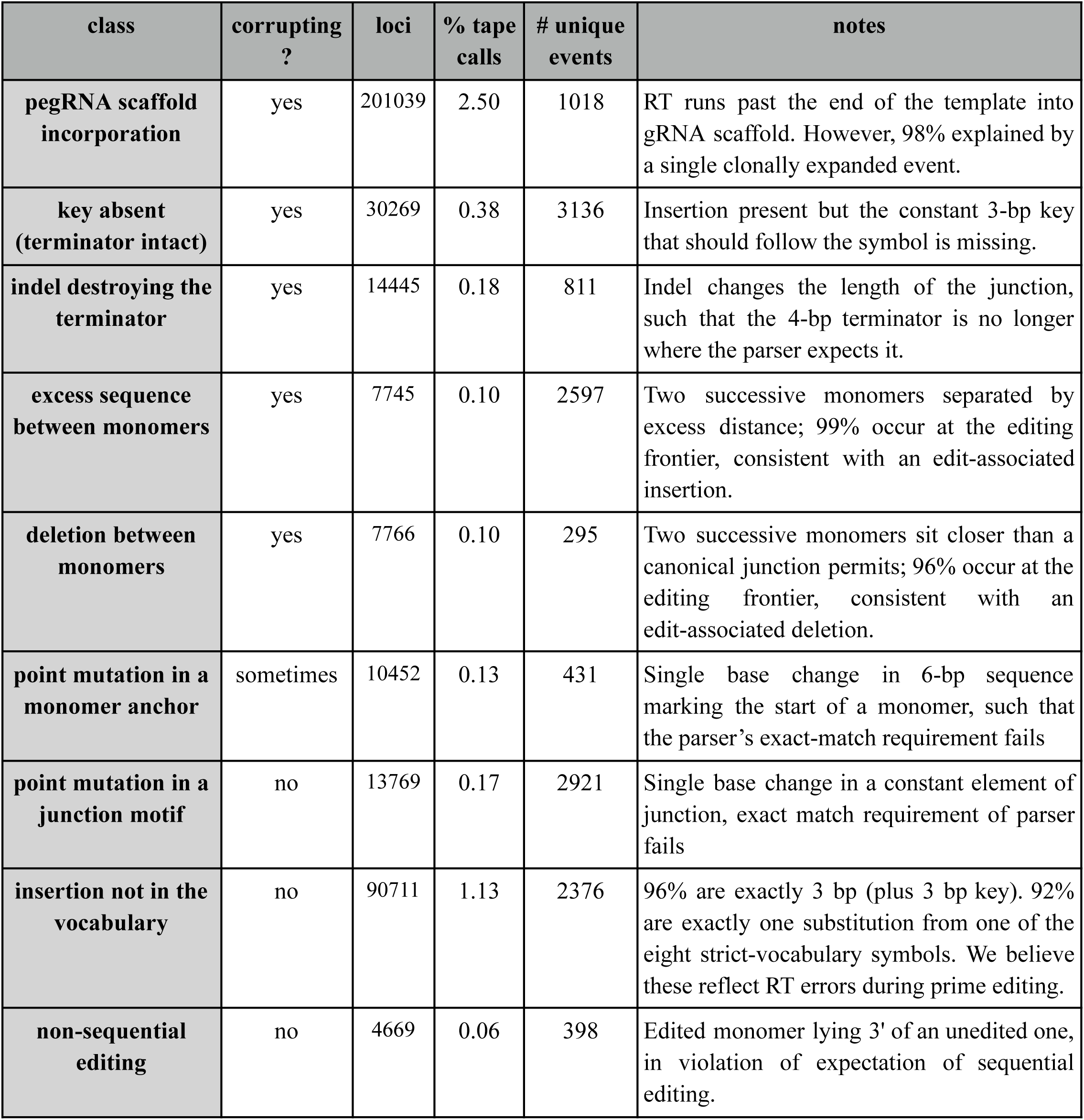

| class | corrupting ? | loci | % tape calls | # unique events | notes |
| --- | --- | --- | --- | --- | --- |
| **pegRNA scaffold incorporation** | yes | 201039 | 2.50 | 1018 | RT runs past the end of the template into gRNA scaffold. However, 98% explained by a single clonally expanded event. |
| **key absent (terminator intact)** | yes | 30269 | 0.38 | 3136 | Insertion present but the constant 3-bp key that should follow the symbol is missing. |
| **indel destroying the terminator** | yes | 14445 | 0.18 | 811 | Indel changes the length of the junction, such that the 4-bp terminator is no longer where the parser expects it. |
| **excess sequence between monomers** | yes | 7745 | 0.10 | 2597 | Two successive monomers separated by excess distance; 99% occur at the editing frontier, consistent with an edit-associated insertion. |
| **deletion between monomers** | yes | 7766 | 0.10 | 295 | Two successive monomers sit closer than a canonical junction permits; 96% occur at the editing frontier, consistent with an edit-associated deletion. |
| **point mutation in a monomer anchor** | sometimes | 10452 | 0.13 | 431 | Single base change in 6-bp sequence marking the start of a monomer, such that the parser's exact-match requirement fails |
| **point mutation in a junction motif** | no | 13769 | 0.17 | 2921 | Single base change in a constant element of junction, exact match requirement of parser fails |
| **insertion not in the vocabulary** | no | 90711 | 1.13 | 2376 | 96% are exactly 3 bp (plus 3 bp key). 92% are exactly one substitution from one of the eight strict-vocabulary symbols. We believe these reflect RT errors during prime editing. |
| **non-sequential editing** | no | 4669 | 0.06 | 398 | Edited monomer lying 3' of an unedited one, in violation of expectation of sequential editing. |

Notes on each error class are provided in the final column. Corrupting classes are those that corrupt the typehead (the target site for subsequent editing) such that further edits of the tape are not possible. The vast majority of instances of the same unexpected event occurring at the same tape integration correspond to a unique event that was propagated to descendant cells. As such, the 380,865 unexpected loci delineated above collapse into only 13,983 unique events. For example, although pegRNA scaffold incorporation seems to be the most common class of unexpected events in terms of how many consensus tapes it impacts (n = 201,039), 98% of these correspond to a single event at monomer-2 of tape “TGACTAAAGCGG” which occurred early in embryo #3 and then clonally expanded. Thus, the most common class of prime editing error in the above table is actually “key absent (terminator intact)”, with 3,136 unique events.

Looking forward, the explicit recognition and appropriate treatment of these error classes, rather than discarding them, represents an opportunity for future improvement for the tape genotyping pipeline. We also note that some additional classes of error events may be undetectable, *e.g.* events that result in deletions or insertions of sufficient size that the workflow for bulk amplicon sequencing renders them undetectable at either the PCR or sequencing stage.

### Note S3. A blood-specific recording slowdown of uncertain origin

Two violations of a strict molecular clock are already apparent: editing slows sharply after the first cleavage and then partially recovers around gastrulation, and the nominally constant recording rate falls as 6-monomer tapes exhaust. However, a third concern, addressed by neither the ceiling nor the timing correction, is that prime editing rate may itself vary across cell types, *e.g.* through dependencies on proliferative state or on the enzymes and nucleotides that prime editing requires (*54–57*).

To assess this, we calculated a per-cell, per-day editing rate for each abundant cell type. The resulting cell type-specific editing rates, in this embryo, fell within a tight 1.6-fold range and were consistent between blastomeres (Spearman’s ρ = 0.92, *p* < 10^-42^). However, the most slowly editing cell types were overwhelmingly of hematopoietic origin (**fig. S9A**). Because these cells’ ancestors presumably edited at typical progenitor rates before adopting a hematopoietic identity, the actual blood-specific slowdown in *in vivo* prime editing rates is likely stronger than these cumulative estimates suggest.

Two explanations are consistent with the data. First, PEmax expression was significantly lower in hematopoietic cell types (**fig. S9B**), suggesting lineage-specific silencing at PEmax integration site(s). Second, *Samhd1* expression was significantly higher (**fig. S9C**), in line with its role in limiting the dNTP pool available to the reverse transcriptase (*57*). However, these patterns are confounded with hematopoietic identity and do not hold up when blood and non-blood cell types are analyzed separately (**fig. S9D-E**). As such, we cannot confidently attribute the slowdown to either, nor can we rule out other blood-specific differences.

Timing *in vivo* lineage trees is therefore more involved than assuming a strict molecular clock (*50*). Our dates should be read as approximations rather than exact ages, most reliable where cell-number estimates anchor them and where recording is active, and least reliable at the earliest divisions, toward the tips, and in hematopoietic cell types. More faithful timing will require models representing cell type-specific rate dynamics directly, particularly as recording extends into later development, which brings both a greater diversity of cell types and a growing fraction of post-mitotic cells.

### Note S4. Robustness of tip-level and clade-level analyses to tree and null

The analyses of tree siblings, heterotypic sibling pairs, and clade co-occurrence rest on the terminal structure of the placement-augmented tree, which our placement heuristic could in principle manufacture, and are evaluated against a global tip-label permutation, which is a permissive null. We therefore repeated them under two alternatives.

#### Backbone tree

On the backbone tree alone, 41% of tree-sibling pairs shared a cell type label, a 7.9-fold enrichment over a permuted null, compared with 51% and 9.4-fold on the placement-augmented tree; enrichments were highly correlated across fates between the two trees (Spearman’’s ρ = 0.94 for cell types, 0.88 for major trajectories). For heterotypic pairings, power is much lower on the backbone (9,405 post-mitotic cells across 219 testable pairings, versus 82,300 across 640 on the augmented tree), but 31 of the 37 significant pairings remained testable, and 24 of those remained significant; the 7 that did not were supported by only 3-7 post-mitotic cells. Off-diagonal maximal clade co-occurrence enrichments agreed closely between trees (Spearman’s ρ = 0.83-0.92 across K=15 and K=200 for both blastomeres). Timed coupling depths were systematically deeper on the backbone (median shift −0.75 days), which may reflect the sensitivity of this heuristic to sampling depth and the poorer reliability of dates at the branchpoints of placed cells, but their rank order was preserved (Spearman’s ρ = 0.86, Kendall’s τ = 0.69).

#### Within-clone null

Replacing global tip-label permutation with a null that permutes labels only among cells descended from the same E7.0 lineage (3,848 clones, median size 32, mean 170) is considerably less permissive, and less arbitrary than permuting within our own major trajectory groupings. Against it, sibling cell-type concordance falls from 7.9- to 2.8-fold, but remains above 1.0 for all 88 cell types with at least five same-type sibling pairs, and cell types strongly enriched under the global null remain so under the within-clone null (Spearman’s ρ = 0.45). Of 86 dated cell-type couplings, 55 survive, with systematically shallower dates (median shift +0.75 days) but consistent rank order (Spearman’s ρ = 0.77).

We conclude that sibling enrichments and the ordering of cell-type couplings are robust to both tree and null, while the absolute dates of couplings are sensitive to sampling depth and should not be overinterpreted.

## Supplementary Figures

**Figure S1.**
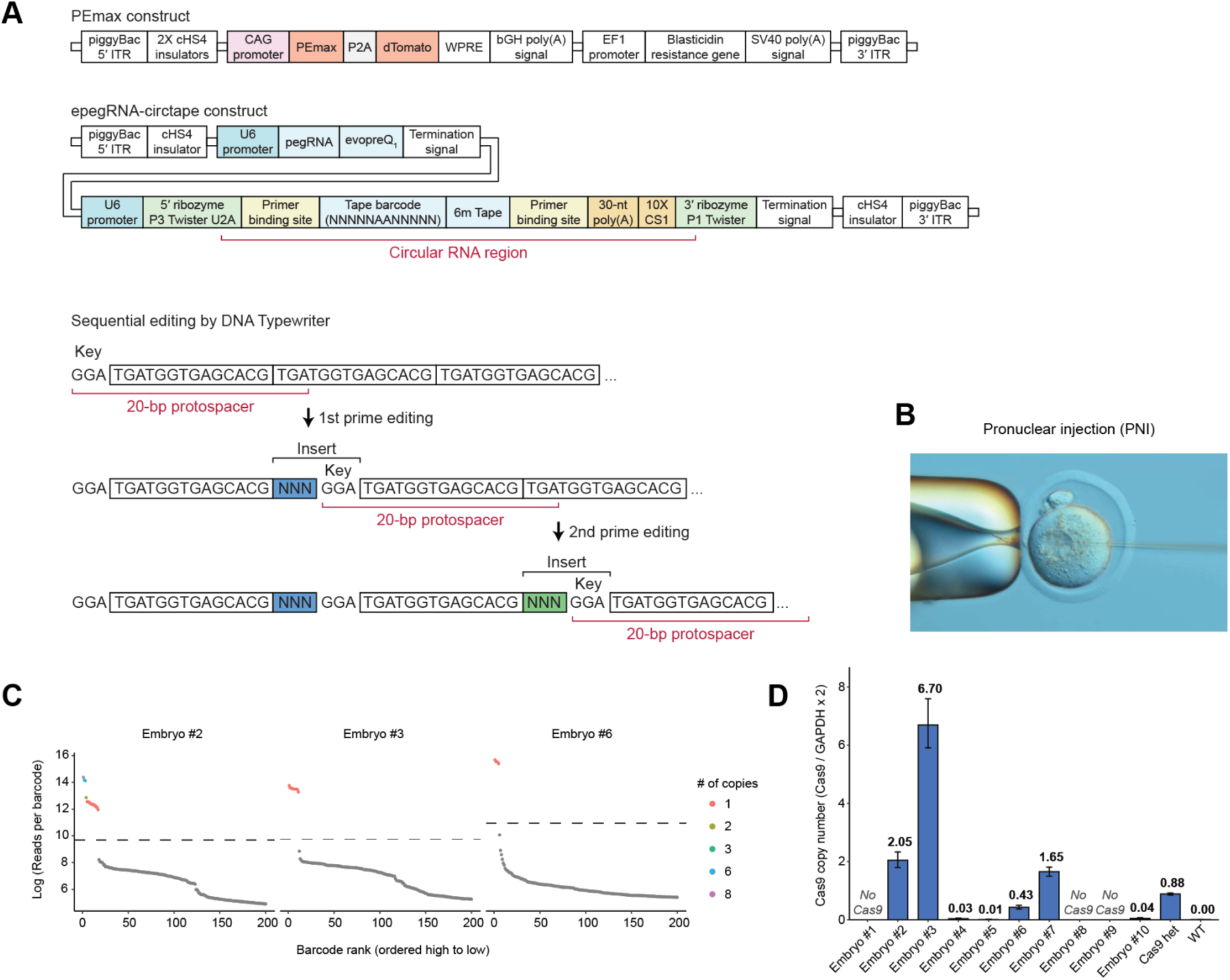
Characterization of epegRNA::tape-BC::circtape and PEmax integrations across E13.5 embryos. **(A)** Schematics of PEmax and epegRNA::tape-BC::circtape constructs (top) and the sequential editing process mediated by the DNA Typewriter system (bottom). CS1, reverse complement of Capture Sequence 1 from the 10x Genomics platform. **(B)** Image of a single-cell zygote during PNI. A finely pulled glass needle was inserted into a pronucleus to deliver a mixture of mRNA and plasmid DNA. Imaged with a 40× DIC objective on a Nikon Ti2A microscope with a Nikon D1000 camera. **(C)** Tape-BC read-count rank-abundance, shown separately for embryos #2, #3 and #6. Reads were first aggregated by correcting each barcode to its nearest whitelist member within 2 mismatches, collapsing one- and two-base error shadows into their parent. Barcodes are ranked by read count (x-axis) against reads per barcode (log scale, y-axis); each point is one barcode, colored by estimated genomic copy number. For embryo #2, some of its 17 tape-BCs appear to be multi-copy, consistent with piggyBac excision and re-integration, which we also observed in high-multiplicity of infection cell culture (*40*). The dashed line marks the whitelist cutoff separating called integrations (left) from background sequences (grey, right). **(D)** PEmax copy number in each E13.5 embryo was assessed by duplex ddPCR (Cas9, FAM; GAPDH, reference), calculated as (Cas9 / GAPDH) × 2. The Cas9 probe targets a region that is common to Cas9, PE2 and PEmax. Embryo #3 carried the highest estimated copy number of PEmax (measured 6.70, rounded to 7), followed by embryo #2 (measured 2.05, rounded to 2). Embryos #1, #8, and #9 yielded zero positive Cas9 droplets (“No Cas9”). Genomic DNA from a Cas9/+ mouse served as a positive control (measured 0.88, rounded to 1), and from a wildtype B6 mouse as a negative control (measured 0.00). Bars show the per-well point estimate; error bars are 95% Poisson confidence intervals propagated through the ratio.

**Figure S2.**
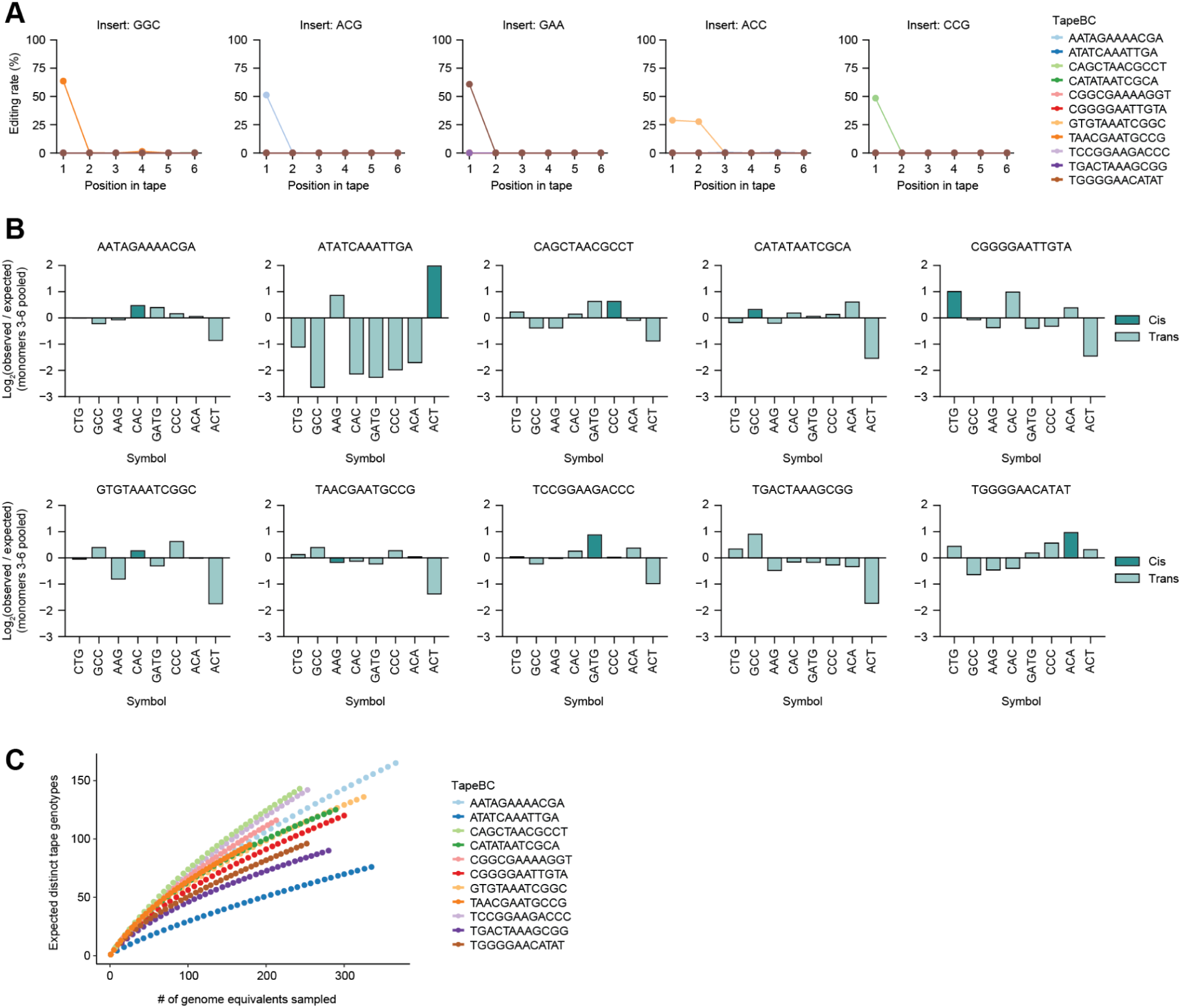
Patterns and diversity of tape genotypes in embryo #3. **(A)** Editing rates of symbols that are abundant despite <u>not</u> being encoded by integrated epegRNAs, across integrations and positions. Of the 13 symbols present at ≥0.5% frequency at any monomer position, the 5 symbols present at ≥0.5% frequency only at monomer-1 and/or monomer-2 are included in this analysis. We interpret their high, integration-specific frequencies at 5’ positions as corresponding to singular, early events. **(B)** Frequencies of symbols written by epegRNAs encoded in *cis* or *trans.* The analysis was restricted to 10 circtapes whose *cis*-encoded epegRNA was identified from bulk amplicon sequencing of epegRNA::tape-BC::circtape integrations, 8 symbols encoded by integrated epegRNAs, and monomers 3-6 to exclude 32 of the 42 founder edits established at the 2-cell stage (**table S2**). From the cell × tape matrix, observed rates for a given integration were calculated as the number of edited monomers bearing a given symbol divided by the total number of edited monomers. Expected rates were calculated in the same way, but across all integrations. **(C)** Rarefaction analysis of recorded diversity in embryo #3. For each of the 11 tape-BC integrations (colored by tape-BC), the expected number of distinct tape genotypes (y-axis) is plotted as a function of the estimated number of genome equivalents sampled (x-axis). The diversity observed for each of the eleven tape integrations lies on the steeply rising portion of its curve, with a terminal slope far from zero, indicating that recorded diversity is far from saturated and is limited by the material sampled rather than by recorder capacity. Of note, the ∼5 ng of genomic DNA used as input to the amplicon sequencing of embryo #3 tape-BC here corresponds to only ∼850 genome equivalents, assuming ∼6 pg of genomic DNA per cell, and likely fewer, as UV quantitation tends to overestimate DNA mass (*101*).

**Figure S3.**
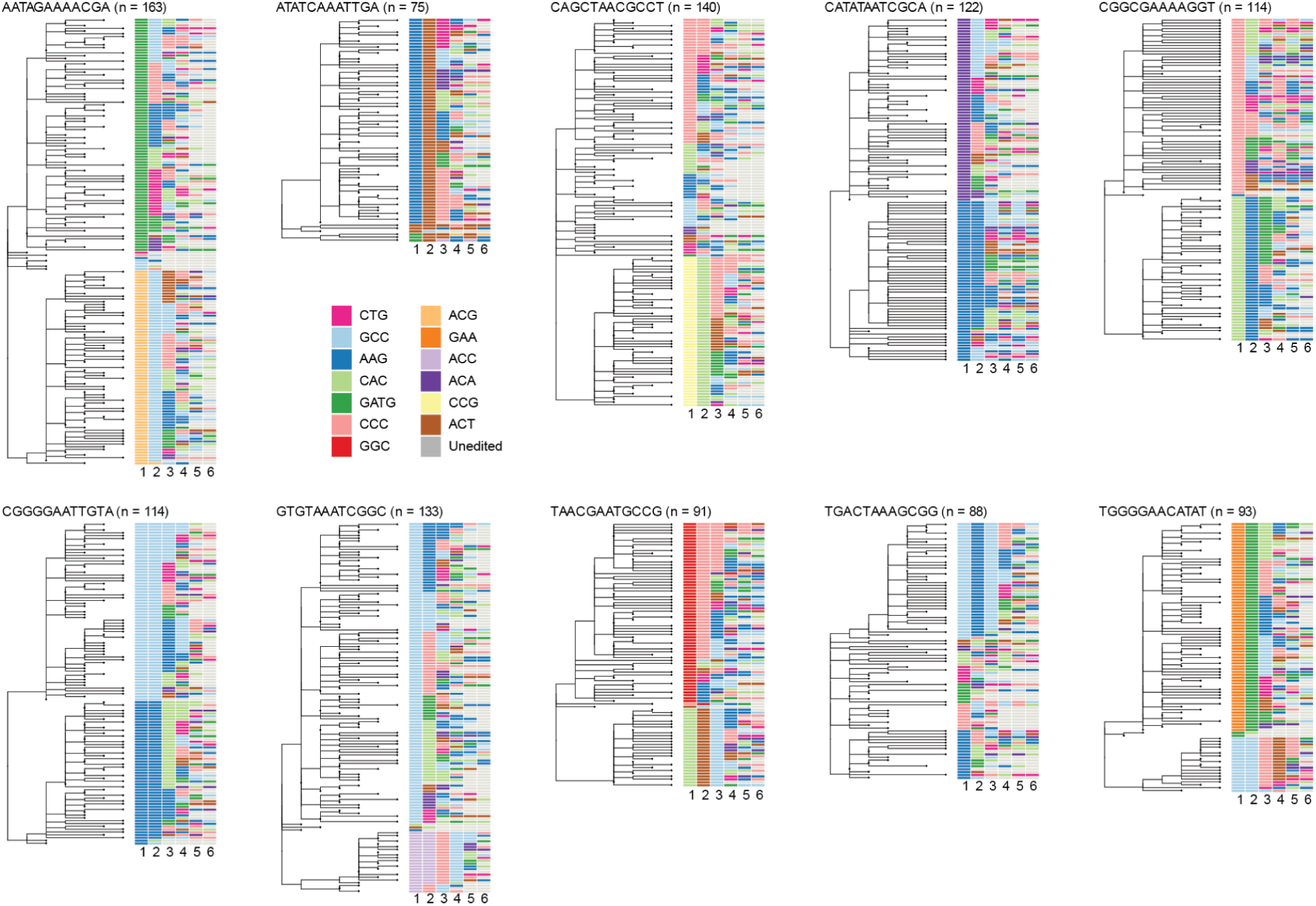
Neighbor-joining trees reconstructed from genotypes observed in bulk amplicon sequencing of tapes. As in Fig. 1F, but for the ten remaining tape-BC-defined integrations, with the corresponding edit patterns shown at right (colored by symbol; grey, unedited).

**Figure S4.**
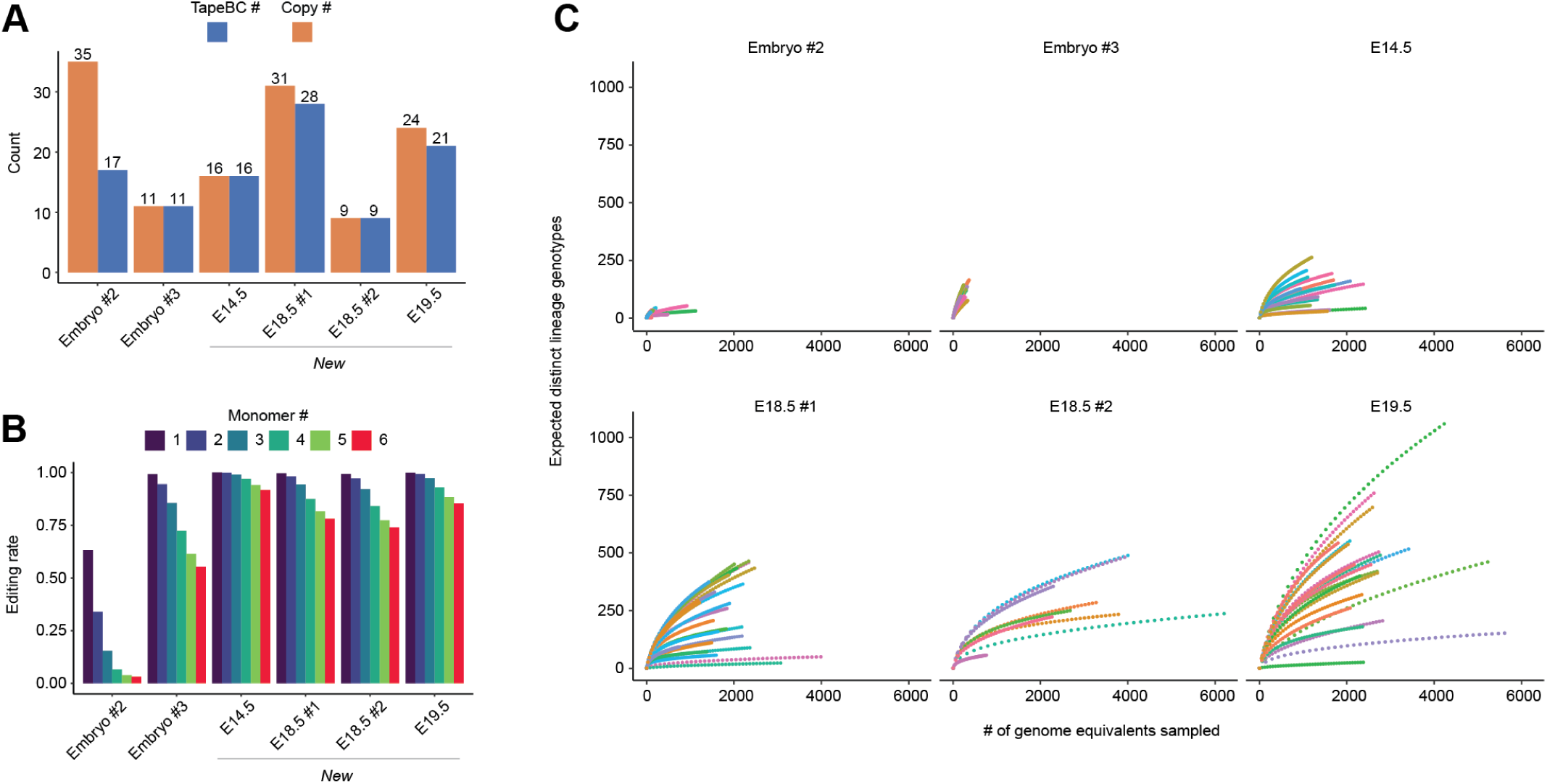
Robustness of recording in DNA Typewriter embryos from additional PNI sessions. **(A)** Number of distinct tape-BC sequences (unique integration barcodes; blue) and the estimated total number of genomic integrations after accounting for multi-copy barcodes (orange), shown separately for six embryos harboring tape integrations from the original PNI session, harvested at E13.5 (embryos #2 and #3), as well as four additional PNI sessions, harvested at E14.5, E18.5 (n = 2), and E19.5. **(B)** Per-monomer editing rates for each embryo, expressed as the fraction of resolved reads edited at each of the six monomers (colored 1-6), collapsed with respect to independent integrations. **(C)** Rarefaction analysis of recorded diversity in each of six embryos. For each tape-BC integration (colored by tape-BC), the expected number of distinct tape-level genotypes (y-axis) is plotted as a function of the estimated number of genome equivalents sampled (x-axis). For all four new embryos, the diversity observed for each tape-BC integrations lies on the steeply rising portion of its curve, with a terminal slope far from zero, indicating that recorded diversity is far from saturated and is limited by the material sampled rather than by recorder capacity. That the overall diversity captured in each the four new embryos exceeds even embryo #3 is attributable to our use of 50 ng rather than 5 ng of genomic DNA as input to the tape amplification reaction.

**Figure S5.**
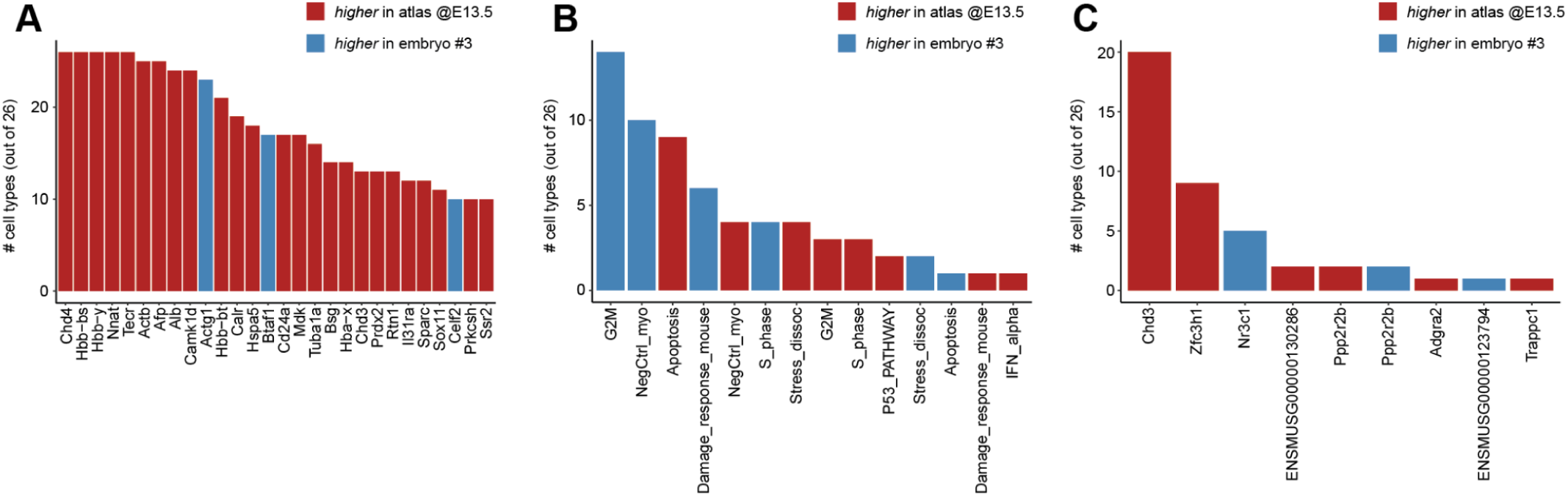
Cell-type-matched transcriptional comparison between embryo #3 and the atlas. **(A)** Recurrently differentially expressed genes between atlas and embryo #3, tallied across 26 shared cell types (each with ≥500 cells per dataset; 500 cells randomly sampled from each per cell type). Genes shown are significant (Bonferroni-corrected p-value < 0.05, Seurat FindMarkers) in ≥10 of 26 cell types. Bar height indicates the number of cell types in which the gene was differentially expressed; color indicates direction (red, higher in atlas; blue, higher in embryo #3). **(B)** Recurrently differential gene programs between atlas and embryo #3, tallied across the same 26 cell types. Each cell was scored per program using Seurat AddModuleScore, and programs were compared per cell type by Wilcoxon test (FDR < 0.05). Bar height indicates the number of cell types in which the program differed significantly; color indicates direction. NegCtrl_myo is a myogenesis negative control scored in non-muscle cell types. **(C)** Same analysis as in panel **A**, restricted to the 91 genes with a putative DNA Typewriter integration site within its gene body or within ±50 kb of its TSS (**table S1**). Bar height indicates the number of cell types in which the gene was differentially expressed (FDR < 0.05) and color indicates direction (red, higher in atlas; blue, higher in embryo #3). ENSMUSG00000130286 is proximal to the same integration site as *Chd3.* ENSMUSG00000123794 is proximal to the same integration site as *Nr3c1*.

**Figure S6.**
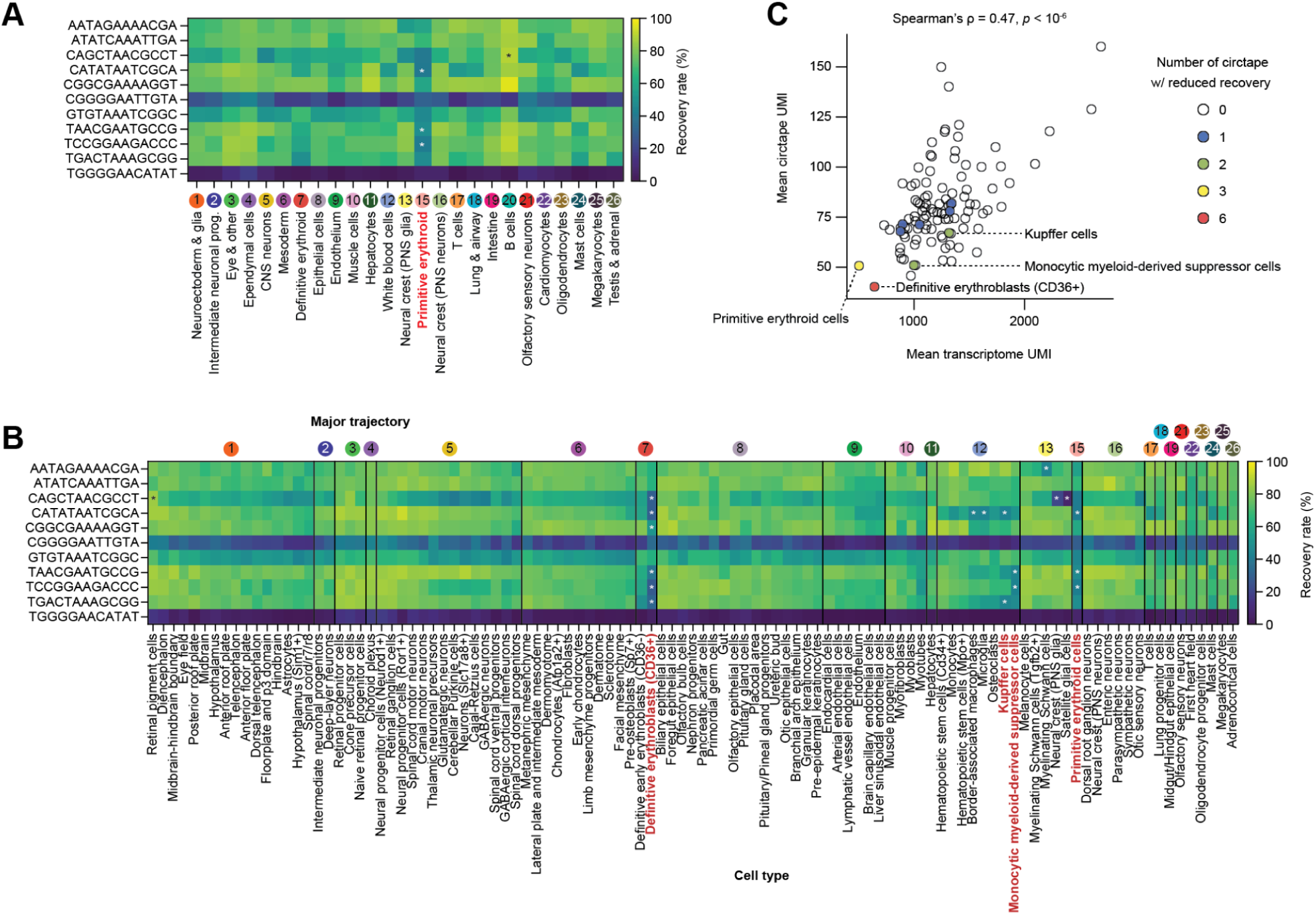
Recovery rate of circtape from 11 integrations by cell type. (A-B) 25 trajectories **(A)** and 105 cell types represented by ≥100 single cells **(B)** are included in this analysis. The recovery rate of nine tape-bc-defined circtapes (all but the two most poorly recovered) was compared to a label-permuted null (100 permutations; two-sided test). Tapes whose recovery rate was significantly enriched or depleted relative to the null are marked with black and white asterisks, respectively (Benjamini–Hochberg FDR < 0.05, ≥1.5-fold change). The labels of trajectories or cell types in which the recovery rates of at least two circtapes were significantly different are shown in red. **(C)** A scatter plot comparing the mean transcriptome UMI count and mean circtape UMI count for each cell type. Dot color represents the number of circtapes that showed reduced recovery under permutation in panel **B**.

**Figure S7.**
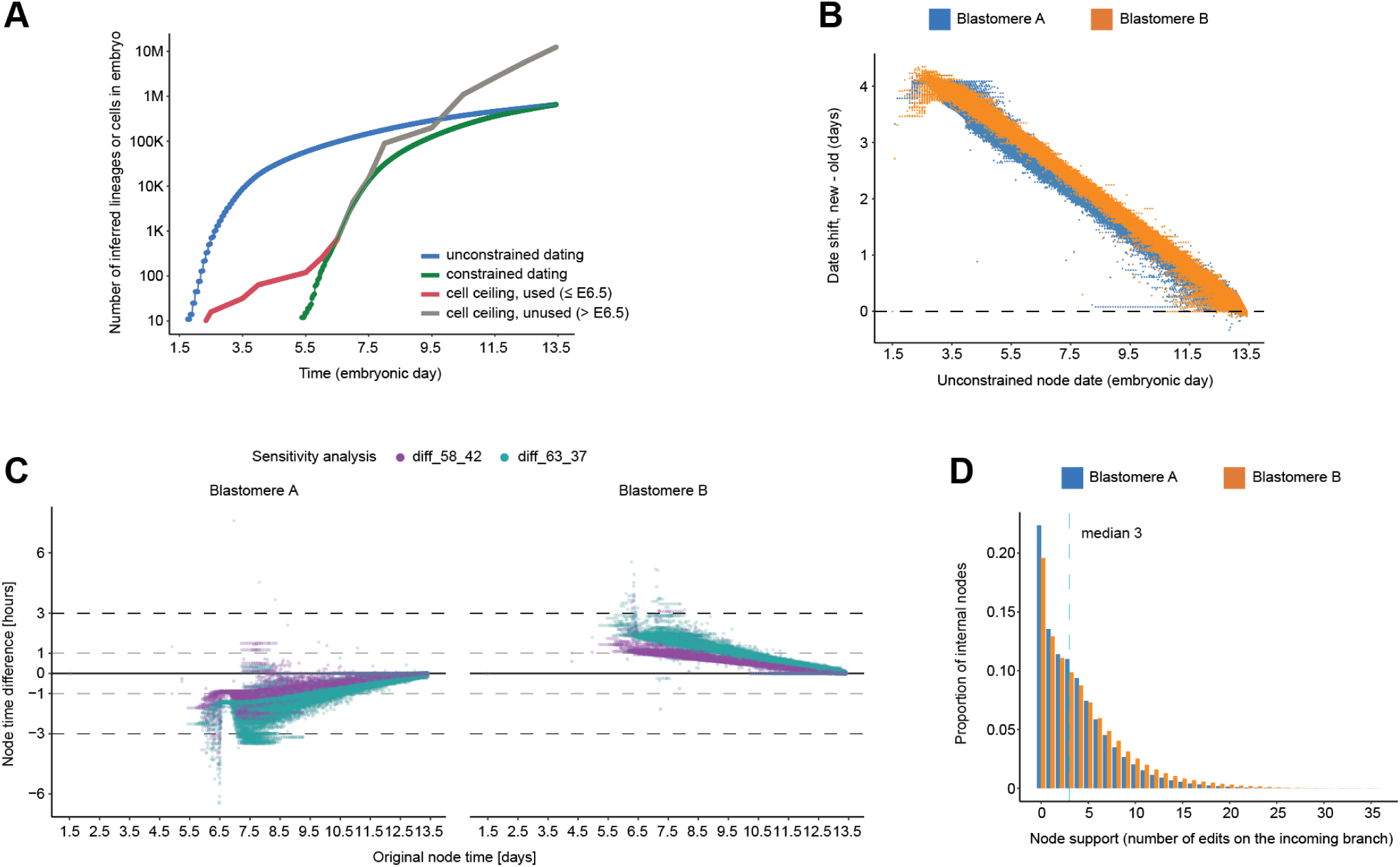
A cell-number ceiling corrects implausibly early node dates, and most internal branches carry multiple independent supporting edits. **(A)** Number of inferred lineages (blue, green) or anticipated number of cells in embryo (red, grey) over developmental time. Blue line, number of ancestral lineages for backbone tree obtained with unconstrained dating. Green line, number of ancestral lineages for backbone tree obtained with constrained dating (2-hour floor, cell number ceiling). Red/grey line, anticipated number of cells in embryo at various developmental times based on literature (**table S3**), colored differently for timepoints where the literature-based ceiling was constraining (red, ≤E6.5) vs. where it was not (grey, >E6.5). **(B)** Scatter plot of shifts in dating (y-axis, constrained minus unconstrained) vs. the unconstrained date (x-axis), for internal nodes of blastomere A (blue) and B (orange) subtrees. **(C)** Shift in node dates, shown separately for blastomere A and B lineages (facets), for the 58/42% (purple) and 63/37% (teal) ceiling splits, relative to the 50/50% ceiling split. Dashed lines mark ±1 and ±3 hour thresholds; most node dates fall within ±1-3 hours even under the more extreme 63/37% scenario. **(D)** For each internal node, support is quantified as the number of edits accumulated on the incoming branch. A normalized histogram is shown that includes the proportion of internal nodes with various levels of support for the A (blue) and B (orange) backbone trees; dashed lines indicate the median. 78% (A) and 80% (B) of internal branches are supported by ≥1 edit. 64% (A) and 68% (B) of internal branches are supported by ≥2 edits.

**Figure S8.**
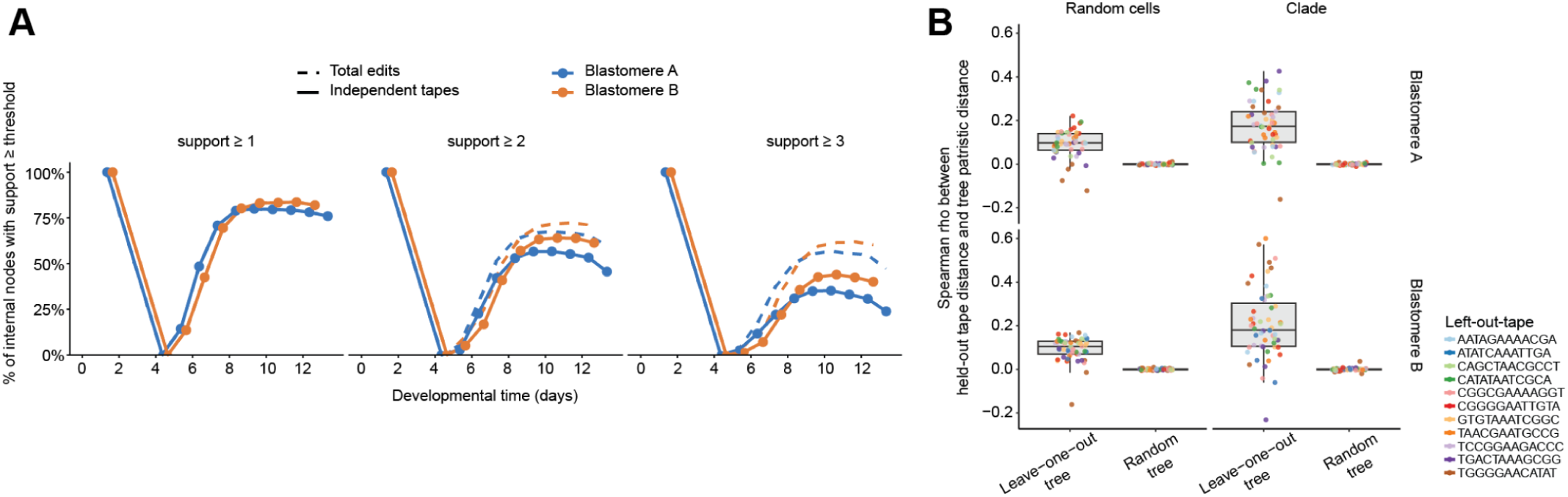
Reliability of backbone tree topology. **(A)** Node support based on total edits versus edits from different tape integrations. The fraction of internal branches meeting support thresholds of ≥1, ≥2, or ≥3 is shown over developmental time for blastomere A and B subtrees. Solid lines indicate support based on the number of edits from distinct tape integrations, whereas dashed lines show the number of total new edits supporting each branch. At the ≥1 threshold the two measures coincide by construction, while at ≥2 the fraction of supported branches reaches a peak of 67%/72% for total edits, falling to 57%/64% for the tape-integration-based measure; and at ≥3 the corresponding peak values are 57%/62% falling to 35%/44% (A/B). **(B)** Leave-one-tape-out validation of reconstructed tree. For each blastomere, we generated five replicate trees under two sampling schemes: 1,000 randomly sampled cells (“Random cells”) or a ∼1,000-cell clade from the backbone tree (“Clade”). For each cell sample, a tree was reconstructed leaving out one tape at a time, and we computed the Spearman’s ρ between the held-out tape’s pairwise cell distances and the tree’s patristic distances (“Leave-one-out tree”). As a negative control, the same correlation was computed after shuffling cell labels across tree nodes (“Random tree”). Points are colored by the identity of the left-out tape.

**Figure S9.**
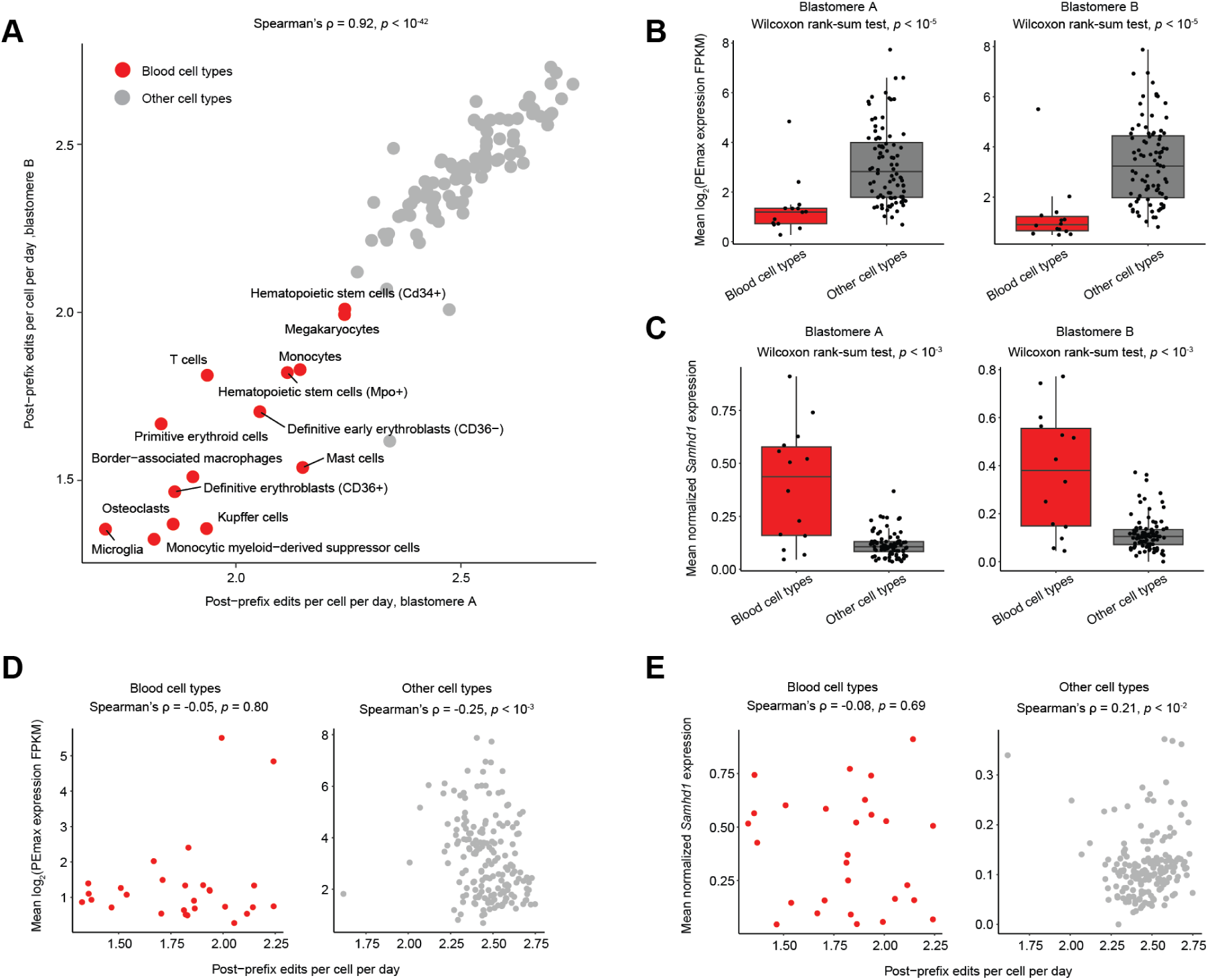
DNA Typewriter’s cumulative activity in embryo #3 was reduced in hematopoietic cell types. **(A)** Scatter plot comparing per-cell-type post-prefix editing rates between the two founding blastomeres (A and B) across 102 abundant cell types (≥0.01% frequency in both blastomeres). For each cell, the number of edited monomers accruing after the 2-cell-stage burst was counted for every recovered tape. These values were averaged across cells within each cell type to yield a per-tape editing rate for that cell type, divided by 13.5 to give a per-day rate, and then summed across tapes to give a per-cell, per-day editing rate for each cell type. Each dot represents one cell type; blood-associated cell types are shown in red and labeled, all others in grey. **(B)** *PEmax* transgene expression levels were inferred from the number of reads mapping to the model sequence of a PEmax-P2A-dTomato bicistronic construct, normalized by transcript length (8,005 bp) and by total reads per million to yield FPKM, then log_2_-transformed with a pseudocount of 1. For each cell type, the mean log_2_(FPKM + 1) was computed separately across cells assigned to blastomere A (left) and blastomere B (right). Cell types are grouped into blood-associated (red) and all others (grey). Two groups were compared with a two-sided Wilcoxon rank-sum test on per-cell-type means. Boxes show median and interquartile range with whiskers extending to 1.5× IQR. **(C)** For each cell type, *Samhd1* expression levels were computed from single-cell transcriptomes, log-normalized with Seurat (scale factor 10,000), and averaged across cells within each cell type, separately across cells assigned to blastomere A (left) and blastomere B (right). Cell types are grouped into blood-associated (red) and all others (grey). Two groups were compared with a two-sided Wilcoxon rank-sum test on per-cell-type means. Boxes show median and interquartile range with whiskers extending to 1.5× IQR. **(D)** Scatter plots of post-prefix editing rates for each cell type, plotted against their *PEmax* expression levels, stratified by color into blood-associated (red) and all other (grey) cell types. **(E)** Same as panel **D**, but plotting against cell types’ *Samhd1* expression levels.

**Figure S10.**
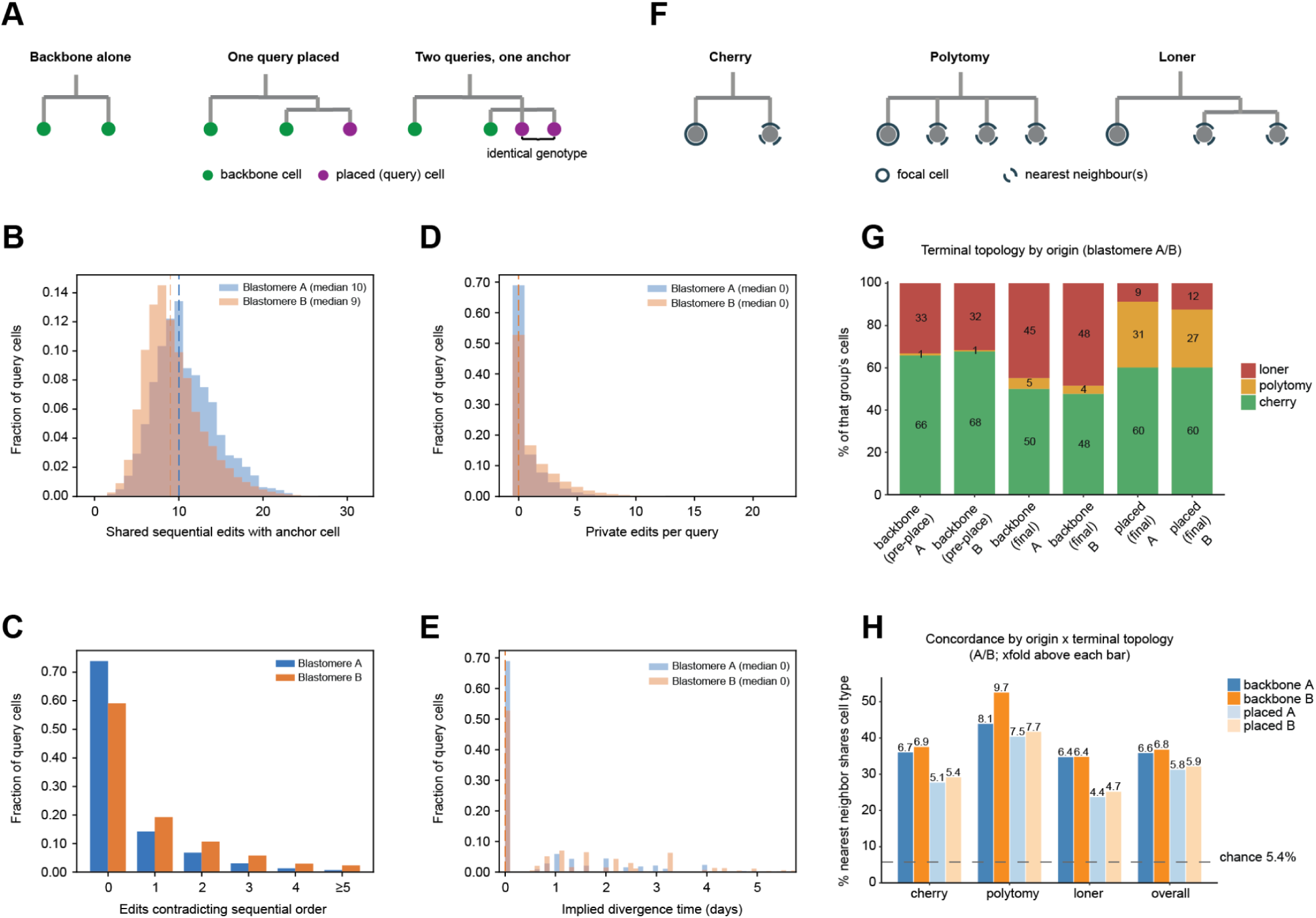
Query cells are placed at short edit distances from their anchors and recover cell-type concordance nearly as well as backbone cells. **(A)** Grafting query cells onto the backbone reshapes tree terminal topology. Backbone cells (green) were first placed by neighbor-joining; query cells (purple) are placed onto their nearest anchor by tape’s editing distance. Placing one query converts a backbone cherry into an anchor-query cherry, displacing the anchor’s former mate into a loner. Query cells with identical genotypes are represented as a single polytomy rather than being arranged into an arbitrary ladder. **(B)** Distribution of shared sequential edits between each query cell and its matched anchor, defined as the length of their common edited prefix summed over tapes genotyped in both cells. Founder-burst edits, which are shared by all cells within a blastomere, are excluded, so the values shown reflect only lineage information written after the first cleavage. **(C)** Fraction of placements, per blastomere, by the number of anchor edits that contradict the sequential edit order, *i.e.* an edit present in the anchor but absent, or differently written, in the query. Most placements are fully consistent, and placements with more than one or two contradicting edits are rare. **(D)** Number of private edits. Zero indicates that the query is genotype-identical to its anchor at every recovered monomer. **(E)** Implied divergence time between each query cell and its anchor, in days, obtained by scaling the pendant edit distance to the pair’s inferred common ancestor by the per-blastomere clock rate. Most query cells are placed as contemporaries of their anchor rather than at a resolved depth of their own. The discrete spacing of the non-zero bars reflects the integer edit distances from which these times are derived. **(F)** Terminal context of a cell and its equally related nearest neighbours, defined by local tree topology. Cherry: a leaf whose parent has exactly two tip children (one nearest neighbour). Polytomy: a leaf whose parent has ≥3 tip children, all counted as neighbours. Loner: a leaf whose only siblings are internal nodes rather than tips (contributes no sibling pairs). **(G)** Terminal topology by origin: cherry (exactly one leaf sibling), polytomy (≥2 leaf siblings), or loner (no leaf sibling; the parent’s other children are internal subtrees). Bars give the percentage of cells in each context for, left to right, the 655,701 backbone cells in the backbone tree alone (before placement) and in the final (after placement) tree, and the 625,440 placed cells in the final tree. Each group is shown separately for blastomeres A and B. Cherries fall from 67% of backbone cells before placement to 49% after, with loners rising from 33% to 46%; placed cells are 60% cherry, 29% polytomy and 10% loner, the excess polytomies arising because queries with identical genotypes are attached as a single polytomy rather than an arbitrary ladder. **(H)** For each cell, the fraction of its equally related nearest neighbors that share its cell type (for a cherry cell, its mate; for a polytomy or loner cell, the leaves tied for the fewest intervening branches). Each cell thus contributes a single value regardless of terminal context, making cherries, polytomies and loners directly comparable. Bars, mean across cells; dashed line, label-permutation null (5.4%, 100 permutations); value above each bar, fold-enrichment. Cell types were not used in placement.

**Figure S11.**
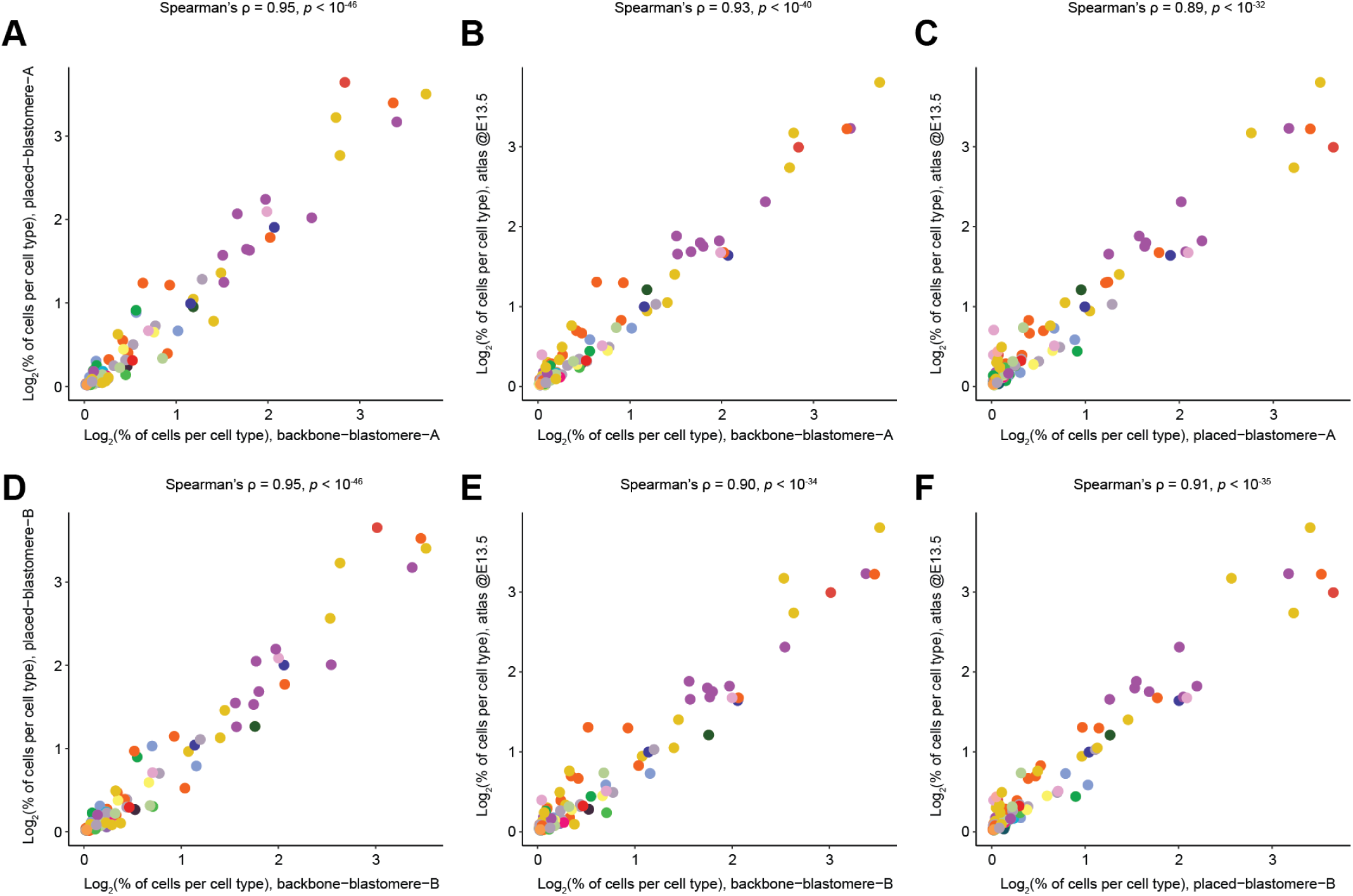
Cell type composition is concordant between backbone cells, placed cells, and the E13.5 embryo atlas. Each panel compares the relative abundance of individual cell types between two datasets, plotted as log_2_(% of cells per cell type + 1). Each point is one cell type, colored by major trajectory. Cell types representing <0.01% of cells in either dataset were excluded. Spearman’s ρ and the associated p value are shown above each panel. **(A)** backbone-blastomere-A versus placed-blastomere-A. **(B)** backbone-blastomere-A versus reference atlas at E13.5. **(C)** placed-blastomere-A versus reference atlas at E13.5. **(D-F)** As in panels **A-C**, but for blastomere B.

**Figure S12.**
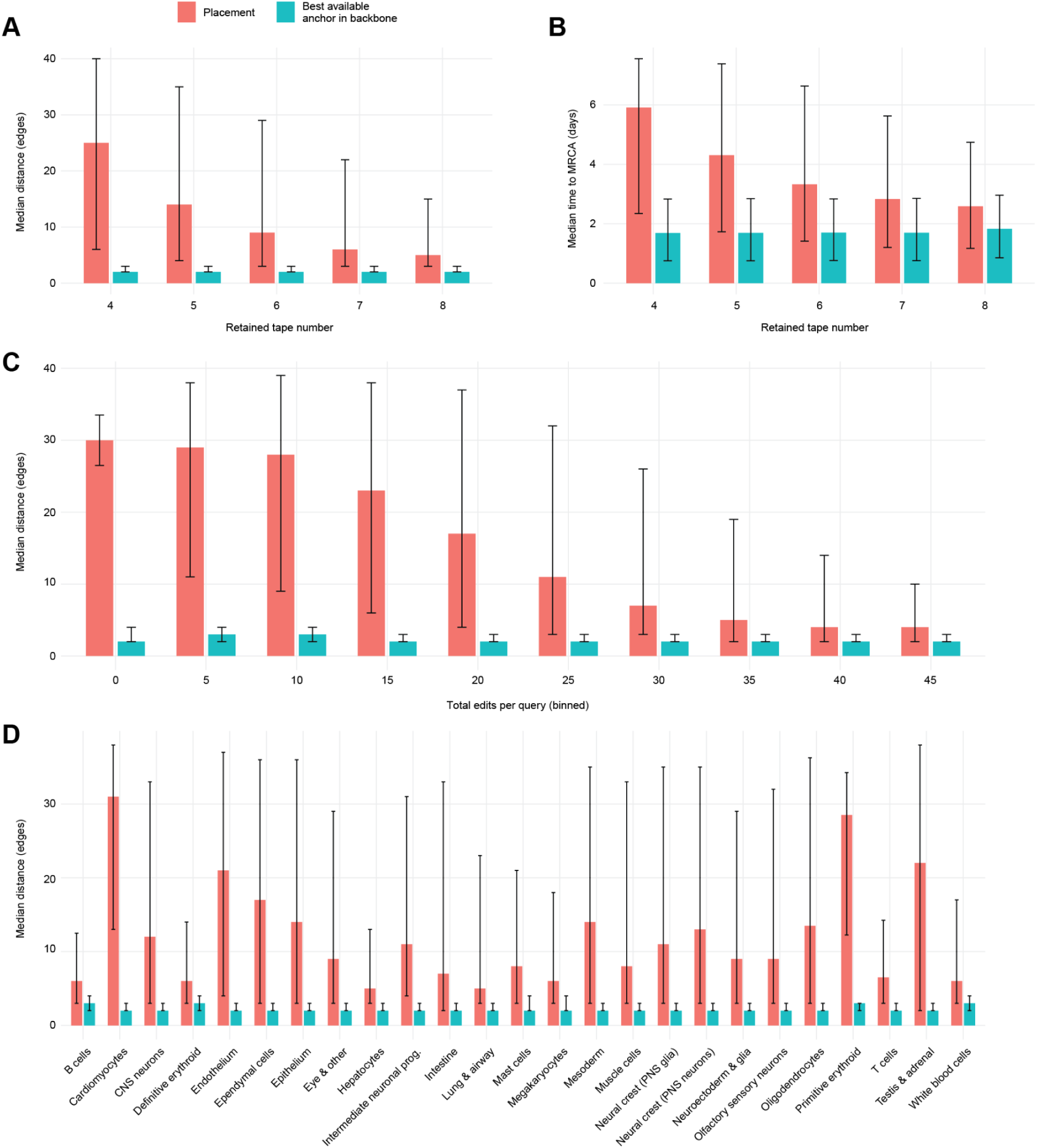
Accuracy of phylogenetic placement of held-out backbone cells. In each of five replicates, 50,000 tips per blastomere were pruned from the dated backbone tree (410,925 and 244,776 tips). Query genotypes were masked to 4-8 recovered tapes, with tapes removed in proportion to their empirical per-tape non-recovery rate across backbone tips, and then re-placed from the remaining tape genotypes (*n* = 500,000 re-placements altogether). Red, topological distance in the full tree between a cell’s true tip and the tip it was placed beside; teal, distance to its closest surviving relative in the pruned backbone, the best achievable target. **(A)** Distance vs. retained tape number. **(B)** Time to the most recent common ancestor vs. retained tape number. **(C)** Distance vs. total edited sites summed over retained tapes. **(D)** Distance by major trajectory. Bars, median; whiskers, interquartile range.

**Figure S13.**
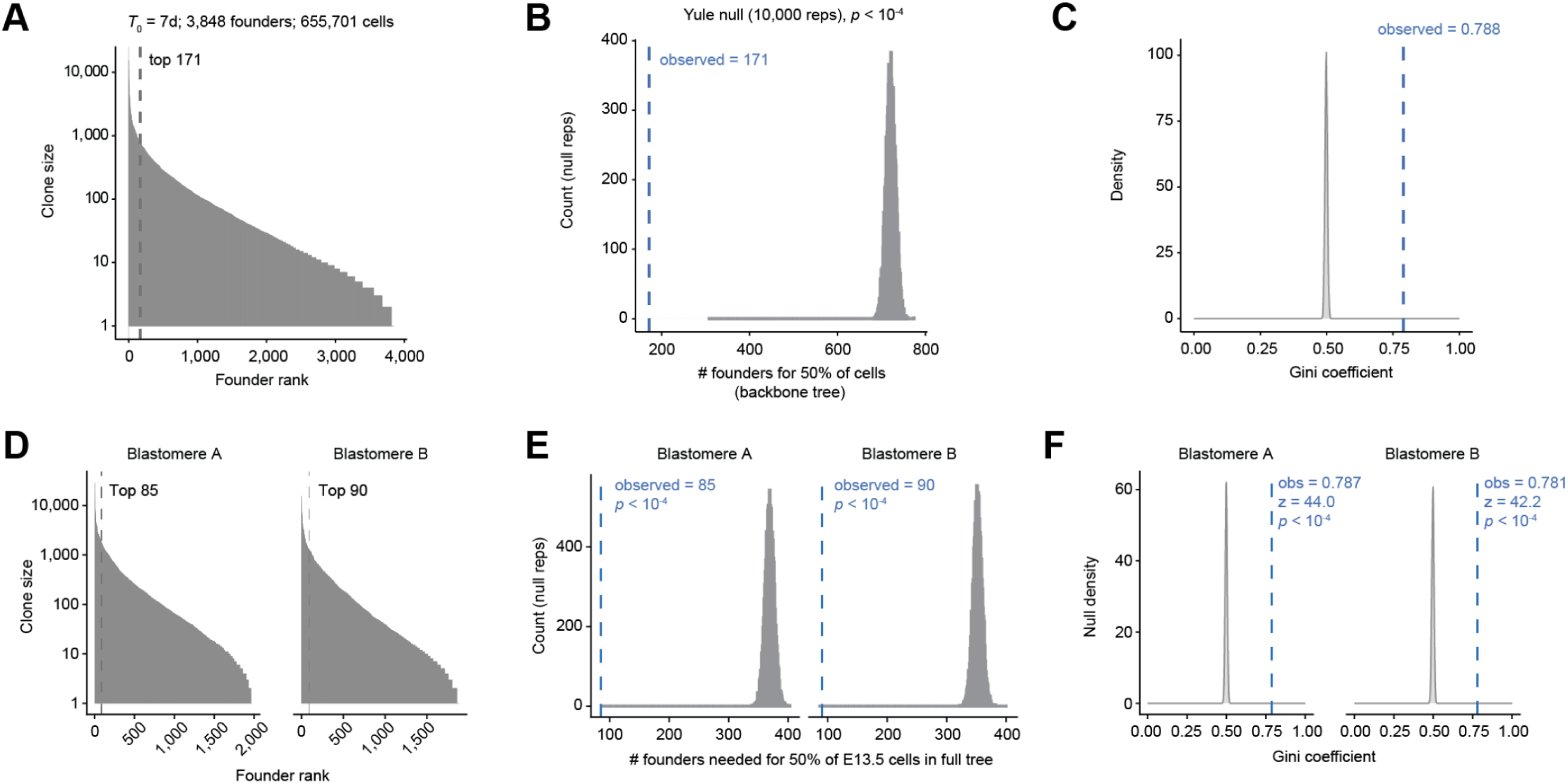
Patterns of clonal dominance are concordant between the full and backbone trees, and between the blastomere A and B subtrees. (A-C) As in Fig. 4A**–C**, computed on the backbone tree (*T*₀ = E7.0; 3,848 founders; 655,701 E13.5 cells). **(A)** Clone sizes ranked by founder; dashed line, minimum number of founders accounting for 50% of cells (171). **(B)** Yule null distribution of this number (10,000 replicates); dashed line, observed value; *p* < 10⁻⁴. **(C)** Yule null distribution of the Gini coefficient; dashed line, observed value (0.788); *p* < 10⁻⁴. (D-F) As in Fig. 4A–C, computed on the blastomere A and B subtrees of the full tree. **(D)** Rank-abundance of founder clone sizes (log_10_ scale) for blastomeres A (1,970 founder clones) and B (1,879 founder clones). Dashed lines mark the number of founders accounting for 50% of E13.5 cells in that subtree. **(E)** Number of founders required to generate 50% of all E13.5 cells, in observed data for blastomeres A and B (blue dashed lines) vs. the Yule null (grey, 10,000 replicates). *p* = *P* (null ≤ observed). **(F)** Gini coefficient of founder clone sizes, calculated separately for blastomeres A and B; the observed value (dashed line) is compared with its distribution under the Yule null (10,000 replicates). *p* = *P* (null Gini ≥ observed).

**Figure S14.**
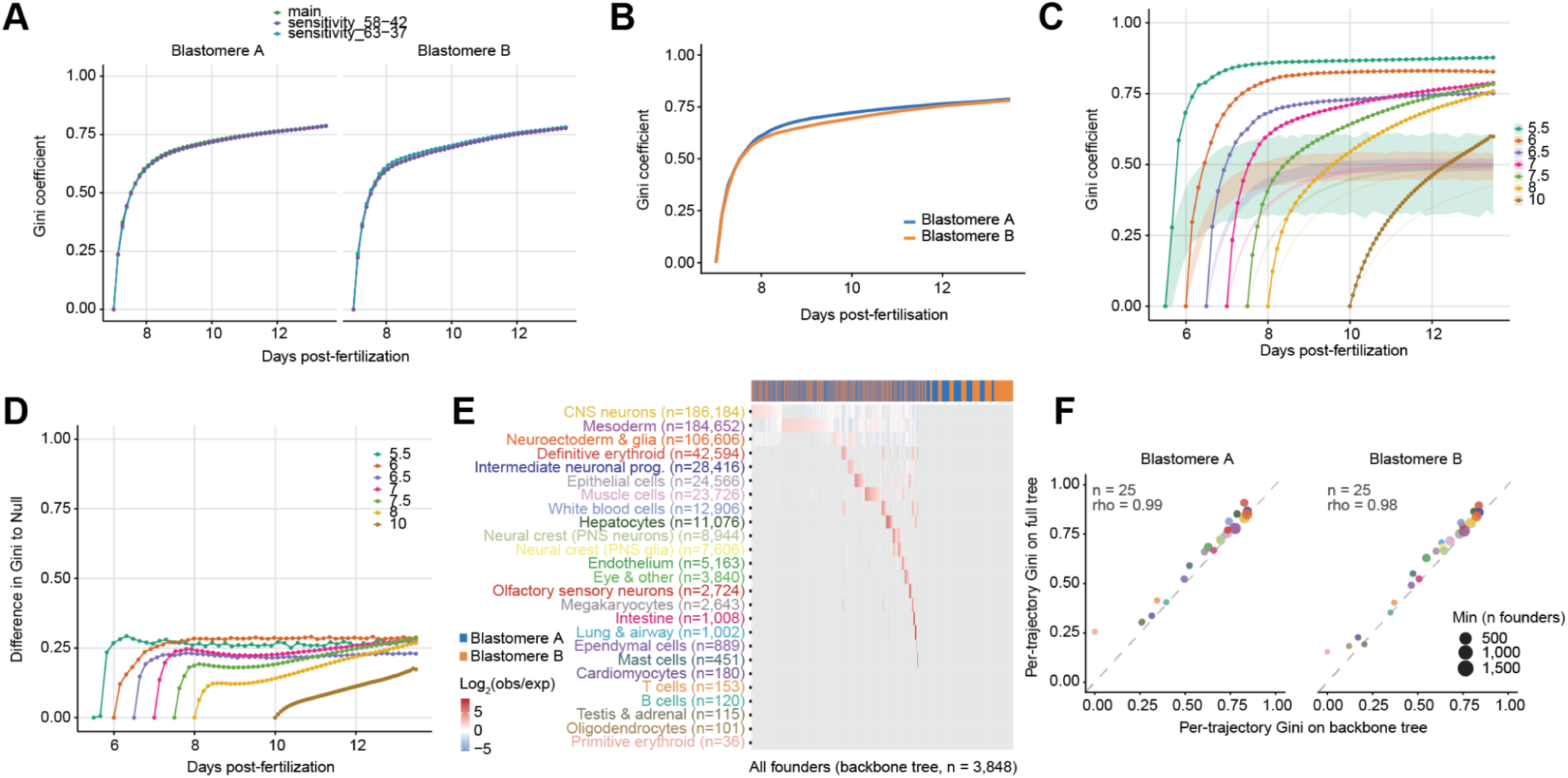
Robustness of clonal dominance to blastomere splits, to alternative *T*₀ values for defining founders, to breakdown by major trajectory, and to full vs. backbone tree. **(A)** Gini coefficient over time, shown separately for blastomere A and B subtrees of the backbone tree, with cell count ceiling split applied equally (50/50% ceiling split, green), as well as for trees dated with alternative cell count ceiling splits (58/42%, purple; 63/37%, teal), compared against the null expectation under a Yule null (grey band). All three dating scenarios show a highly similar rise in clonal dominance and plateau at comparable final Gini values, well above the null. **(B)** Emergence of dominance, calculated separately for founders derived from blastomeres A and B on the backbone tree. The Gini coefficient computed across founders’ descendant-lineage counts at each time *T*, tracking the fixed founder set forward from *T*₀ to E13.5 (blue or orange). **(C-D)** Gini coefficient over time **(C)** and difference between the Gini coefficient and the Yule-null expectation **(D),** for founders assigned at values of *T*₀ ranging from E5.5 to E10.0. Both panels are based on the backbone tree. Shaded bands in panel C show the corresponding null distributions. Earlier *T*₀ values yield a much steeper rise and higher final Gini coefficient, with the largest deviation from the null observed for *T*₀ between E5.5 and E7.5, consistent with the strongest gain in clonal dominance occurring during early gastrulation. **(E)** Heatmap showing the contribution of individual founder clones to each major trajectory, calculated on the backbone tree. See Fig. 4E for layout details. **(F)** Reproducibility of per-trajectory Gini indices at E13.5 compared between the full and backbone tree for blastomeres A and B, for 25 trajectories. Point size shows the smaller of the two founder counts (*i.e.* between blastomere A vs. B). Dashed line, *y* = *x*.

**Figure S15.**
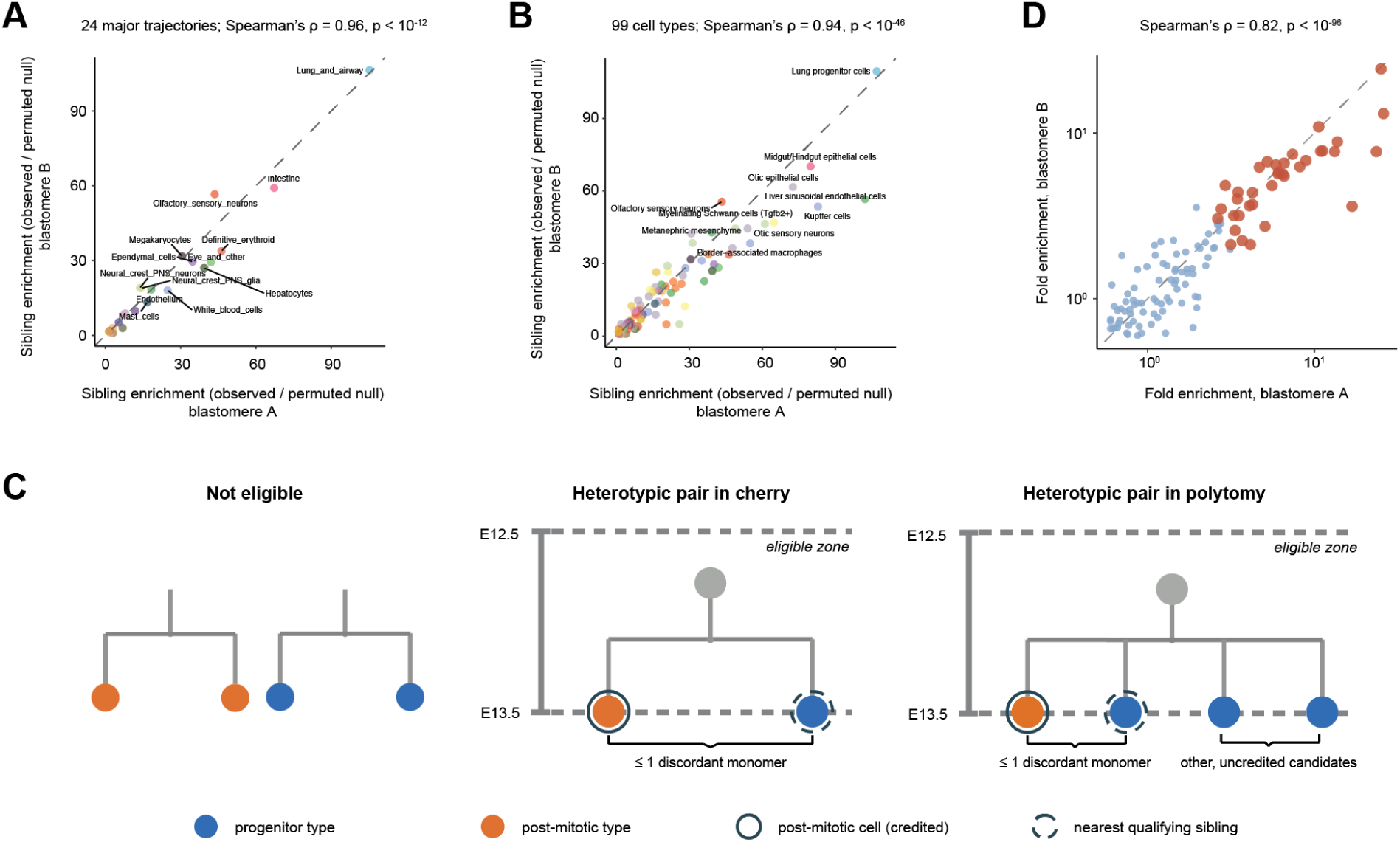
Tree siblings share annotations far in excess of chance, and reproducibly so. **(A)** Sibling enrichment for each of the 24 major trajectories, computed independently for descendants of blastomere A (x-axis) and blastomere B (y-axis). Trajectories were restricted to those present at ≥0.01% of cells in both the A and B subtrees. Enrichment is the ratio of same-trajectory sibling pairs to the mean count under a null that shuffles major trajectory labels across tips within A/B-defined half-embryo subtrees, holding sibling structure fixed (100 permutations), with a pseudocount of 1 in numerator and denominator to stabilize rare fates. Each point is one major trajectory, colored as in Fig. 2A; dashed line, *y = x*. Trajectories with the highest mean enrichments are labeled. Spearman’s ρ = 0.96, p < 10^-12^. Values for all trajectories are given in **table S4**. **(B)** As in panel **A**, but computed at cell-type resolution across 99 cell types present at ≥0.01% in both the A and B subtrees. Spearman’s ρ = 0.94, p < 10^-46^. Values for all cell types are given in **table S5**. The most enriched annotations are those committing from small, spatially restricted progenitor pools that then expand within an anatomical niche: *e.g.* lung and gut progenitors budding from the ventral endoderm (lung progenitor cells: 145-fold) (*102*, *103*), fetal liver-resident hematopoiesis (megakaryocytes: 35-fold) (*104*, *105*), and otic and dorsal root ganglia arising from placodal or neural-crest pools (otic sensory neurons: 79-fold) (*106*, *107*). **(C)** Sibling relationships at a shared ancestor, classified by post-mitotic status. Of 138 cell types, 27 were classified as post-mitotic, as listed in **table S6**. Not eligible: both members post-mitotic, or both members progenitor type. Heterotypic pair in cherry: one post-mitotic and one progenitor cell. Heterotypic pair in polytomy: an example in which a post-mitotic cell has more than one progenitor candidate, with the cell credited to the candidate with the fewest discordant tape monomers, with ties broken in favour of the candidate sharing the most called monomers. In both cases, the two cells should differ at no more than one discordant tape monomer, and their shared ancestor should be dated ≥E12.5. **(D)** Fold-enrichment of each progenitor→post-mitotic pairing computed separately for A and B, for the 389 of 640 evaluated pairings observed in both subtrees (log_10_-scale axes; dashed line, *y* = *x*). Red, 36 reliable progenitors → post-mitotic pairings that clear BH FDR < 1% and are ≥3-fold enriched in the overall tree (*i.e.* the subset shown in Fig. 5A; the 37th, retinal progenitor cells → retinal ganglion cells, is not observed in subtree B and so cannot be plotted here). Blue, the remaining 353 pairings. Spearman’s ρ = 0.82, p < 10^-96^, when calculated on all 389 pairings shown.

**Figure S16.**
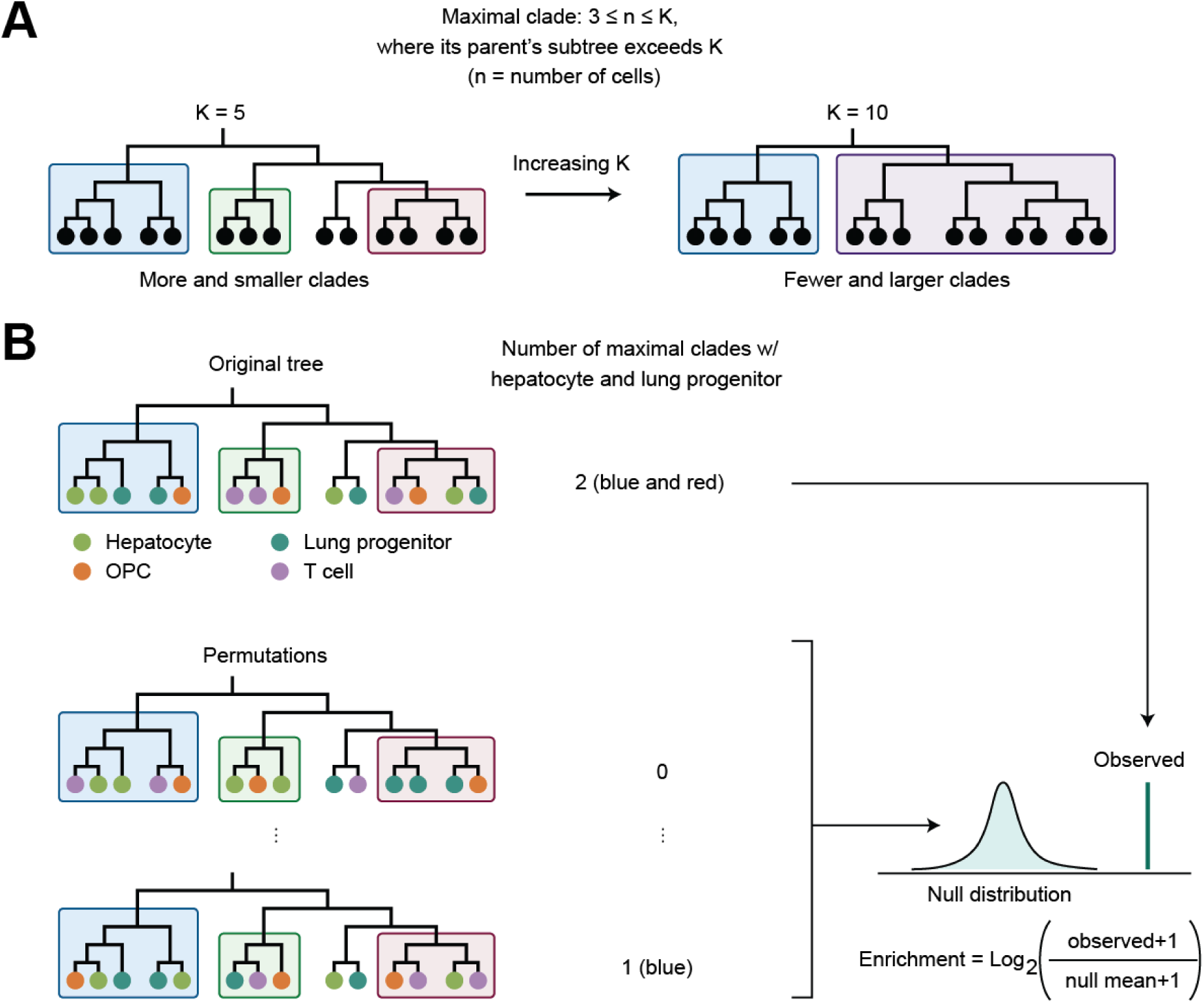
Schematic of maximal-clade analysis. **(A)** Definition of a maximal clade for a given size cap K. At each internal node of the lineage tree, the number of descendant cells in its subtree (n) is evaluated together with the number of descendant cells in its parent’s subtree. A node’s subtree is called a “maximal clade” when 3 ≤ n ≤ K while its parent’s subtree exceeds K. Colored boxes denote the maximal clades identified under two illustrative values of K (5 or 10) applied to the same underlying tree. Because clades can only merge with neighboring clades as K increases, a larger K yields fewer, larger clades. **(B)** Cell-type co-occurrence enrichment analysis, illustrated for the example pair hepatocyte and lung progenitor. The permutation-based enrichment procedure was used to test whether two cell types co-occur within maximal clades more than expected by chance. Top left, the original tree with tips colored by cell type and maximal clades outlined as in panel **A**. The number of maximal clades containing both cell types is counted (observed value; here, 2). Bottom left, the same tree topology and clade boundaries are held fixed while cell-type labels are randomly permuted across tips, and the co-occurrence count is recomputed for each permutation. Right, the resulting permutation values form a null distribution against which the observed co-occurrence count is compared. Enrichment for the pair is calculated as indicated, with a pseudocount added to stabilize pairs with few co-occurrences. This procedure is repeated for every pair of cell types.

**Figure S17.**
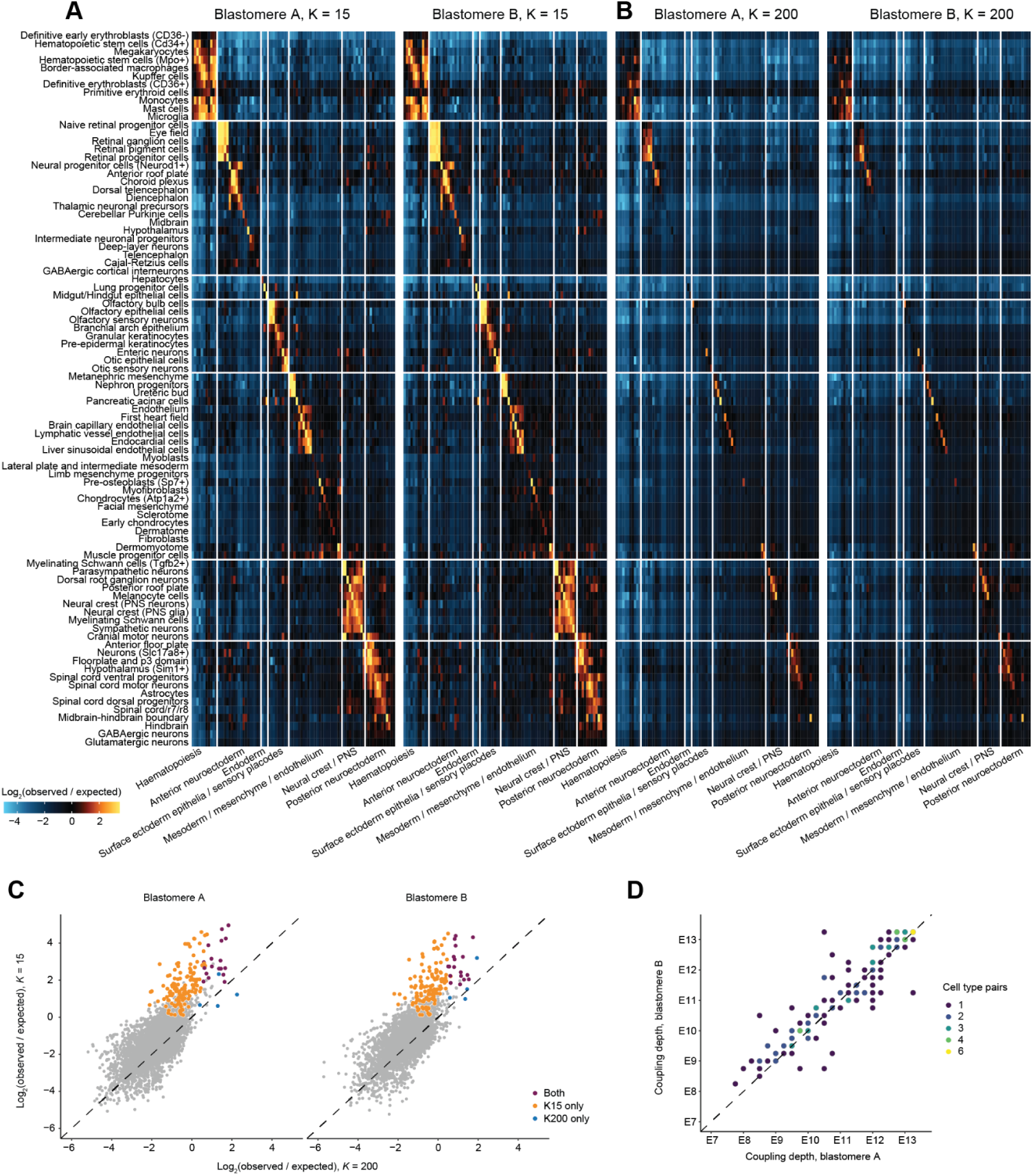
Clade co-occurrence analyses yield consistent patterns when conducted independently on subtrees defined by blastomeres A and B. **(A)** Timed fate couplings between cell types are reproducible between A and B subtrees at K=15. Details as in Fig. 5B, but here calculated on A and B subtrees independently. **(B)** Timed fate couplings between cell types are reproducible at K=200 as well, but show weaker block structure than K=15. Details as in Fig. 5B, but here calculated on the blastomere A and B subtrees independently and at K=200 instead of K=15. **(C)** Co-occurrence enrichment at recent vs. ancient clade scales is highly reproducible. Each point is one of the 3,828 pairs of the 88 cell types (≥100 cells in each blastomere subtree), plotted by its log_2_ enrichment at K=200 (*x*) against K=15 (*y*), with results based on blastomeres A and B’s subtrees shown as separate panels. Color indicates significance (enrichment of ≥3 Poisson standard deviations over the permutation expectation, on ≥5 co-occurring clades) at K=15 only, at K=200 only, or at both values of K; in every case significance is required in both blastomere subtrees. Dashed line, *y* = *x*. Most pairs that reach significance do so only at K=15 and lie well above the identity line, *i.e.* their co-occurrence is specific to recent clades. **(D)** For each of the 130 cell-type pairs whose coupling depth is estimable independently in each blastomere (129 of which were called jointly), coupling depth was re-estimated independently in blastomere A (x-axis) and B (y-axis). Because both axes are quantized to the 0.25-days, points are colored by the number of pairs falling on each grid position. Dashed line, *y* = *x*.

**Figure S18.**
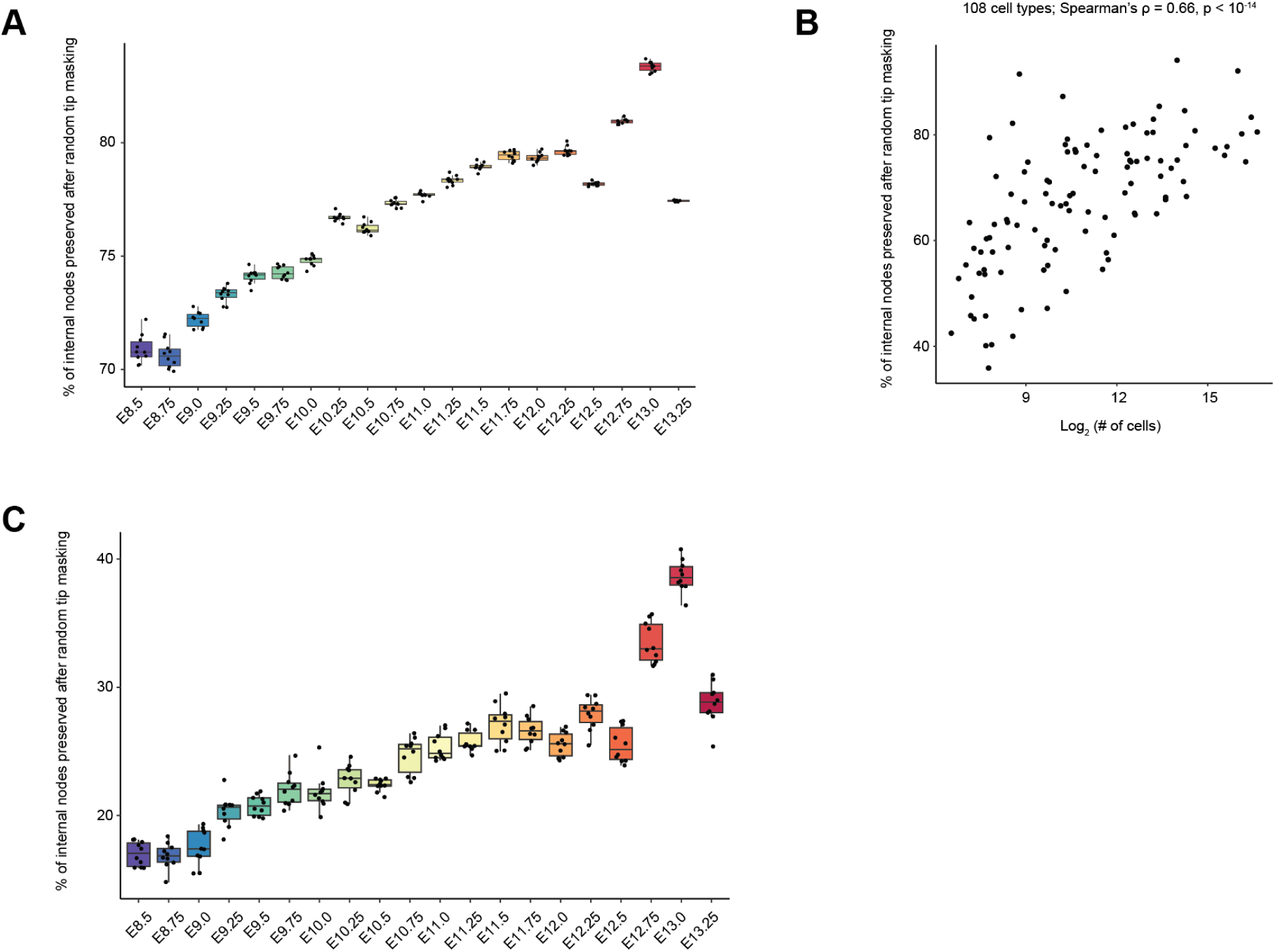
Benchmarking of imputed cell type labels of internal nodes, by random tip masking and by cousin-based prediction. A randomly selected 20% of E13.5 tips were masked, PCA coordinates of internal nodes were re-imputed by bottom-up averaging over unmasked descendants, and each internal node was reassigned a cell-type label via mutual k-nearest neighbors (K=200) to same-timepoint cells in PCA space. The reassigned labels were then compared to those imputed on the unmasked tree. This procedure was repeated 10 times with different random seeds. **(A)** Percentage of internal nodes retaining the same cell-type label (masked vs. unmasked) among the 926,852 successfully assigned internal nodes, stratified by embryonic day. Each dot represents one of 10 masking runs. **(B)** Across 108 abundant cell types (>0.01%), the percentage of internal nodes that retained the same cell-type label after masking of a randomly selected 20% of tips (y-axis; pooled across 10 masking runs) is compared to the log_2_-scaled number of internal nodes originally assigned to each cell type in the unmasked reference (x-axis). **(C)** For each timepoint, we randomly sampled 1,000 internal nodes as queries. For each query, we computed patristic (branch-length) distances to all other internal nodes from the same timepoint, excluded its ancestors and descendants, and averaged the imputed PCA coordinates of the K=10 nearest remaining nodes. Cell-type labels were then reassigned by majority vote over mutual nearest neighbors (K=200) to same-timepoint reference cells in PCA space (50 dimensions), following the same procedure as the original imputation pipeline. Performance was quantified as the percentage of query nodes whose cousin-based label matched the assignment from the original pipeline. Each dot represents one of 10 independent random query runs.

